# Large-scale comparative analysis reveals top graph signal processing features for subject identification

**DOI:** 10.1101/2025.08.20.671319

**Authors:** Thomas A. W. Bolton, Mikkel Schöttner, Jagruti Patel, Hugo Fluhr, Yasser Alemán-Gómez, Patric Hagmann

## Abstract

In magnetic resonance imaging, graph signal processing (GSP) is an analytical framework that enables to express regional functional activity time courses in terms of the underlying structural connectivity backbone. To this end, several parameters must be set during the processing of structural and functional data, and a variety of output features have been proposed. While emerging applications of the GSP framework have shown clear merits to reveal the neural underpinnings of brain disorders, behavioural facets or individuality, at present, the optimal parameter choices and feature types for an outcome of interest remain unknown. Here, we fill this gap by conducting a large-scale comparative analysis across parameter choices and candidate feature types. First, we show that all the studied factors of variation within the GSP pipeline significantly modulate feature vector patterns and feature coefficient values, evidencing the importance of an exhaustive characterization. Second, focusing on the ability to fingerprint individual subjects, we demonstrate that power spectral density and the structural decoupling index are the most all-around feature types, which harmoniously balance robustness to external sources of variation (head movement and acquisition settings), parsimony of the telling feature set, and generalization to altered parcellation specificities. Our results emphasize the importance, for future GSP studies, of carefully considering the undertaken structural connectivity and functional parameter choices as a function of the outcome measure of interest. More globally, they also highlight the relevance of large-scale comparative strategies in optimizing an analytical pipeline towards a specific goal. Our reported methodology can seamlessly be extended to other analytical approaches and outcome measures of interest, which we hope will be of use for future researchers in and outside the GSP subfield.

**Key points:**

- Graph signal processing (GSP) analysis involves many parameters to select and can yield diverse types of features.
- All investigated parameters significantly modulated feature vector patterns and values.
- Power spectral density and structural decoupling index are recommended for fingerprinting overall.

## Introduction

In magnetic resonance imaging (MRI) research, the past decade has consolidated network neuroscience approaches as an important part of the neuroimaging toolkit ^1,2^. Indeed, structural and functional networks (typically inferred, respectively, from diffusion MRI and functional MRI data) enable a versatile description of the brain, making it possible to investigate cross-regional interplays at various spatiotemporal scales ^3^. When a network is formed, graph theoretical metrics (see ^4^ for an overview) then offer a concise characterization of information flow. Such data have contributed to refine our knowledge of brain disorders ^5,6^, their commonalities ^7^, and, more globally, of human behavior ^8^.

A limitation of the above approaches, however, is their traditional reliance on an independent characterization of brain structure (reflected by a structural connectome, SC) and function (encoded in a functional connectome, FC), when in fact, the two are related: for example, in the mouse brain, hemodynamic changes induced by optogenetic stimulation were tied to feedforward connection strength and neuronal distributions, in a way that subsided with hierarchy^9^. The complexity of structure/function coupling is also acknowledged in human MRI, as it is impacted by development, aging, and disease ^10^, follows molecular, cytoarchitectonic, and functional hierarchies ^11^, and shows regional heterogeneity^11^.

Accordingly, many approaches have been devised to further characterize the links between brain structure and function. The conceptually simplest ones include the investigation of selected connections, correlating several metrics on both ends ^12^, a direct spatial correlation between SC and FC patterns ^13^, or the use of multivariate methods such as canonical correlation analysis and its variants ^14,15^ to extract patterns of correlation/covariance across structural and functional modalities ^16^. Inter-individual variability (*i*.*e*., how conform to the average a subject is regarding regional connectivity profiles) in structure and function have also been contrasted ^17,18^, and some have used multivariate regression ^19–21^, or deep learning^22^, to predict functional connectivity from a set of structural markers. Yet another family of approaches treat the structural network as a scaffold over which functional activity propagates, effectively offering a way to model structure and function at the whole-brain level ^23^. Such generative models enable to more directly tap into the mechanisms that underlie structure/function coupling, and offer the opportunity to study the impact of perturbations applied to the system. Prominent examples include differential equation-based modelling through sets of coupled oscillators ^24,25^, or a linear algebra-based representation with control theory ^26,27^.

A conceptually related method, which will be the one explored in the present work, is graph signal processing (GSP; see ^28–30^ for general reviews and ^31^ for a review of links with connectivity gradients). Note that we distinguish GSP from graph neural networks and graph learning, which will not be further touched upon therein; for a review of their applications on biological data, the reader is pointed to^32^. Conceptually speaking, GSP in neuroimaging involves three core elements: a connectome, which typically quantifies physical wiring across brain regions; a set of regional blood oxygenation level-dependent (BOLD) time courses, reflective of brain activity changes over time; and an operator (often a subtype of Laplacian), which enables the study of functional time courses in terms of their similarity to the underlying structure ^33–35^ (for a detailed mathematical description, see **Materials and Methods**).

An increasing amount of studies have been harvesting GSP to dissect the neural underpinnings of human behavior. For example, performance in a cognitive switching task was higher for subjects whose functional activity was more decoupled from the underlying brain structure ^36^, and a gradient of structure/function coupling from low-to high-level brain regions recapitulated behavioral domains ranging from sensorimotor functions to higher-order cognition^37^, enabling successful task decoding and fingerprinting in healthy subjects^38^. GSP also enabled to pinpoint novel markers of several brain disorders, such as autism spectrum disorder ^39,40^, 22q11 deletion syndrome ^41^, or internalized psychopathology^42^.

While the perspectives provided by GSP are promising, a currently unaddressed complicating factor is the impact of parameter choices made within a typical analysis pipeline: for example, different measures of connectivity can be considered in graph generation ^20,43^, with atlases of various scales ^44–46^, and many different approaches have been proposed to extract features reflective of structure/function coupling ^34,36,38–40,47^. So far, GSP studies have largely been conducted independently, on distinct sets of subjects, using different parameters settings, and with various end goals in mind. Thus, at present, an optimal methodology cannot be pinpointed.

To palliate this, a solution is the systematic comparison of possible methodological choices on the same high-quality benchmark dataset, so that confounding sources of variance are minimized ^48^. The collection and organization of sets of information has proved scientifically useful ^49^, and has been recently applied in functional neuroimaging to quantify the test-retest reliability^50^ and the cognitive relevance ^51^ of selected dynamic functional connectivity approaches, or the impact of preprocessing parameters on functional MRI data^52^. When it comes to GSP, such an endeavor is likely to both help interested newcomers steer their analyses towards the direction that best fits their goal, and foster the development of novel extensions that will fill existing gaps of performance.

In this work, we conducted a large-scale comparative analysis across a comprehensive set of candidate GSP parameter choices and output feature types. We exploited the high-quality dataset made available by the Human Connectome Project (HCP) ^53^, for which a collection of structural MRI, diffusion MRI and resting-state functional MRI data is available across more than 1000 healthy subjects. First, we studied the impacts of factors of variation within the GSP analytical pipeline on the patterns and the values of extracted feature vectors. We found that all the investigated factors play significant roles in shaping GSP features, including complex interactions and a diversity in terms of dominating contributions across feature types.

Second, we compared all candidate parameter choices and feature types in terms of a complementary set of quality criteria to pinpoint optimal choices. We quantified the robustness to unwanted sources of variance (acquisition settings and head movement), contrasted inter-individual variability to that between individual scans of the same subject, and examined sensitivity to differences in the used parcellation. In addition, our primary measure of interest was the ability to accurately fingerprint individual subjects from GSP features. The understanding of what distinguishes different individuals (*inter-individual variability*) has been the subject of many studies leveraging resting-state functional connectivity^10,54–60^, and gave rise to the now notorious concept of *connectome fingerprint* ^61–65^. This notion has been extended to subsequently derived network measures of interest^66^, to functional connectivity fluctuations over time ^67^, and to structure/function coupling ^38^. While uniqueness does not necessarily imply behavioral relevance^65,68^, the understanding of individual as opposed to group-wise effects remains essential towards the successful implementation of personalized medicine strategies ^69^. Furthermore, the contrast between intra- and inter-subject variability is also important to more clearly distinguish between state-like and trait-like features of brain function ^70^.

## Materials and Methods

### Subjects

For our analyses, we considered *S* = 100 subjects from the Human Connectome Project ^53^ (HCP) Young Adult dataset (28.32 *±* 3.53 years old, 52 males, 9 Hispanics, 87 Whites/9 Africans/4 Asians), for whom diffusion MRI and resting-state fMRI (four 15-minute scans) data were acquired. FMRI scans were acquired in pairs over two different days (short-handed REST_1_ and REST_2_, respectively): in each day, the first scan (referred to as SES_1_) was obtained with a left-right phase encoding direction, while the second (SES_2_) instead utilized a right-left phase encoding direction.

In order to limit the potentially deleterious impact of artifacts ^71,72^ as much as possible in our analyses, in addition to conducting state-of-the-art preprocessing (see below for details), subjects were selected to have less than 10% of frames scrubbed out at a threshold of 0.5 mm framewise displacement (FD) ^71^, and less than 0.2 mm average framewise displacement, across all four resting-state scans. For average FD, the resulting values in retained subjects were 0.13 *±* 0.02 (minimum 0.08, maximum 0.18) mm, 0.13 *±* 0.02 (0.08, 0.19) mm, 0.13 *±* 0.02 (0.07, 0.18) mm and 0.13 *±* 0.02 (0.09, 0.19) mm for REST_1_ SES_1_, REST_2_ SES_1_, REST_1_ SES_2_ and REST_2_ SES_2_, respectively. For the percentage of scrubbed frames, the resulting values were 0.34 *±* 0.68 (0, 4.86)%, 0.31 *±* 0.54 (0, 2.76)%, 0.29 *±* 0.52 (0, 2.43)% and 0.47 *±* 0.92 (0, 5.7)%.

### Magnetic resonance imaging data acquisition and processing

The imaging data considered in this study included the minimally preprocessed T1-weighted and diffusion-weighted MRI scans of the analyzed subjects, as well as their four resting-state scans (including preprocessing with ICA-FIX). We outline the appended preprocessing steps below for the generation of SCs and regional functional activity time courses.

### Generation of structural connectomes

Connectome Mapper 3 (CMP3) RC4^73,74^ was used to compute the SCs, taking T1-weighted structural and diffusion MRI data as input and calling routines from dedicated software packages at each step. For segmentation of the T1-weighted images, FreeSurfer^75^ recon-all version 6.0.1 was used, and parcellation was done with the Lausanne 2018 atlas ^45,76–79^, which is available at five different scales (respectively comprising 126, 172, 274, 504 and 1060 regions). We contrasted the results obtained at scales 1 and 3 (featuring numbers of regions consistent with the existing GSP literature, ranging from 90^47^ to 360^37–39^) to explore the impact of atlas granularity (**Figure 1A**). For scale 3, we also assessed the impact of discarding subcortical areas from the atlas (yielding a total of 216 remaining brain regions), as the neuroimaging GSP literature to date has remained mixed between adopting a purely cortical representation ^37–39^, or also including cerebellar and subcortical structures ^35,36,40–42,47^. In what follows, we will refer to scale 3 results as S3, scale 3 results without cerebellum/subcortex as S3^-^ and scale 1 results as S1.

**Figure 1:**
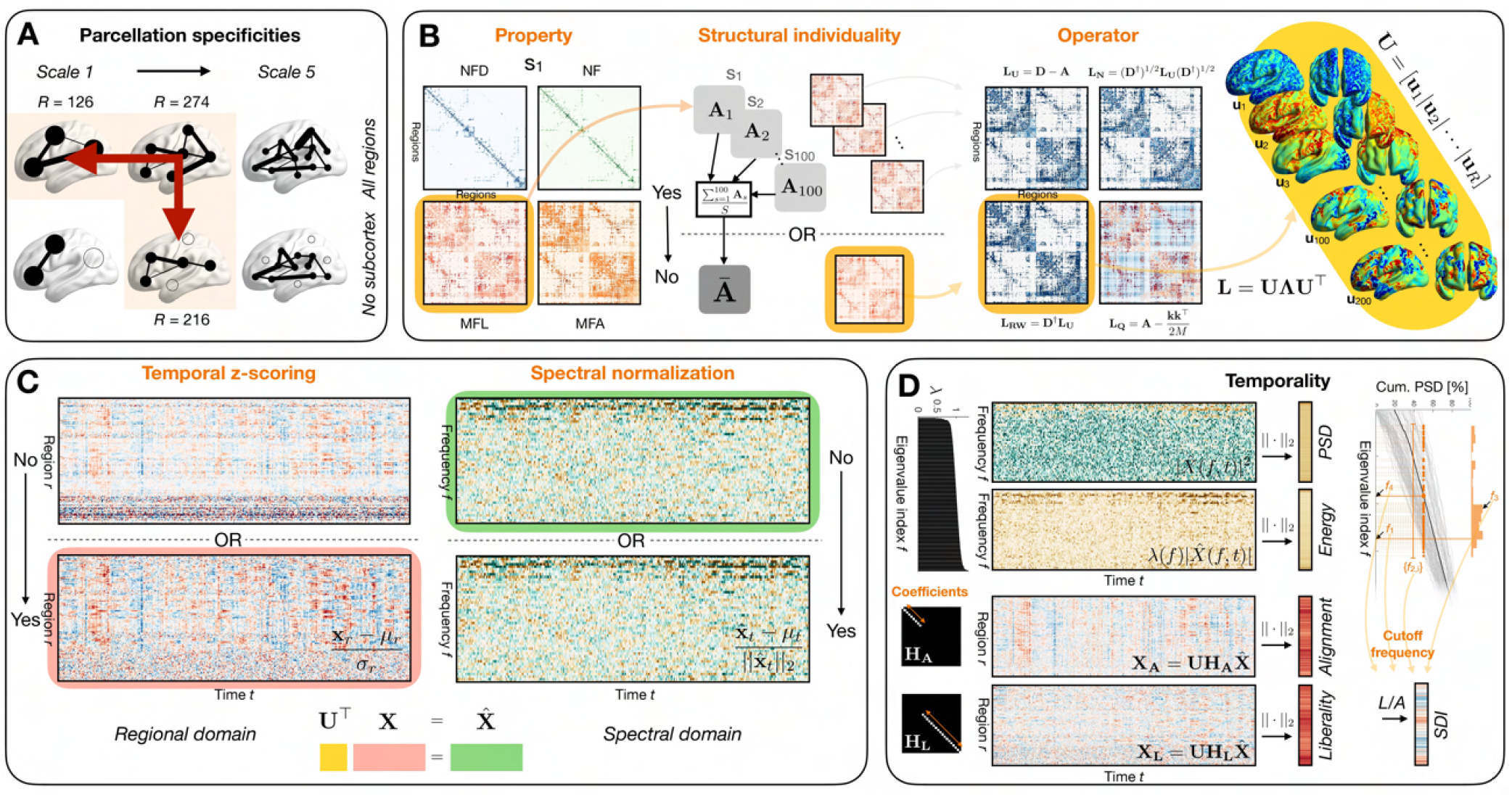
Main steps of graph signal processing analysis. **(A)** We considered a parcellation of the brain into *R* = 274 regions of interest (Lausanne 2018 atlas ^45,76–79^ at scale 3), and compared it to the use of an atlas with fewer regions of interest (*R* = 126, scale 1), or of the scale 3 atlas without subcortical and cerebellar structures (*R* = 216). By this mean, generalization capability across parcellation specificities can be assessed. **(B)** At the structural level, we probed the impacts of three factors of variation (highlighted in orange): the property used to construct an SC, structural individuality (*i*.*e*., use of an average SC as opposed to subject-specific SCs), and the operator used to extract the eigenmodes. **(C)** At the functional level, we addressed the impacts of temporal z-scoring, and of normalization of spectral coefficients at each time point. **(D)** We derived five types of features for each parameter combination: in the spectral domain (top half of the panel), we considered the power spectral density (PSD) and the energy. In the regional domain (bottom half), we studied alignment and liberality, which depend on the retained number of spectral coefficients upon spectral filtering (bottom left illustrations), and the structural decoupling index (SDI). This last measure depends on a cutoff frequency that partitions low- and high-frequency information, which can be inferred in various ways from the cumulative PSD distribution (top right illustration).

The diffusion MRI data were used to generate tractograms through MRtrix^80^, with the following parameters: deterministic tractography, white matter-seeded, constrained spherical deconvolution of order 8 with 10 million output streamlines. Then, the parcellated structural images and tractograms were combined to generate SCs for four different types of edge weights: normalized fiber density (NFD), number of fibers (NF), median fiber length (MFL) and median fractional anisotropy (MFA).

### Extraction of regional activity time courses

For processing of the resting-state functional data, a combination between FSL^81^ routines and in-house Python scripts (version 3.10.9) was used. For each subject, atlas volumes at scales 1 and 3 were warped (FSL’s applywarp command, nearest neighbour interpolation) from diffusion space to Montreal Neurological Institute (MNI) space. Then, nilearn’s clean img function was used for voxel-wise detrending, high-pass filtering (at 0.01 Hz), and confounds’ removal (6 head motion parameters and their derivatives, as well as the average white matter and cerebrospinal fluid signals, all of which were downloaded from the HCP beforehand). Importantly, clean img ensures that the removal of confounds is performed orthogonally from filtering, which avoids reintroducing artifacts to the data ^82^. Finally, averaging into regions of interest was performed with nilearn’s NiftiLabelsMasker.fit transform function. Output time courses were generated both with and without eventual temporal z-scoring.

### Graph signal processing overview and factors of variation

The main steps of GSP analysis, the explored factors of variation and the features under examination are summarized in **Figure 1B-D**. First, a structural connectome (SC), which recapitulates the physical wiring across brain regions, must be constructed. Mathematically, it takes the form of a matrix **A** ∈ ℝ^*R×R*^, with *R* the number of brain regions at hand and *A*_*i,j*_ the edge weight between regions *i* and *j*. Equivalently, one can represent this information as a graph 𝒢 with |𝒱 | = *R* nodes and |ℰ| ≤ *R*(*R* − 1)/2 edges.

Regarding SC generation, we considered two factors of variation (**Figure 1B**, left and middle left panels):

1. The type of information encoded by an edge *A*_*i,j*_ (which we refer to as *Property* hereafter), as recent research has shown that structure-function relationships significantly differ as a function of the chosen structural markers ^20^. We considered *normalized fiber density, number of fibers, median fiber length* and *median fractional anisotropy*.
2. The use of a group-averaged structural connectome (that is, the same underlying brain structure is hypothesized for all analyzed subjects) as opposed to subject-specific structural connectomes; we refer to this choice as *Structural individuality* in what follows. In the latter case, we have one SC **A**^(*s*)^, *s* = 1, …, *S* per subject *s, ergo*, each individual is characterized by a different structural connectivity backbone. While the former approach is likely over-simplistic, given that brain structure is itself behaviorally relevant^83^, it is also conceptually simpler, because functional activity is then decomposed as a function of the same mathematical basis across subjects. To date, the large majority of GSP studies have used the former approach ^34,35,37,38,40^, but see ^41^ for an application of the latter. In the remainder of this section, for simplicity, we will treat structural connectivity as a group-averaged matrix, but unless otherwise indicated, similar computational steps apply when using subject-specific SCs instead.

Following SC generation, a graph shift operator can be extracted. We considered the subtype of operator (*Operator*) as a factor of variation (**Figure 1B**, middle right panel), contrasting the following options:

1. The standard Laplacian **L**_**U**_ = **D** − **A**, where **D** is the diagonal matrix of regional degrees.
2. The normalized Laplacian **L**_**N**_ = (**D**^*†*^)^1/2^**L**(**D**^*†*^)^1/2^, with **D**^*†*^ the Moore-Penrose inverse of **D**.
3. The random-walk Laplacian **L**_**RW**_ = **D**^*†*^**L**.
4. The modularity matrix 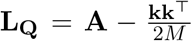, where 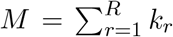 is the total edge weight (*k*_*r*_ is the degree for node *r*), as it has recently been suggested as a novel operator potentially more suitable to neuroimaging analyses ^84^. Notice the presence of a second term reflecting the probability of an edge in a degree-matched null model.

The eigendecomposition of the operator yields *connectome harmonics*, or *eigenmodes* (**Figure 1B**, right panel); taking the standard Laplacian as an example, we have **L**_**U**_ = **UΛU**^⊤^. Each harmonic (each column in **U**) showcases signal oscillations across the brain at a characteristic frequency (**Λ**_*k,k*_, with *k* the index of the eigenmode at hand). For the modularity matrix, instead of frequency, *modularity* is quantified by the eigenvalues. Throughout this work, we always assume sorted eigenvalues in ascending order. Interestingly, the harmonics with the smallest frequencies typically contrast large-scale brain attributes, such as both hemispheres or anterior *versus* posterior regions; consequently, resting-state networks can be well approximated by a linear combination of only a few ^33^, and functional connectivity can also be predicted from a limited subset of eigenmodes ^85^. This makes connectome harmonics a well-suited basis to characterize functional signals. As noted earlier, in the case of subject-specific SCs, one would obtain one set of harmonics and eigenvalues per subject: {**U**^(*s*)^, **Λ**^(*s*)^}, *s* = 1, …, *S*. This amounts to interpreting functional signals with regard to a different structure-encoding basis for each subject at hand.

Let us denote the regional time courses of activity for subject *s* by **X**^(*s*)^ ∈ ℝ^*R×T*^, with *T* the number of time points available. As a factor of variation in functional data, we examined the impact of *Temporal z-scoring* (**Figure 1C**, left panel), which scales all regional time courses so that they have a mean of 0 and a standard deviation of 1. Individual frames of activity 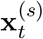 (that is, columns of **X**^(*s*)^) can then be projected onto the basis of eigenmodes (the so called *Graph Fourier transform* ^28^), which yields a characterization as spectral coefficients: 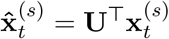 for all *s* = 1, …, *S, t* = 1, …, *T*. At this stage, as another factor of variation, one can decide to normalize the energy of the data in the spectral domain^40^ (*Spectral normalization*), so that it remains equal throughout time (**Figure 1C**, right panel).

From there, several strategies have been proposed to extract features reflective of structure/function coupling (**Figure 1D**), and are compared therein:

1. The computation of *power spectral density* (PSD) 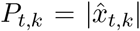 ^2 34^, where *k* denotes the *k*^th^ harmonic, which is larger if a given harmonic more strongly underlies the regional signals at time *t*.
2. The associated computation of *energy* 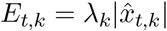, in which power for a given harmonic is weighted by the associated eigenvalue. Power spectral density and energy are both *spectral domain metrics*.
3. The computation of regional *alignment* and *liberality* ; for this purpose, filtering is first performed in the spectral domain. Let a diagonal matrix **H** ∈ ℝ^*R×R*^ with 1/0 elements for the frequencies to keep/discard, one can obtain filtered regional time courses as **X**_**F**_ = **UHU**^⊤^**X** ∈ ℝ^*R×T*^. Alignment denotes the component of the functional time courses in line with the underlying structure (*i*.*e*., low-pass filtering), while liberality reflects decoupled activity from the underlying structure (that is, high-pass filtering). A factor of variation specific to alignment and liberality is the number of harmonics *n*_*H*_ to retain for filtering (*Coefficients*). Here, we compared values ranging from *n*_*H*_ = 5 to 20% of *R* in 10 uniform steps. Past works have used *n*_*H*_ = 10 while ensuring robustness for other neighbouring values ^36,42^. Alignment and liberality are examples of *regional domain metrics*.

In all the above cases, one ends up with a representation of the data across both frequencies/regions and time (*i*.*e*., an *R × T* matrix). In order to simplify this into a feature vector, the L2-norm was computed over time. Following this step, a final feature type can be extracted: the *structural decoupling index* (SDI) ^37^, which directly contrasts the extent of alignment and liberality in each region. Let ||**X**_**L**_||_·,2_ and ||**X**_**A**_||_·,2_ the L2-norms of the high-pass filtered and low-pass filtered signals across time, respectively, then the SDI is given by 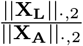.

Importantly, the SDI requires each spectral frequency to be included in either the aligned or the liberal signal contribution. To partition the eigenmodes between these two separate low-frequency and high-frequency sets, an SDI-specific factor of variation is then the selection of the cutoff frequency (*Cutoff frequency* ; **Figure 1D**, right column), which we define as the highest frequency index included in the low frequency set. We compared four candidate approaches to extract the cutoff frequency: first, a half-half split of the eigenmodes (that is, 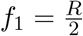), which does not utilize any information other than *R*. For the three other approaches, we considered splits that aimed at equating the energy of the regional signals within the low- and high-frequency sets ^37,38^, based on the cumulative PSD distribution which we will denote as ℙ_*t*_(*f*) for the signals at time *t*, with *f* = 1, …, *R* and ℙ_*t*_(*R*) = 1. In the second approach, we computed *f*_2,*t*_ so that ℙ_*t*_(*f*_2,*t*_) *<* 0.5 and ℙ_*t*_(*f*_2,*t*_ + 1) *>* 0.5 for all *t* = 1, …, *T*, yielding one cutoff frequency per time point: {*f*_2,*t*_}, *t* = 1, …, *T*. In the third approach, we computed the mode across individual cutoff frequency values: *f*_3_ = mode({*f*_2,*t*_}). In the fourth approach, we equated energy on the average cumulative PSD distribution across time points 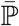, so that 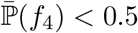 and 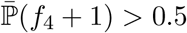.

### Impacts of factors of variation

We assessed to what extent the considered factors of variation would impact GSP features. We considered two complementary characterizations: (1) an analysis in terms of similarity in the overall feature vector pattern, and (2) another directly quantifying the changes in values of individual coefficients.

### Pattern similarity

We sought to project the high-dimensional feature vector information into a summarizing two-dimensional space in which closer data points exhibit a more similar feature vector pattern. To do so, we resorted to spectral clustering ^86^, following the same strategy as in ^87^. Briefly, for each feature type, subject *s* and scan *a*, we computed a similarity matrix **S**_*s,a*_ of size *n*_*p*_ *× n*_*p*_, with *n*_*p*_ the number of parameter combinations at hand (*n*_*p*_ = 128 for PSD and energy, *n*_*p*_ = 1280 for alignment and liberality, *n*_*p*_ = 512 for SDI). Each entry of a similarity matrix took the form [**S**_*s,a*_]_*i,j*_ = 0 if *i* = *j*, and 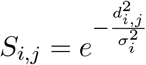 otherwise, where *d*_*i,j*_ is the cosine distance between feature vectors *i* and *j*, while *σ*_*i*_ is the mean distance between feature vector *i* and all the others. Note that the obtained similarity values are thus normalized across rows, and may not always be identical to absolute ones. For each feature type, we averaged similarity matrices first across scans, and second across subjects, to obtain a population-level description. Each final matrix was subjected to spectral clustering ^86^;*i*.*e*., its eigendecomposition was computed, and the two summarizing dimensions were **u**_2_ and **u**_3_ (*i*.*e*., the second and third eigenvectors).

### Feature values

In addition to examining pattern similarity, we quantified the impact of factors of variation on feature values *per se* using a mixed-effects model, where factors of variation were treated as fixed effects and subject identity as a random effect. In more details, if *F*_*r*_ denotes the feature value of interest for a given coefficient *r* (that is, a given frequency for spectral domain metrics, or region for regional domain metrics), we have a total of 4*S* ·*n*_*p*_ data points available (four scans per subject, with output values for *n*_*p*_ parameter combinations), and we assume the following model:

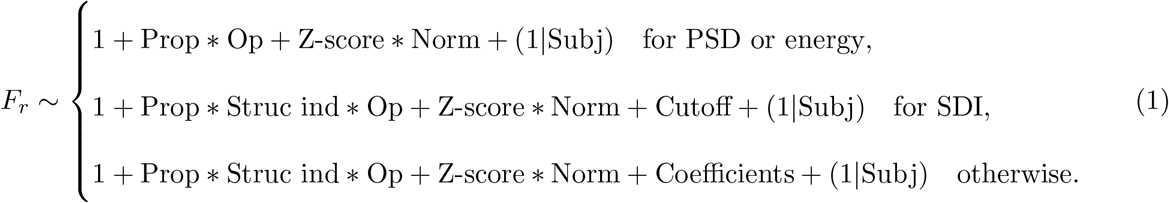

In the above model, we included interactions between structural factors of variation (*Property, Structural individuality, Operator*) and between functional factors of variation (*Temporal z-scoring, Spectral normalization*). Note that for spectral domain features, we cannot consider the type of SC (individual versus group-wise), as eigenmodes with the same index may not represent the same pattern in the former case. For the relevant approaches, we also study the impact of the extra parameter (number of coefficients retained in filtering for alignment/liberality, cutoff frequency selection scheme for SDI). The last term is a subject-specific intercept. The same approach was applied for each spectral frequency/region. All *p*-values associated to the statistical analysis were Bonferroni-corrected for 5*R* parallel tests (5 feature types times *R* regions/spectral frequencies).

In addition to information regarding the modelled fixed effects, for each GSP feature type, we are also provided with an *S × R* matrix summarizing the intensity of random effects across subjects and frequencies/regions. Put simply, a given coefficient (*s,r*) in this matrix quantifies how much should be added or subtracted to the *r*^th^ feature value of subject *s*, across the four associated scans, in complement to the impacts due to the factors of variation (that is, the fixed effects in our model). We considered this representation as an introductory insight into cross-subject variability, computing (1) the subject-wise mean across frequencies/regions as a proxy for how much a given subject would differ from the population average, and (2) the subject-wise standard deviation across frequencies/regions as an estimate of *richness* across individual feature values. We also quantified the range of values taken by cross-subject similarity (Pearson’s correlation) of the *R*-dimensional vectors of random effects.

### Comparison in terms of quality criteria

After examining the impacts of factors of variation on GSP features, we sought to come up with practical recommendations regarding the best GSP features, as well as the optimal parameter choices, as a function of a battery of relevant quality criteria spanning three core domains: robustness of the values to external sources of bias, potential for subject fingerprinting, and generalizability. Below, we outline how all devised quality criteria were constructed, and how approaches were eventually compared.

### Robustness to external sources of bias

Our first criterion regarding robustness made use of the fact that fMRI scans were acquired in pairs over two different days. Both scans from the same day only differ based on their phase encoding direction, an acquisition parameter; thus, a robust feature type should yield similar outputs across these two cases. Two scans acquired with similar phase encoding direction on separate days, however, may yield more variable outputs as a function of the extent of encoded state-like brain activity. Based on these notions, we computed cross-scan similarity (Pearson’s correlation coefficient) between feature vectors of same-day sessions, and feature vectors of different-day sessions with similar phase encoding direction, respectively termed *R*_=_ and *R* _*≠*_. For a given feature type and parameter combination case, *R*_=_ and *R*_≠_ both have size *S ×* 2 (as there are two pairs of sessions each time). We averaged *R*_=_ and *R* _*≠*_ across session pairs into 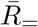 and 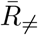, and then computed 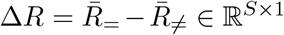; a negative value Δ*R*(*s*) means that for subject *s*, similarity was larger for sessions acquired on a different day, which should ideally never happen. We thus devised a penalty measure: 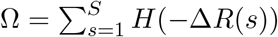, where *H*(*x*) is the Heaviside function. The penalty is larger if more subjects show negative Δ*R* values, and/or if these values are more strongly negative. Because the range of similarity values would differ across feature types, to enable a fair comparison, we also quantified the percentage of subjects for whom Δ*R*(*s*) was negative (*P*_Δ*R<*0_ ∈ [0, 100]) as our first quality criterion regarding robustness to acquisition settings.

In addition, we examined the sensitivity of feature vectors to head movement, a notorious confound in fMRI analyses ^71,72^. For this purpose, for a given feature type and parameter combination case, for each scan at hand, mean FD was separately correlated (Pearson’s correlation coefficient) with the data from each feature coefficient. Then, we computed the mean percentage of explained variance by mean FD across coefficients, and averaged it across scans. Because some feature coefficients were not used for our final assessment of fingerprinting potential (see below for details), as a second quality criterion regarding sensitivity to head movement, we considered the same quantity, but averaged only within the coefficients selected for fingerprinting.

### Fingerprinting criteria

Two different parameter combination cases may be equally robust, yet differ in their ability to distinguish across subjects. Indeed, for this latter point to be examined, *intra-subject variability* (studied above) must be contrasted to *inter-subject variability*, and the latter should be higher. To quantify this, we computed the intra-class correlation coefficient (ICC)^88,89^, considering a group as the four scans available for a given subject. We computed the degree of absolute agreement among measurement (MATLAB function ICC.m^1^, A-1 setting). As some feature coefficients were not used for our final assessment of fingerprinting potential (see below for details), as a third quality criterion reflective of cross-subject discriminability, we considered ICC averaged only within the coefficients selected for fingerprinting.

Next, we examined the ability to fingerprint subjects from the feature data. We computed cross-subject similarity between full feature vectors, focusing on pairs of session acquired over two different days with similar acquisition settings. The obtained subject-by-subject similarity matrix was used as input for one-to-one matching with the Hungarian algorithm ^90^, and fingerprinting accuracy was simply quantified as the correct percentage of assignments. As the resulting accuracy values were consistently similar across both considered session pairs, we averaged them together prior to analysis.

We then examined whether the use of a more restricted set of feature coefficients could be beneficial to fingerprinting accuracy. For this purpose, mean ICC for each coefficient was computed using SES_1_ data only as a mean to select the most informative features (*i*.*e*., with the largest ICC values). We gradually included more features based on mean ICC, and used SES_2_ data to compute fingerprinting accuracy as above on the restricted feature set, so that double dipping was avoided. The maximal accuracy was taken as a fourth quality criterion for fingerprinting accuracy, and the smallest associated percentage of kept coefficients as a fifth quality criterion for parsimony.

### Generalizability to parcellation changes

As a sixth quality criterion, we considered the change in fingerprinting accuracy resulting from a switch from S3 to S3^−^ data (removal of subcortical and cerebellar areas), or from a switch from S3 to S1 data (downscaling to a more restricted set of regions). The average change across these two cases was taken as measure of interest to quantify generalization capability.

### Comparison across feature types/parameter combinations and ranking

To rank all candidate pipelines, we used the six aforementioned quality criteria. For each, the data across all the cases to compare (that is, feature types and parameter combinations) were concatenated and normalized between 0 and 100, so that in each case, a larger value denotes a better outcome. Then, we obtained an overall ranking score as the average across all 6 criteria. In addition, the top 20 cases resulting in the best fingerprinting accuracies were selected for each feature case, and displayed in a spider plot representation^2^. To exemplify the usefulness of our strategy for other types of fMRI measures, we also computed the same ranking scores for selected graph measures: strength (positive and negative), betweenness centrality, clustering coefficient (positive and negative), and eigenvector centrality, using the Brain Connectivity Toolbox ^4^.

## Results

### Parameter choices strongly influence feature patterns and values

Figure 2. displays the similarity, in a two-dimensional summary space, between the feature vectors obtained across parameter combinations for all examined feature types (panels A to E, left half). This is complemented by indicative feature vectors uniformly sampled throughout the summary space (right half). For all five feature types, parameter choices largely impacted vector patterns, but the most impactful parameters differed from case to case: for spectral domain measures, *Operator* (particularly the use of the modularity matrix as opposed to other choices) and *Temporal z-scoring* exerted the largest influence, while *Structural individuality* and *Spectral normalization* had milder effects. For regional domain measures, *Operator* was also a key factor of variation, contrasting the unnormalized and random walk Laplacians to the modularity matrix and normalized Laplacian. For liberality, *Property* also stood out as an influential factor, discriminating between NF(D) and MFL/MFA. Milder contributors included *Spectral normalization* and *Temporal z-scoring* (for alignment), as well as *Structural individuality* (for liberality). In addition, factors of variation also interacted together in shaping feature vector patterns. As an example of two-way interaction, for PSD, *Property* had a greater impact when z-scoring was not performed. For alignment, a three-way interaction can be seen in that *Spectral normalization* only matters when z-scoring is performed and the random walk or unnormalized Laplacians are considered.

**Figure 2:**
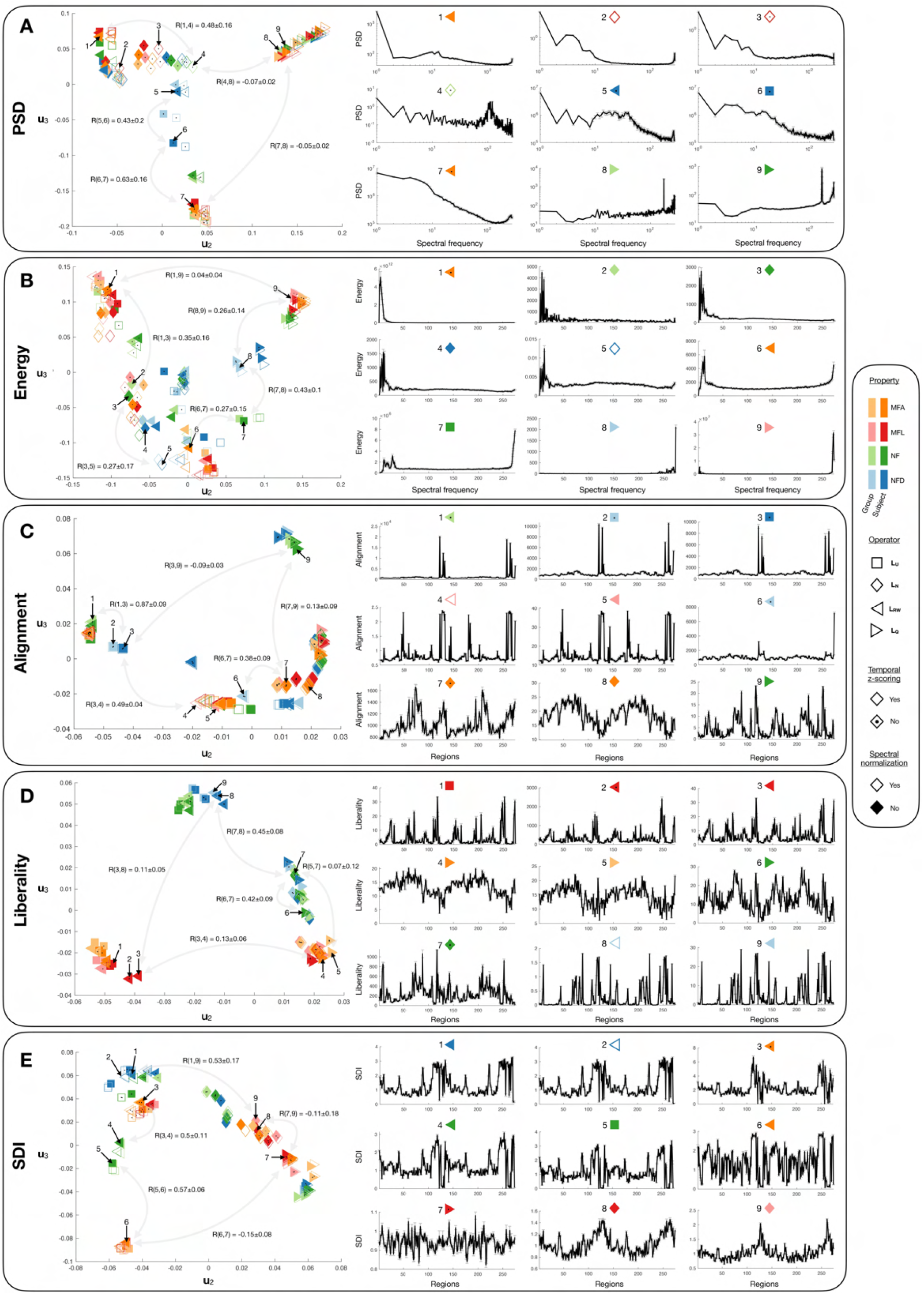
Feature vector patterns are strongly shaped by parameter choices. For PSD **(A)**, energy **(B)**, alignment **(C)**, liberality **(D)** and SDI **(E)**, two-dimensional summary of similarity relationships between feature vectors (left plots), where each data point reflects one particular choice of parameters as summarized in the legend on the right hand side. For each feature type, 9 indicative example feature vectors were sampled from the summary space (numbered from 1 to 9 in each display). Indicative Pearson’s correlation coefficients are highlighted for specific pairs to appreciate how the summary space dimensions encode similarity changes. Example feature vectors are plotted on the right hand side. Note that PSD examples are displayed in log-log scale. Error bars denote standard error of the mean across subjects, computed on data previously averaged across all four available scans.

It can already be appreciated, from **Figure 2**, how the *values* of individual feature coefficients are also largely influenced by parameter choices (see the large differences in scales in the indicative feature vector plots). To thoroughly quantify this, we ran a mixed-effects model analysis for each feature type (**Figure 3** and **Table 1**; see **Supplementary Figure 7** for a visual summary of associated Bonferroni-corrected *p*-values). Overall, the majority of factors of variation and associated interactions exerted significant effects even following Bonferroni correction: for spectral domain measures, more than 95% of coefficients were significant for all fixed effects and interactions. For alignment, there were only less significant coefficients for *Structural individuality* (65.33%), and the interactions of this factor with *Property* (89.78%) and *Operator* (89.78%). For liberality, *Structural individuality* was the only such factor (83.94%). For SDI, in addition to *Structural individuality* (77.37%), there were also *Spectral normalization* (90.51%) and, most largely, its interaction with *Temporal z-scoring*, for which only 37.23% of coefficients reached significance.

**Table 1:** For the scale 3 atlas, summary of mixed-effects model results across feature types for all investigated fixed effects. The *Extra parameter* field stands for *Coefficients* for alignment/liberality, and *Cutoff frequency* for SDI. F-statistics are provided as mean *±* standard deviation across coefficients, complemented by the minimum, median and maximum in brackets. The percentages of frequencies/regions for which significance was reached following Bonferroni correction are displayed in parentheses.

|  | PSD | Energy | Alignment | Liberality | SDI |
| --- | --- | --- | --- | --- | --- |
| Intercept | 4647.8 ± 1521.6 (100%)<br>[271.63, 5051.5, 6982.7] | 611.67 ± 265.23 (99.64%)<br>[271.63, 5051.5, 6982.7] | 3656.4 ± 1436.3 (100%)<br>[271.69, 3834.6, 6505.8] | 7115.6 ± 2505.5 (100%)<br>[1860.9, 6929.8, 14342] | 14658 ± 7082.6 (100%)<br>[1206.6, 13684, 40553] |
| Property (P) | 436.3 ± 280.67 (99.64%)<br>[0.18, 373.11, 1797.5] | 330.67 ± 96.98 (100%)<br>[21.94, 345.88, 700.39] | 190.12 ± 133.2 (97.81%)<br>[0.17, 177.19, 772.87] | 7636.5 ± 6127 (100%)<br>[249.83, 5968.6, 35576] | 9538.2 ± 9910.7 (100%)<br>[8.48, 5453.6, 40308] |
| Struct. indiv. (S) | n.a. | n.a. | 66.77 ± 79.31 (65.33%)<br>[1.19 · 10<sup>-4</sup>, 36.28, 402.18] | 513.88 ± 684.96 (83.94%)<br>[0.02, 230.71, 3374.9] | 434.92 ± 586.43 (77.37%)<br>[0.002, 171.24, 3657.6] |
| Operator (O) | 801.64 ± 407.34 (100%)<br>[10.89, 729.67, 2060.6] | 519.75 ± 173.55 (100%)<br>[16.99, 524.83, 902.09] | 509.08 ± 338.92 (100%)<br>[32.38, 456.1, 1447.4] | 20196 ± 9893.5 (100%)<br>[365.46, 22766, 34810] | 39858 ± 62735 (100%)<br>[271.09, 19695, 3.9 · 10<sup>5</sup>] |
| Z-scoring (Z) | 14005 ± 9467 (100%)<br>[550.08, 12368, 47987] | 738.85 ± 277.97 (100%)<br>[52.15, 717.98, 1799.9] | 8869.7 ± 3732.1 (100%)<br>[769.57, 9685.5, 18236] | 1.9 · 10<sup>5</sup> ± 1.19 · 10<sup>5</sup> (100%)<br>[32204, 1.69 · 10<sup>5</sup>, 1.51 · 10<sup>6</sup>] | 11549 ± 11657 (99.27%)<br>[0.91, 7935.1, 54326] |
| Normalization (N) | 14019 ± 9477.7 (100%)<br>[550.9, 12381, 48036] | 738.85 ± 277.97 (100%)<br>[52.15, 717.98, 1799.9] | 8900.9 ± 3750.6 (100%)<br>[771.93, 9716, 18291] | 2.05 · 10<sup>5</sup> ± 1.29 · 10<sup>5</sup> (100%)<br>[33380, 1.8 · 10<sup>5</sup>, 1.64 · 10<sup>6</sup>] | 396.18 ± 510.14 (90.51%)<br>[0.02, 243.83, 3285.7] |
| Extra parameter | n.a. | n.a. | 8090.4 ± 3210.7 (100%)<br>[733.69, 8794.9, 13087] | 2682.4 ± 1108.2 (100%)<br>[824.24, 2432.5, 7271.9] | 5065.7 ± 3526.7 (100%)<br>[105.85, 4541.5, 15894] |
| P * S | n.a. | n.a. | 56.68 ± 55.27 (89.78%)<br>[0.002, 40.25, 376.83] | 177.83 ± 207.34 (95.99%)<br>[1.24, 95.27, 1233.5] | 161.01 ± 181.86 (96.35%)<br>[0.26, 109.56, 1391.2] |
| P * O | 349.37 ± 114.36 (99.64%)<br>[0.14, 346.1, 760.5] | 279.42 ± 85.74 (100%)<br>[7.45, 280.73, 456.98] | 150.39 ± 90.46 (99.64%)<br>[0.15, 124.67, 438.83] | 1842.3 ± 1423.3 (100%)<br>[165.05, 1342, 7102.1] | 5293.3 ± 5086.2 (100%)<br>[76.61, 3048.5, 20283] |
| S * O | n.a. | n.a. | 55.12 ± 49.84 (89.78%)<br>[0.014, 39.58, 248.14] | 261.94 ± 259.3 (99.27%)<br>[3.73, 190.63, 1466.3] | 194 ± 193.15 (97.45%)<br>[2.2, 136.8, 1401.2] |
| Z * N | 14005 ± 9467 (100%)<br>[550.08, 12368, 47987] | 738.85 ± 277.97 (100%)<br>[52.15, 717.98, 1799.9] | 8869.9 ± 3732.1 (100%)<br>[769.59, 9685.8, 18228] | 1.91 · 10<sup>5</sup> ± 1.19 · 10<sup>5</sup> (100%)<br>[32171, 1.69 · 10<sup>5</sup>, 1.51 · 10<sup>6</sup>] | 23.26 ± 40.51 (37.23%)<br>[4.87 · 10<sup>-4</sup>, 8.53, 356.88] |
| P * S * O | n.a. | n.a. | 52.9 ± 43.85 (97.81%)<br>[0.003, 40.76, 258.41] | 90.11 ± 102.6 (100%)<br>[8.35, 56.99, 659.78] | 97.74 ± 82.41 (100%)<br>[4.58, 75, 471.23] |

**Figure 3:**
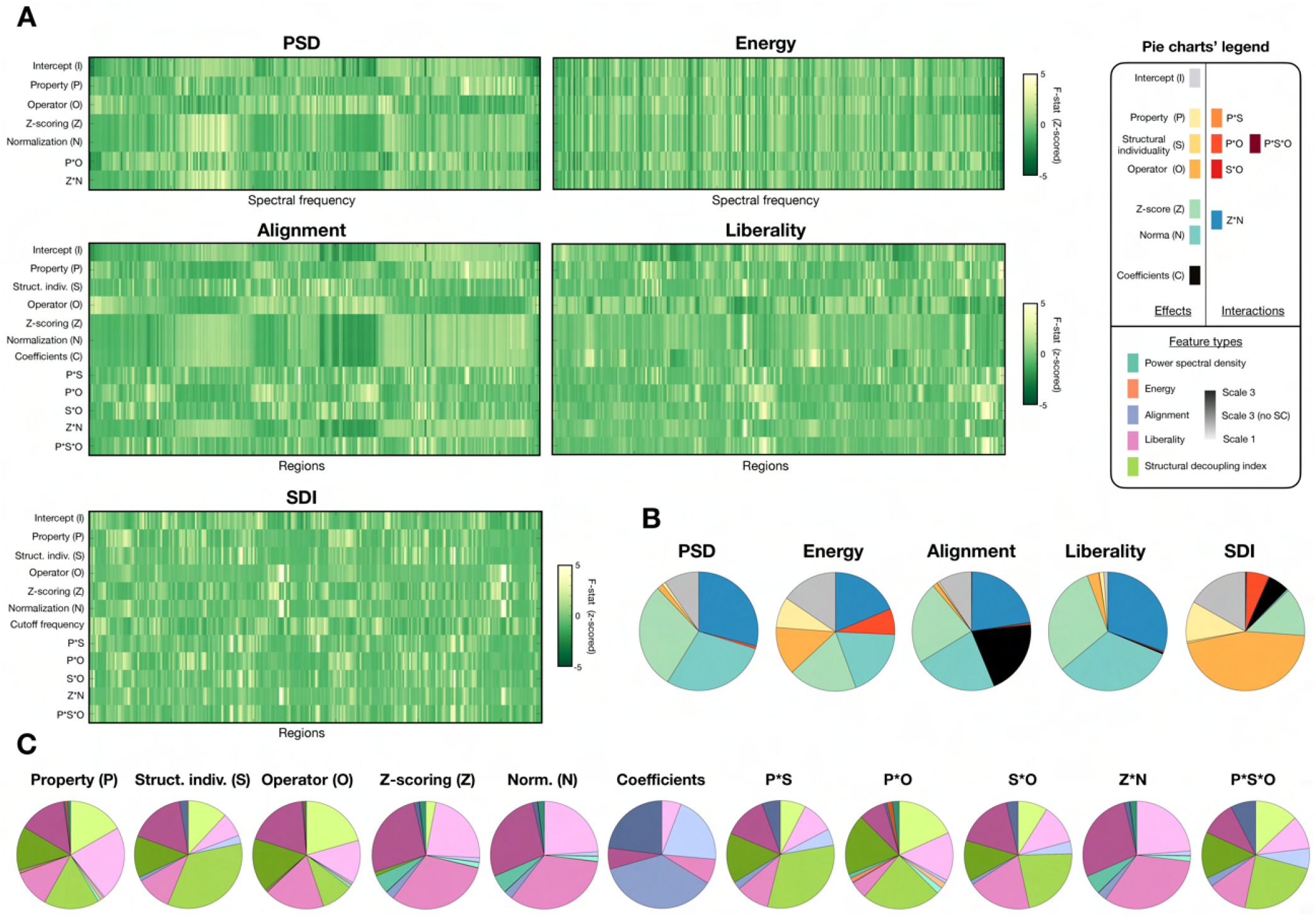
Feature coefficient values are strongly shaped by parameter choices. **(A)** For all five feature types of interest, F-statistics reflecting the strength of the examined fixed effects (*i*.*e*., factors of variation and their interactions). Note that for spectral domain metrics, *Structural individuality* cannot be examined because when using subject-specific SCs, eigenmodes with similar indices may differ across subjects. The values shown in the heatmaps are z-scored in each row to enable a comparison of the underlying patterns; actual F-statistics, and associated *p*-values, are summarized in **Table 1** and **Supplementary Figure 7. (B)** For each feature type, pie chart summary of the respective contributions of individual fixed effects, quantified as the average F-statistic across individual coefficients. Structural factors and their interactions are shown in warm colors, while functional factors and their interactions are displayed in cold colors. Additional parameters (number of coefficients or cutoff selection scheme) are shown in black. **(C)** For each fixed effect of interest, pie chart summary of the respective impact on all feature types of interest, quantified as the average F-statistic across individual coefficients. PSD is shown in turquoise, energy in salmon, alignment in blue, liberality in pink and SDI in green, while the shade highlights which data type was considered between S3 (darkest shade), S3^-^ (intermediate shade) and S1 (lightest shade). Note that spectral domain features do not contribute to the pie chart for *Structural individuality*, and that only alignment and liberality contribute to the pie chart for *Coefficients*.

In terms of patterns of F-statistics across coefficients (**Figure 3A**), *Temporal z-scoring* and *Spectral normalization* almost always exerted similar impacts, and the same was seen for *Coefficients* for alignment. The only exception was for the SDI, where *Spectral normalization* insteadshowed a pattern similar to *Operator*. Comparing the respective impacts of factors of variation and their interactions on each feature type (**Figure 3B**), for PSD, functional factors largely dominated, while for energy, their influence was on par with that of *Property, Operator* and their interaction. Functional factors also dominated for alignment and liberality, together with *Coefficients* in the former case. SDI was the only feature type for which structural factors dominated (*Property, Operator* and their interaction). When instead contrasting the impacts of a given factor across feature types (**Figure 3C**), for structural factors and their interactions, liberality and SDI were by far the most impacted. In the SDI case, the absence of cerebellum and subcortex (S3^-^ input data) rendered *Structural individuality* and its interactions with other structural factors more influential compared to S3 and S1 cases. For functional factors, liberality was the most modulated feature, while alignment was more modulated than liberality in terms of *Coefficients*.

### Preliminary insights into fingerprinting potential of candidate features with random effects

Turning to the random effects obtained upon modelling (**Figure 4A**), for all feature types, there were clear differences in the values obtained across different subjects, reflecting the fact that beyond the impacts of factors of variation, there is also variability in the data linked to subject-specific attributes. For PSD and alignment, given subjects were overall associated to similar values regardless of the exact eigenmode/region at hand (resulting in uniformly colored rows), while for energy, liberality and SDI, there was more diversity. To further ascertain similarities between feature types, we computed the mean and the standard deviation of random effects across coefficients, respectively informing on how much a subject is pinpointed as different from the group average on the whole, and on how rich (how variable) the assessment is across individual coefficients. We then quantified the correlations between the resulting vectors (**Figure 4B** and **Table 2**).

**Table 2:** Correlations between patterns of random effects across features (PSD, energy, alignment, liberality, SDI) and input data types (S3, S3^-^, S1). Only comparisons across input data types for the same feature type, or across feature types for the same input data type, were considered. Entries above the main diagonal reflect similarities in terms of pinpointed subjects overall (mean of random effect patterns across coefficients, corresponding to the left matrix in **Figure 4B**), while entries below the main diagonal denote similarities in terms of richness (standard deviation of random effect patterns across coefficients, corresponding to the right matrix in **Figure 4B**). **: *p <* 10^−3^, *: *p <* 0.05.

| $\sigma \backslash \mu$ | | PSD | | | Energy | | | Alignment | | | Liberality | | | SDI | | |
| --- | --- | --- | --- | --- | --- | --- | --- | --- | --- | --- | --- | --- | --- | --- | --- | --- |
|  |  | S3 | S3<sup>-</sup> | S1 | S3 | S3<sup>-</sup> | S1 | S3 | S3<sup>-</sup> | S1 | S3 | S3<sup>-</sup> | S3 | S3 | S3<sup>-</sup> | S1 |
| PSD | S3 | - | 0.82** | 0.9** | 0.82** | - | - | 1.0** | - | - | 0.93** | - | - | -0.43** | - | - |
|  | S3<sup>-</sup> | 0.97** | - | 0.48** | - | 0.75** | - | - | 0.93** | - | - | 0.9** | - | - | -0.05 | - |
|  | S1 | 0.89** | 0.78** | - | - | - | 0.88** | - | - | 1.0** | - | - | 0.87** | - | - | -0.12 |
| Energy | S3 | 0.34 | - | - | - | 0.59** | 0.96** | 0.84** | - | - | 0.65** | - | - | -0.11 | - | - |
|  | S3<sup>-</sup> | - | 0.29 | - | 0.54** | - | 0.34* | - | 0.69** | - | - | 0.66** | - | - | -0.01 | - |
|  | S1 | - | - | 0.6** | 0.93** | 0.2 | - | - | - | 0.89** | - | - | 0.66** | - | - | 0.01 |
| Alignment | S3 | 1.0** | - | - | 0.33 | - | - | - | 0.72** | 0.91** | 0.91** | - | - | -0.39* | - | - |
|  | S3<sup>-</sup> | - | 0.85** | - | - | 0.2 | - | 0.82** | - | 0.41** | - | 0.94** | - | - | -0.11 | - |
|  | S1 | - | - | 0.98** | - | - | 0.6** | 0.88** | 0.59** | - | - | - | 0.86** | - | - | -0.11 |
| Liberality | S3 | 0.68** | - | - | 0.38* | - | - | 0.68** | - | - | - | 0.88** | 0.95** | -0.46** | - | - |
|  | S3<sup>-</sup> | - | 0.51** | - | - | 0.44** | - | - | 0.47** | - | 0.78** | - | 0.74** | - | -0.06 | - |
|  | S1 | - | - | 0.54** | - | - | 0.19 | - | - | 0.54** | 0.91** | 0.72** | - | - | - | -0.33 |
| SDI | S3 | 0.52** | - | - | 0.13 | - | - | 0.52** | - | - | 0.32 | - | - | - | 0.09 | 0.95** |
|  | S3<sup>-</sup> | - | 0.63** | - | - | 0.1 | - | - | 0.67** | - | - | 0.27 | - | 0.71** | - | -0.05 |
|  | S1 | - | - | 0.47** | - | - | 0.03 | - | - | 0.46** | - | - | 0.32 | 0.92** | 0.6** | - |

**Figure 4:**
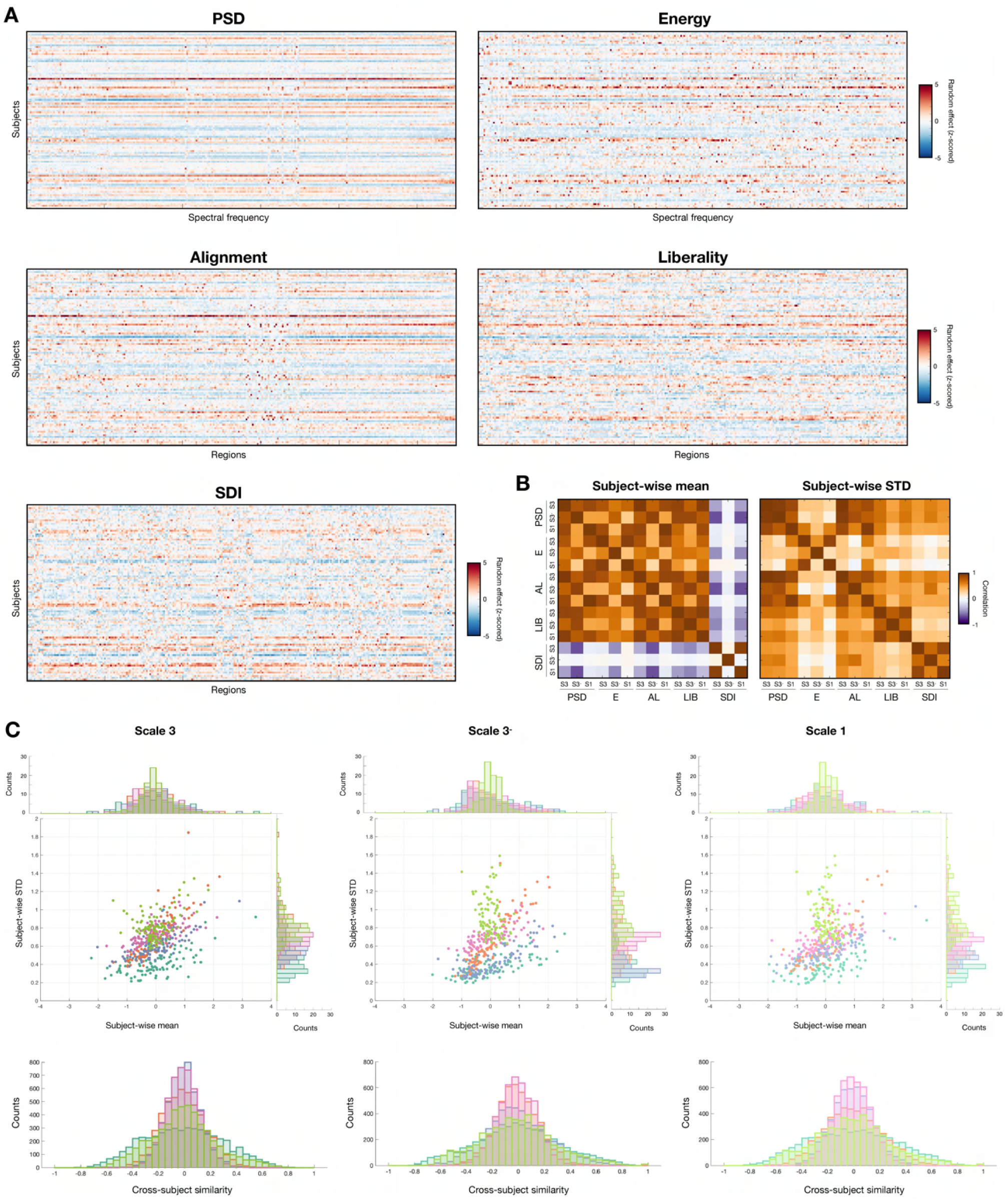
Similarities and differences between feature types in terms of random effects. **(A)** For all five feature types of interest, random effects across subjects (rows in each heatmap) and individual feature coefficients (columns). Values are z-scored across subjects. **(B)** Following the computation of the mean (left matrix) or standard deviation (right matrix) of random effects across coefficients, Pearson’s correlations between the resulting vectors across feature and input data types. **(C)** (Top) Across input data types (S3: left column, S3^-^: middle column, S1: right column), subject-wise mean (X-axis) and standard deviation (Y-axis) of random effects across coefficients, where each data point stands for one subject and a given feature type. Data points are colored in turquoise for PSD, salmon for energy, blue for alignment, pink for liberality and green for SDI. (Bottom) For the same input data types, distributions of cross-subject similarity values when Pearson’s correlation is computed between random effect vectors of different subjects (that is, between different rows from the heatmaps shown in **(A)**). E: energy, AL: alignment, LIB: liberality, STD: standard deviation.

PSD, energy, alignment and liberality each identified similar subjects regardless of which input data was used (S3, S3^-^ or S1; left matrix). For SDI, however, while the same subjects were pinpointed for S1 and S3 (*R*_*S*3,*S*1_ = 0.95, *p <* 10^−3^), the removal of the subcortex and cerebellum completely destroyed any association (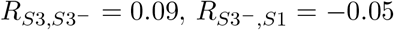, both *p* = 1). Comparing across features, when the same input data was used, PSD, energy, alignment and liberality always identified the same subjects as well, while for SDI, a significant, negative-valued association was only observed for S3 between SDI and PSD (*R* = −0.43, *p <* 10^−3^), alignment (*R* = −0.39, *p <* 10^−3^) or liberality (*R* = −0.46, *p <* 10^−3^). The subjects exhibiting the greatest richness in information were almost always similar across input data types, to the exception of energy for S1 compared to S3^-^ (*R* = 0.2, *p* = 1; right matrix). Across features, PSD, alignment and liberality consistently yielded similar outcomes, while SDI was similar to PSD and alignment, but not liberality. Energy clearly stood out, as the associations with other feature types were not consistent from a data type to another, only reaching significance in few occasions.

In addition to correlational assessments, we plotted individual data points in a two-dimensional space where the X-axis is subject-wise mean (how strongly a subject is pinpointed), and the Y-axis stands for subject-wise standard deviation (how variable or rich random effects were across coefficients; **Figure 4C**, top row). For S3, S3^-^ and S1 input data, SDI always exhibited the lowest mean and highest standard deviation values. Conversely, standard deviation was consistently the lowest for PSD. Despite these opposite properties, PSD and SDI were also the two feature types for which cross-subject similarity values in random effect vectors exhibited the largest dynamic range (**Figure 4C**, bottom row).

### Robustness to acquisition settings and head movement

To assess robustness to acquisition settings, we examined similarity between scans acquired on the same day (with different phase encoding directions) compared to similarity between scans acquired with similar settings, but across two separate days. **Figure 5A** schematically recapitulates how, from this data, quality metrics were extracted: on top of mean similarity between same-day sessions (X-axis in panels B to F), we quantified the percentage of subjects for whom similarity between different-day sessions was larger (X-axis in panels G to K; *i*.*e*.,the data points lying in the upper left half of the display, which we refer to as *percentage of incongruent cases P*_*I*_ because if a measure is robust to acquisition settings, similarity between same-day sessions should be larger), as well as the sum of distances from the diagonal (Y-axis in panels B to F, symbolized by the vertical lines; we call this latter quality criterion the *penalty* Ω). To study robustness to head movement, we computed the mean percentage of variance explained by average FD (*P*_FD_; Y-axis in panels G to K).

**Figure 5:**
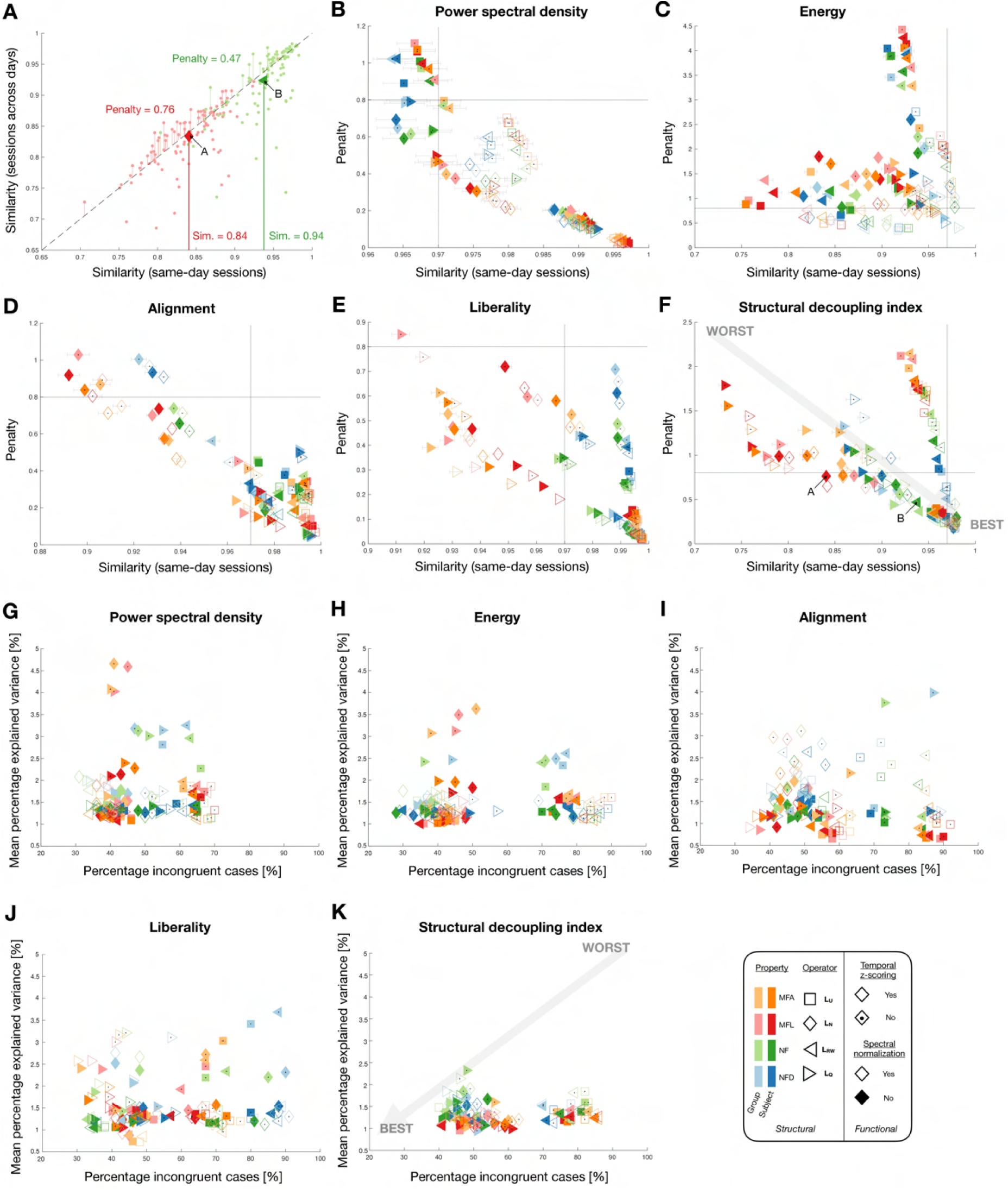
Robustness to acquisition settings and head movement. **(A)** Illustration of how quality measures regarding robustness to acquisition settings were computed for two indicative cases (red and green data points, representing two SDI parameter combinations and respectively labeled A and B). Average similarity of feature vectors between same-day sessions across subjects is considered as a first quality measure, and is the X-axis in panels **(B)** to **(F)**. The Y-axis in the same panels stands for the penalty Ω, taken as the sum, across subjects, of differences between across-day and same-day similarity values for cases when sessions acquired on two separate days were more similar (denoting an impact of acquisition settings, and symbolized by the vertical lines in panel **(A)**). The percentage of subjects for which this applied (so called *incongruent* cases) makes the X-axis in panels **(G)** to **(K)**, while the Y-axis depicts the average percentage of variance explained by mean framewise displacement. **(B-F)** Average similarity between same-day sessions across subjects, and penalty Ω, for the five feature types of interest. Error bars depict standard error of the mean. The light grey horizontal and vertical bars highlight values of 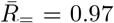 and Ω = 0.8 to enable comparison across panels despite the different ranges. The best parameter combinations yield data points at the bottom right of the displays (large similarity between same-day sessions combined with low penalty). **(G-K)** Percentage of incongruent cases *P*_*I*_, and average percentage of variance explained by mean framewise displacement *P*_FD_. The best parameter combinations yield data points at the bottom left of the displays (low *P*_*I*_ and *P*_FD_).

For all feature types, similarity between same-day sessions reached values up to 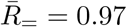 for the best parameter combinations. Similarity could go down to 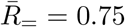 for energy and SDI, but never much lower than 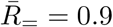 in other cases. For Ω, values were larger overall for energy and SDI. For *P*_*I*_, PSD stood out as no parameter combination yielded values larger than 70%, against up to 90% for other feature types. Furthermore, the best cases yielded down to *P*_*I*_ = 30% for all feature types but SDI, where only 40% could be reached. As for head movement, mean FD explained as little as 0.6% in some alignment and liberality cases, against 1% at best for other feature types. Furthermore, explained variance of more than 2% when z-scoring was not performed was seen for all feature types except SDI. In fact, the absence of z-scoring yielded bad outcomes across all examined quality criteria and feature types, with the exception of energy for same-day sessions’ similarity, which improved overall without z-scoring. Normalization only tangibly impacted spectral domain metrics, in a way that differed across quality criteria: it improved similarity and penalty for PSD and energy, and incongruency was specifically improved for PSD in the most robust subcases. The use of the normalized Laplacian was particularly detrimental to alignment, while for liberality and SDI, it also extended to the modularity matrix.

For PSD, optimal parameter choices across robustness quality criteria appeared to be the use of MFL or MFA combined with the unnormalized or the random walk Laplacian and z-scoring (across compatible combinations, 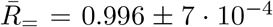, Ω = 0.035 *±* 0.012, *P*_*I*_ = 40.75 *±* 3.04%,*P*_FD_ = 1.15 *±* 0.15%). For energy, there was no clear consensus across criteria, but the combination of the modularity matrix and normalization helped for similarity and penalty, while the use of (N)FD yielded better incongruency but worse sensitivity to mean FD compared to MFL/MFA. For alignment, NFD or MFL with the modularity matrix and z-scoring were the consensus best choices (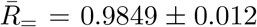, Ω = 0.0947 *±* 0.0518, *P*_*I*_ = 39 *±* 2.56%, *P*_FD_ = 1.2 *±* 0.43%). For liberality, NF combined with the unnormalized or random walk Laplacians and z-scoring performed best (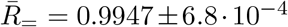, Ω = 0.0392 *±* 0.0048, *P*_*I*_ = 36.13 *±* 2.7%, *P*_FD_ = 1.26 *±* 0.42%), while for SDI, the combination of MFL and random walk Laplacian or NFD and normalized Laplacian both offered good performance across criteria (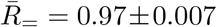, Ω = 0.25*±*0.06, *P*_*I*_ = 45*±*3.55%, *P*_FD_ = 1.3 *±* 0.27%).

### Fingerprinting ability

To determine to what extent fingerprinting could be achieved, we started by contrasting inter-individual differences to intra-individual variability by quantifying the intra-class coefficient (ICC) and contrasting it to 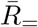 (**Figure 6**, right half). For all feature types, mean ICC across coefficients differed even between parameter combinations that yielded similar 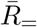, showing that robustness does not necessarily imply subject discriminability. The most impactful factor of variation regarding mean ICC was clearly the use of a subject-specific SC as opposed to a group-wise one: in the former case, discriminability was consistently enhanced. For PSD, energy and liberality, the absence of z-scoring also tended to yield larger mean ICC values. At the level of individual feature coefficients (**Figure 6**, left half), the whole set exhibited large ICC values when a subject-specific SC was used, consistently across feature types. When resorting to a group-wise SC, however, only sparser sets of coefficients had high ICC values, hinting at the fact that when a common structural basis is used, only specific features may enable to distinguish between distinct individuals.

**Figure 6:**
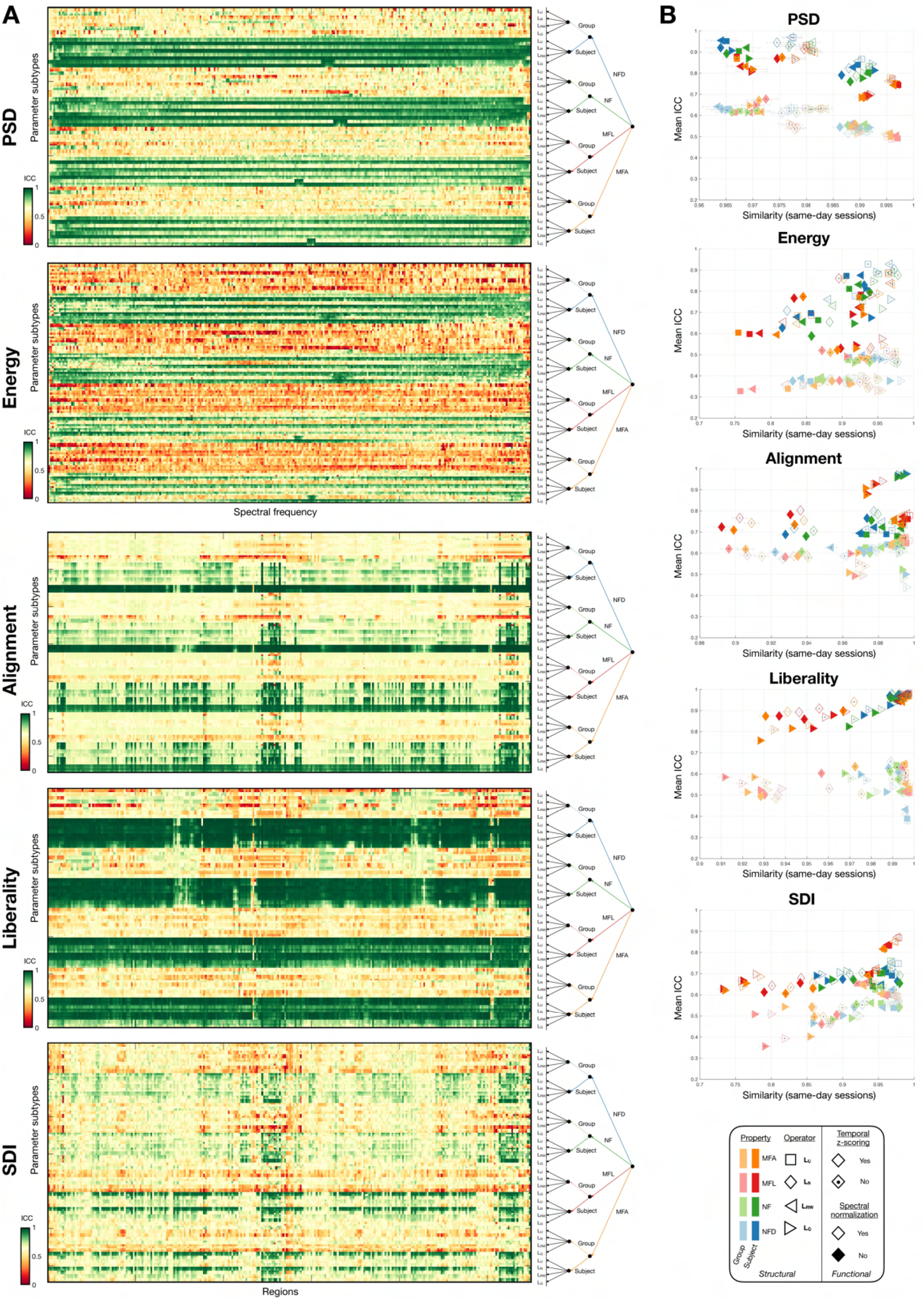
Disentangling intra-subject variability from inter-individual differences. **(A)** For the five feature types of interest, intra-class coefficient (ICC) values across parameter combinations (rows in each heatmap) and individual feature coefficients (columns). A decision tree on the right of each heatmap summarizes which parameter settings applied to each row of the displays. **(B)** For the five feature types of interest, similarity between same-day sessions (a measure of intra-subject variability) is contrasted to the mean ICC across coefficients, which is larger when cross-subject differences more strongly outweight intra-subject variability.

Next, we quantified fingerprinting accuracy, first when using the whole set of available feature coefficients and contrasting the results as a function of input data type (S3, S3^-^ or S1; **Figure 7**). When using a subject-specific SC (data points with darker colour shades), fingerprinting accuracy reached 100% across a breadth of parameter combinations for all feature types, regardless of input data type. However, the values were the most scattered in the case of energy, where accuracy sometimes dipped below 20%, while conversely, liberality outcomes almost always remained at 100%. Turning to data points associated to a group-wise SC (lighter colour shades), there was a much bigger variability in reached accuracy, ranging from 0 to 80% in the majority of cases. Values close to 100% could only be reached in some settings for SDI (*e*.*g*., modularity matrix for NFD with z-scoring and without normalization, yielding an accuracy of above 95% for both S3 and S3^-^).

**Figure 7:**
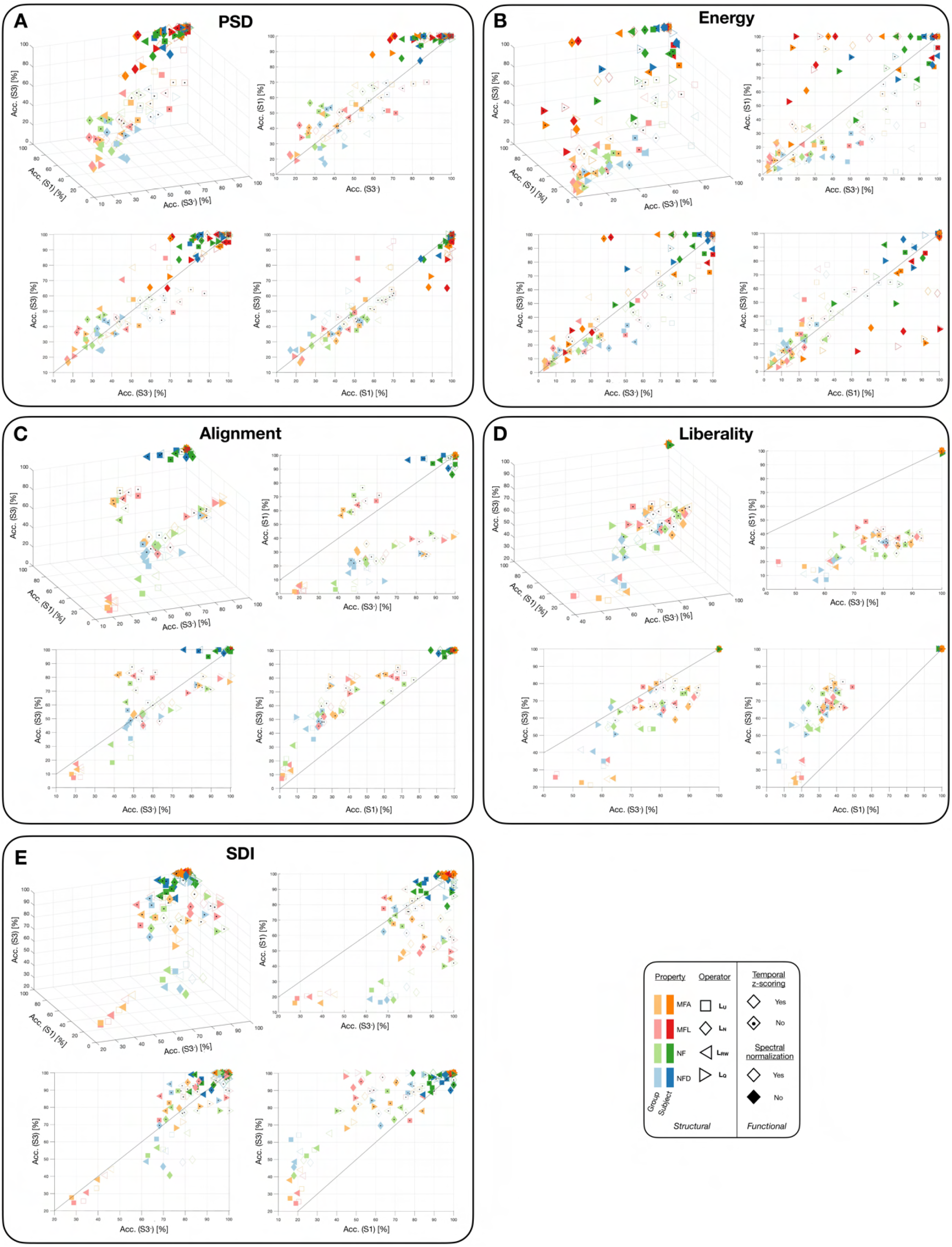
Fingerprinting accuracy across feature and input data types. **A-E** For the five feature types of interest, fingerprinting accuracy is shown when using S3, S3 and S1 input data, as a three-dimensional representation (top left plot) and when focusing on specific pairs (*i*.*e*., individual two-dimensional planes). Each data point stands for one parameter combination, as summarized in the bottom right legend.

Spectral domain metrics performed better when normalization was performed, but did not show clear differences in fingerprinting accuracy as a function of atlas scale or completeness. On the contrary, for alignment, liberality and SDI, fingerprinting was much more effective when working at scale 3 (both S3 and S3^-^). In addition, liberality performed better when the cerebellum and subcortex were not considered (S3^-^).

We then assessed whether the use of a more restricted set of feature coefficients could improve fingerprinting accuracy (**Figure 8**). We observed that in many cases, maximal accuracy was reached when only using a subset of available coefficients (see panel A for indicative examples), and quantified the reached accuracy as well as the parsimony in the associated feature set (panels B-F). As the use of subject-specific SCs led to almost perfect accuracy in most scenarios, here, we focused on cases for which a group-wise SC was used. Perfect fingerprinting accuracy in this setting could be achieved both by PSD and SDI, but with different properties: in the former case, there were at most 50% of coefficients required, while in the latter case, it could range from 10 to 100%. For alignment and liberality, a high accuracy of 90% or so could, in many cases, be reached when the full feature set was leveraged. Finally, energy outcomes fluctuated a lot with no clear pattern, and rarely reached high accuracy. The need of a full set of coefficients for alignment and liberality to enable fingerprinting extended to S3^-^ and S1 cases (see panels G-I). For PSD and SDI, S3^-^ results were comparable to those of S3, but for S1, perfect accuracy was never achieved: in this last case, SDI-based results were the best, reaching up to 90% accuracy and requiring a full set of features to do so.

**Figure 8:**
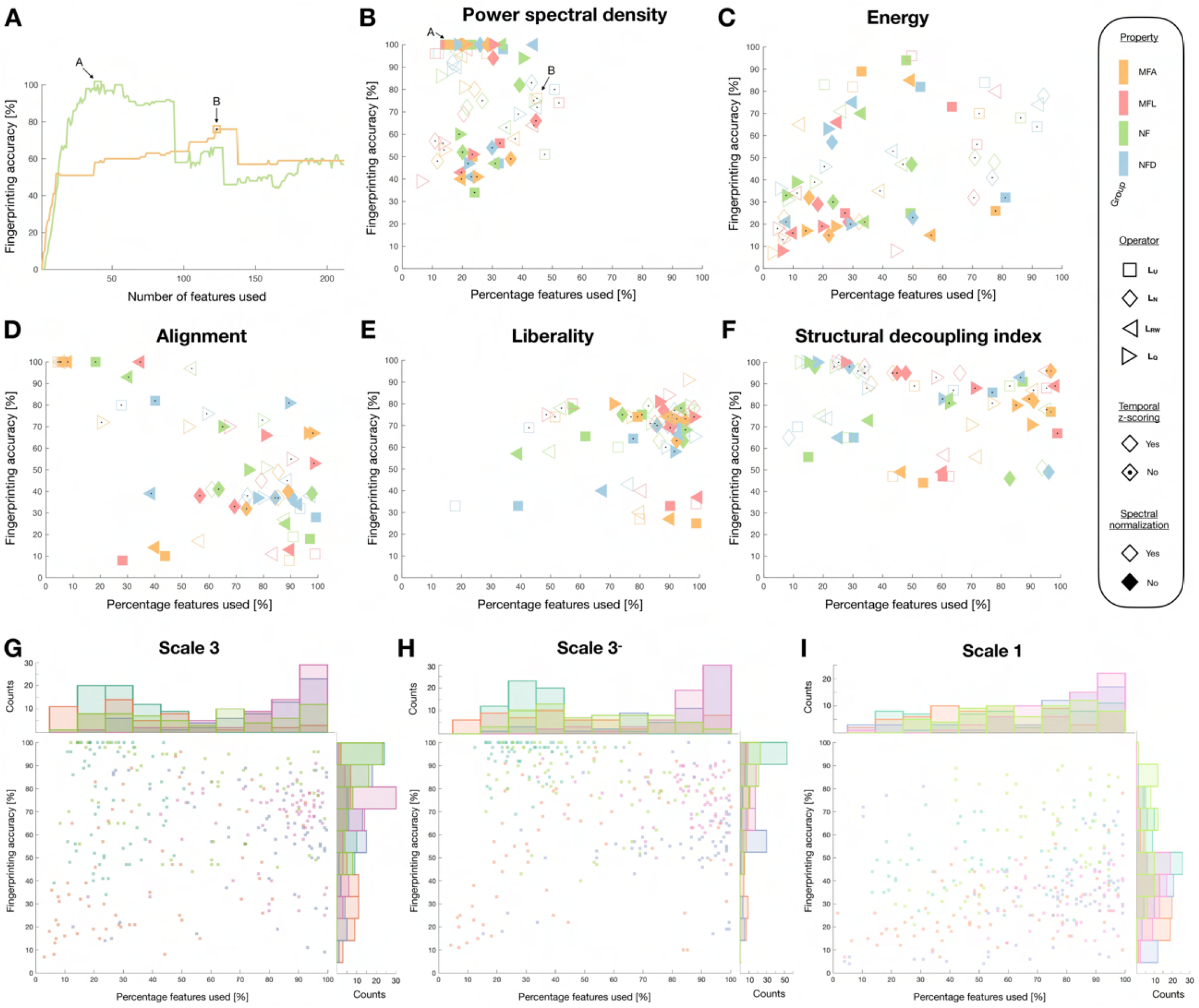
Fingerprinting accuracy is sometimes optimized using a sparse feature set. **(A)** For two indicative cases (green and orange, representing two PSD data points labeled A and B), fingerprinting accuracy as a function of the number of used feature coefficients. **(B-F)** For the five feature types of interest, maximal reached fingerprinting accuracy as a function of the percentage of features used. The S3 input data was considered for these representations, and only parameter combinations associated to the use of a group-wise SC are considered. **(G-I)** Maximal reached fingerprinting accuracy as a function of the percentage of features used for S3 (left), S3^-^ (middle) and S1 (right) input data. Each data point represents one parameter combination for a given feature type (PSD: turquoise, energy: salmon, alignment: blue, liberality: pink, SDI: green).

### Comparison of candidate feature types and parameter combinations

Eventually, we reconsidered all our quality criteria reflective of robustness and fingerprinting ability to converge onto optimal feature types and associated parameter choices (**Figure 9**). First, we ordered all parameter combinations in order of decreasing quality for each of the studied criteria (**Figure 9A**). For robustness to acquisition settings (*P*_*I*_), PSD, energy and liberality yielded the best values, and PSD consistently outperformed all other feature types. For sensitivity to head movement (*P*_FD_), conversely, PSD was the worst overall alongside liberality, while alignment yielded the best cases and SDI was consistently good. Discriminability (computed only on the subset of features used for fingerprinting in **Figure 8**) was consistently the highest for alignment and PSD, while it was the worst for energy. PSD and SDI clearly stood out in terms of fingerprinting accuracy overall, but the former enabled overall more parsimonious representations. However, PSD was also the feature type that generalized the least from S3 to S3^-^ or S1, while SDI was amongst the best. PSD and SDI thus stand out in terms of their fingerprinting abilities, but exhibit different assets and weaknesses. Their superiority is further evidenced in **Figure 9B**: for 61 out of 64 possible parameter combinations, the best consensus rating was achieved by either of the two (50 times for PSD, 11 times for SDI). When decomposing the top 20 cases for which fingerprinting accuracy was maximized as a function of individual quality criteria (**Figure 9C**), the all-around performance of PSD and SDI approaches further stood out, whereas for other feature types, high accuracy could not be achieved while retaining high robustness and generalizability.

**Figure 9:**
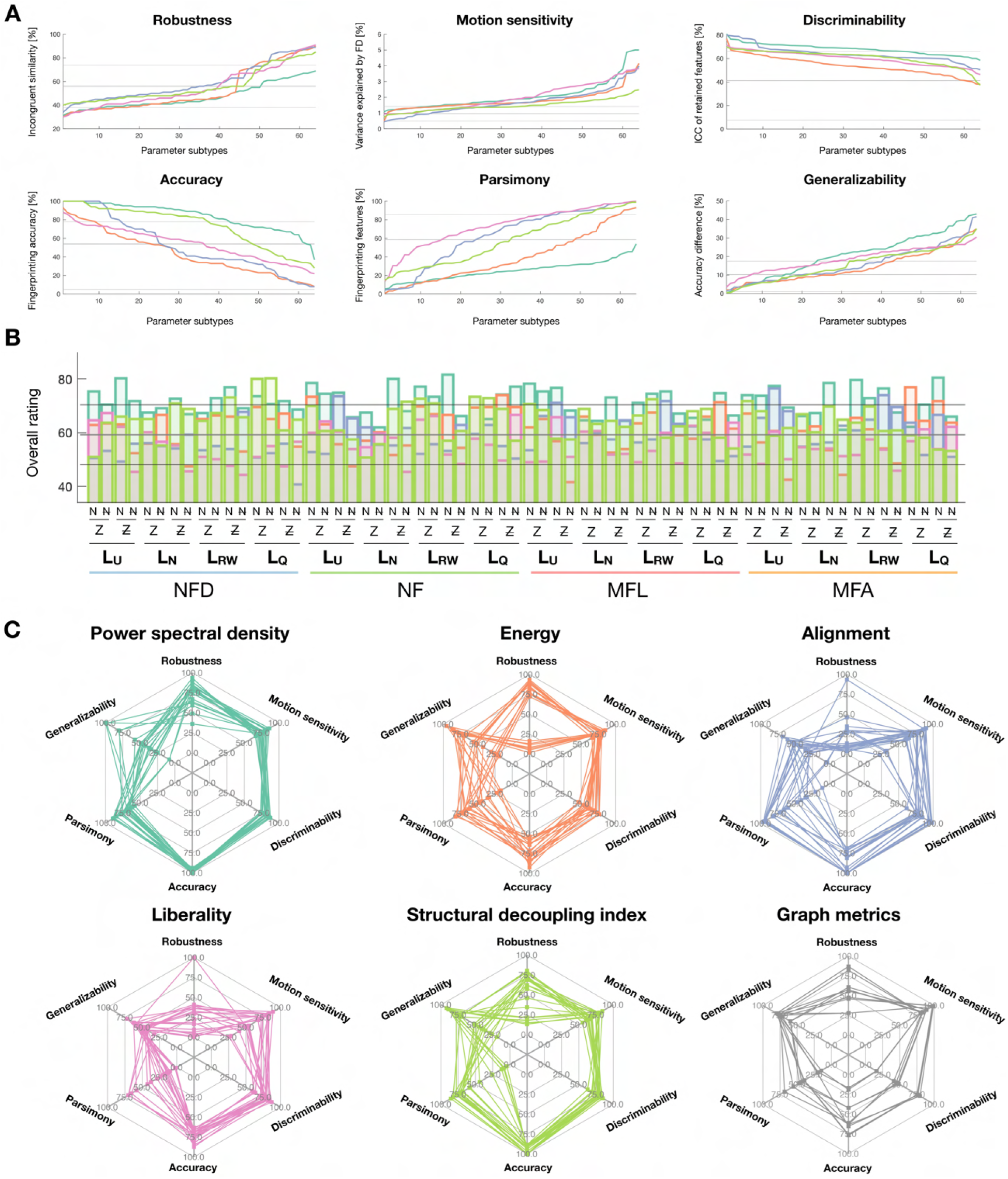
Summary of feature types across investigated quality criteria. **(A)** For the six probed quality metrics, sorted values from best (left) to worst (right) parameter combinations for the five feature types of interest (PSD: turquoise, energy: salmon, alignment: blue, liberality: pink, SDI: green). Only parameter combinations associated to the use of a group-wise SC are considered. Note that better outcomes imply lower percentages of incongruent cases (first plot) and sensitivity to head movement (second plot), a smaller set of feature coefficients used for fingerprinting (fifth plot) and smaller differences between fingerprinting accuracies using S3, S3^-^ and S1 input data types (sixth plot), but larger mean intra-class coefficient values (third plot) and fingerprinting accuracy values (fourth plot). The median, minimum and maximum values reached across the investigated graph metrics cases are also shown by horizontal grey bars. **(B)** Across investigated parameter combinations (from left to right), overall quality score achieved for the five feature types of interest (same color coding as in **(A)**). The median, minimum and maximum values reached across the investigated graph metrics cases are also shown by horizontal grey bars. **(C)** For the five feature types of interest (same color coding as in **(A)**) and graph metrics (grey), summary of the top 20 parameter combinations in terms of fingerprinting accuracy across the six probed quality metrics, where each metric can take values between 0 (worst) and 100 (best).

Finally, to illustrate the potential of our comparative approach for other types of fMRI measures, we quantified the same quality criteria when generating subject-wise functional connectivity matrices, and subsequently computing selected graph metrics (nodal strength, clustering coefficient, betweenness centrality and eigenvector centrality). Fingerprinting accuracy remained below 80% at best, and more than 60% of available coefficients were always required, which reflect poorer performance compared to GSP features. However, graph metrics also yielded more generalizable results, and suffered from particularly little head movement sensitivity, so that in selected parameter combination cases, they could even outperform GSP features (**Figure 9B**).

## Discussion

The combination of structural and functional brain information to improve endeavours such as behavioural prediction or clinical diagnosis has gained popularity in recent years. Here, we focused on graph signal processing as such a multimodal approach. We investigated whether the parameter choices made in the computation of features of interest play an important role, and found that all examined factors of variation were strongly influential. We then conducted an indepth comparative study of existing features and parameter combinations regarding the ability to fingerprint individual subjects. On top of quantifying fingerprinting accuracy *per se*, we also assessed robustness to external sources of variance (acquisition settings and head movement), sensitivity to specificities of the involved brain parcellation, and we probed the dimensionality of the telling feature coefficients. Power spectral density and the structural decoupling index emerged as the most balanced feature types in terms of the assessed criteria.

### All GSP parameters matter and must be carefully selected

We addressed the impact of parameter choices on GSP features at two levels: that of the global feature vector pattern (**Figure 2**), and that of the feature coefficient values (**Figure 3**). Both viewpoints are important to consider, as they relate to dedicated analytical approaches: while methods such as representational similarity analysis ^91^ or kernel ridge regression operate at the level of similarity between full feature vectors, other regression approaches such as the elastic net ^92^ instead weight individual feature coefficients. Both types of methods have often been used, and compared, in structural and functional neuroimaging ^93,94^.

A first important observation was that virtually all the investigated parameters exerted significant impacts, in one way or another, towards the computation of all the studied GSP feature types. This included choices on the structural end of the GSP pipeline, modulations of the functional time courses fed into the analysis, as well as additional parameters specific to only some of the studied feature types. This highlights the logical but often underappreciated fact that in neuroimaging, every single decision made from the raw data to the final product of a research project counts, and should be carefully considered. While not directly addressed here owing to the associated computational burden, we remark that it also extends to preprocessing of the data, whose variants should ideally also be investigated in parallel to choices made within an analytical pipeline of interest.

While all parameters were impactful, it was nonetheless possible to pinpoint a selected few that more largely modulated GSP features. Furthermore, the most influential parameters differed as a function of the feature type at hand. For spectral features (PSD and energy), the use of the modularity matrix strongly reshaped feature vectors compared to other choices. This is explained by the fact that eigenmodes are then sorted in ascending order of *modularity* (not spectral frequency), and the most anti-modular eigenmodes are not necessarily the ones with the smallest spectral frequency. Accordingly, the largest PSD coefficients became shifted to higher indices when using the modularity matrix (see **Supplementary Figure 1**). For regional features, the type of operator was also important, but there was rather a distinction between **L**_**U**_ and **L**_**RW**_ on the one hand, and **L**_**N**_ and **L**_**Q**_ on the other hand. This is likely because the first two conserve the information regarding regional degree, while **L**_**N**_ normalizes it and **L**_**Q**_ does not include it, staying at the level of the adjacency matrix **A**.

Z-scoring and normalization were the two most influential parameters regarding feature values, for all feature types except the SDI. Furthermore, they consistently exerted similar effects on the results in terms of F-statistics. In fact, z-scoring also applies a sort of normalization to each row of the data matrix, subtracting the mean and dividing by the standard deviation. Spectral normalization rather modulates each column of the data matrix (in the spectral domain), subtracting the mean and dividing by the L2-norm. The large impact of both operations on individual feature values is then logical given the division step. Interestingly, z-scoring also exerted strong effects on the patterns of spectral features, leading to the emergence of new subsets of eigenmodes linked to large PSD values when not performed (see **Supplementary Figure 1**). These eigenmodes include subcortical regions, whose time courses of activity take particularly large values when z-scoring is not performed, owing to the greater extent of noise (see **Figure 1C**).

Specifically for liberality, property type was the strongest modulator of feature patterns, contrasting NF(D) to MFL/MFA. This implies that while the lowest frequency eigenmodes resulting from each property type strongly resemble each other (yielding only little impact of property type on alignment feature vectors), there are also more subtle and localized property-specific attributes that can be accessed. From a neurophysiological perspective, the implication is a possible case-specific dissociation between the number of fibers connecting two regions, and the length (MFL) and extent of insulation (MFA) of the tract. In future work, it will be interesting to determine what is the purpose of such a dissociation.

For alignment (but not liberality), the number of retained eigenmodes (*Coefficients* factor of variation) also largely impacted the output feature values. Additional investigations (see **Supplementary Figure 6**) revealed that in fact, feature values would start converging only when including a minimum of 10 eigenmodes. This likely arises because the smallest frequency eigenmodes are directly linked to the canonical resting-state networks, which a sparse subset can efficiently approximate ^33^. When enough eigenmodes are retained to faithfully approximate these networks, the largest contributions to regional time series can be retained in the alignment coefficients, and further broadening of the retained set becomes less impactful.

Structural individuality was the only factor with only mild (albeit significant) impacts on feature patterns and values. For spectral features, the reliance on subject-specific SCs resulted in a smoothening of the feature vectors, which can be explained by the fact that from a subject to the next, the same eigenmode may lie a few indices apart in the set. The impacts of structural individuality could most clearly be seen for liberality and SDI (which is directly tied to liberality), which implies that structural differences between individual subjects involve subtle and localized patterns. We find it a good lesson that while *Structural individuality* was arguably the least impactful factor of variation in terms of feature patterns and values, it was in fact the most crucial determinant of fingerprinting accuracy across all features (**Figure 7**; see below for details). Once more, this highlights the importance of an exhaustive approach when attempting to pinpoint optimal parameter settings.

When contrasting the influence of structural and functional parameters on GSP features, we observed an overall balanced impact on feature vector patterns, but a stronger influence of functional parameters on the majority of GSP feature types for coefficient values. The only exceptions were energy (somehow balanced contributions) and the SDI (domination of structural parameters), putting these two feature types apart and hinting at the specific nature of their properties. For energy, the greater influence of structural choices likely arises from the definition of the metric itself, which involves a multiplication of individual coefficients with their respective eigenvalue *λ*_*i*_, a structural quantity. For SDI, we suspect that the division between liberality and alignment results in the *canceling out* of functional factors.

Comparing the influence of each parameter across features, liberality (for all factors of variation) and SDI (for structural ones) were the most strongly modulated. In the SDI case, interestingly, the absence of subcortex and cerebellum (S3^-^ input data) largely altered the scale of impacts for *Operator* (smaller) and *Structural individuality* (larger), much more so than the switch from S3 to S1. Our analysis of random effects (**Figure 4**) confirmed this, and reinforced the distinct nature of energy and the SDI: indeed, the subjects pinpointed as outliers were completely different for SDI and S3^-^ input data compared to all other cases, and in terms of richness, energy was the most different feature type to all others. The particular sensitivity of the SDI to the set of included brain regions can be seen both as an asset or a drawback: on the one hand, perfect fingerprinting was possible both from S3 and S3^-^ input data with the SDI (**Figure 9, Supplementary Figure 20**), which implies that distinct information of interest is available in both cases and speaks for a particular relevance of the SDI for fingerprinting. On the other hand, the particular sensitivity to included areas raises concerns regarding the robustness of SDI as a measure, which should be more clearly investigated in future studies. As for energy, given its overall mediocre performance across investigated quality criteria as further discussed below, our view is that it exemplifies a metric that becomes too unstable by the joint modulations of structural and functional factors.

In addition to individual impacts, we also observed that all studied parameters exhibited significant interactions. These interactions were not only between structural or functional parameters, but across all of them. They help us discard some candidate strategies when it comes to the design of GSP studies: in particular, one cannot assume that if only one parameter changes with respect to a previous analysis, it will not impact the others. Furthermore, it should discourage from a perturbation-oriented analysis of parameter impacts, when only one parameter is varied while keeping all others fixed.

### PSD and SDI are the most promising GSP features for fingerprinting

A challenge associated to brain data-based fingerprinting is the potentially overlapping contribution of several different factors. First, to reliably detect subject-specific neural features reflected in GSP metrics, stability of the measures across repeated sessions with potentially different acquisition settings, and across distinct data preprocessing strategies, is required. Second, spatiotemporal patterns of head movement inside the scanner have been shown to differ across individuals in ways that directly relate to behaviour^87^, raising the possibility that they could also contribute to spuriously boost fingerprinting performance. Third, if a large set of feature coefficients is leveraged in fingerprinting, overfitting cannot be ruled out without further analyses on yet unseen data.

The quality criteria that we devised were directly aimed at addressing these potentially overlapping influences. First, we explored the stability in GSP measures across sessions acquired on the same day with different phase encoding directions, and across separate days with the same acquisition settings. Robustness to acquisition settings was addressed by means of three metrics: the penalty Ω, the mean similarity across same-day sessions, and the percentage of incongruent cases *P*_*I*_. Only the third one was included in our final comparison, because it enabled to not only compare candidate parameter combinations within a given feature type, but also potential pipelines across feature types. On the other hand, the ranges of Ω and 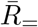 differ by nature between feature types, owing to the specific data structure at hand. For example, similarity values were always the highest for PSD because in this case, there is always a decreasing logarithmic relationship between PSD and eigenvalue index, which inflates correlation estimates.

Looking at the three metrics collectively, in some occasions, the impacts of a given factor of variation were not consistent. For example, spectral normalization for energy tended to improve 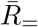 and Ω, but to worsen *P*_*I*_. This provides evidence for the fact that the fraction of subjects showing incongruent features (rendered by *P*_*I*_) may be partly independent from the severity of the dissociations (captured by Ω). A direct implication is the need to devise, for future studies, a more complete set of metrics to quantify robustness, while enabling comparisons across feature types. Importantly, while it is likely that a larger Ω reflects less stable measures (*i*.*e*., a larger impact of acquisition settings), there is an alternative explanation: that of increasingly resolved state-like features of brain function, resulting in lower 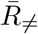 values. In order to explicitly address this, one would ideally require a dataset containing densely sampled individuals, and concurrent behavioural assessments at each scanning time. Based on existing literature, we would however not expect to detect strong state-like effects, as several reports have shown only very limited cross-session variability (on sessions acquired with identical settings) compared to group-wise or individual contributions in terms of overall functional connectivity profiles^59^, small surface patches assigned to a given functional network ^58^, or network variants (that is, locations where functional connectivity often parts away from the group average) ^60^.

As a second core criterion for robustness, we quantified the percentage of variance explained by average FD. Although we specifically focused our analyses on the 100 subjects moving the least within the HCP dataset, and performed stringent preprocessing, there remained clear patterned influences of head movement on all features types (**Supplementary Figure 14**). Thus, one cannot rule out the possibility that head movement partly alters the structural and/or fMRI data, leading to artificially inflated fingerprinting performance. However, selecting parameter combinations that minimize the extent of the association enables to limit potential impacts as much as possible.

Interestingly, robustness to acquisition settings and sensitivity to head movement were largely uncorrelated (see **Figure 5J-K**), which justifies their parallel consideration. Regarding the main impacts of factors of variation, z-scoring consistently yielded higher robustness when applied, while spectral normalization impacted spectral, but not regional features. This can be explained by the fact that in the former case, it is the second to last step leading to the obtention of feature coefficients, while in the latter case, the spectral data is converted back into regional contributions with dedicated filtering schemes, which likely down-weights the importance of normalization on the final coefficients.

Our robustness results can be contemplated from two different perspectives. On the one hand, one may wish to select a feature type which minimizes sensitivity to external factors *when the best parameters are used*. If this is the goal, our investigations revealed that liberality was the best feature type overall (as it was tied to smaller *P*_*I*_), followed by PSD and alignment, then SDI (larger *P*_*I*_), and eventually energy for which it was not possible to converge on consensus parameters that would jointly optimize with respect to all criteria. All feature types yielded similar robustness to head movement overall, but with varying impacts of acquisition settings. On the other hand, one may wonder which feature type performs best overall, *without having specified exact parameters*. In this case, our comparative analysis (**Figure 9A**) suggests that for robustness to acquisition settings, PSD and liberality win, while sensitivity to head movement is lower for SDI and alignment.

While stability across sessions is desirable, it does not imply cross-subject discriminability. To address the latter, we quantified the ICC (**Figure 6**). The main conclusion from this analysis was the fact that while *Structural individuality* was the least influential factor of variation overall, it largely dominated in terms of modulating ICC values: when a group-wise SC was used for GSP analysis, only a few coefficients at best reached excellent values, but when using a subject-specific SC instead, in the majority of cases, almost all coefficients became strongly discriminative. It followed that when relying on subject-specific SCs, fingerprinting could yield perfect accuracy for all feature types, with very little remaining influence of other factors of variation. Several intermingled effects can explain these observations. First, when eigenmodes are separately extracted for each subject, even if a match across subjects exists, they may not necessarily be ordered similarly as it depends on the associated eigenvalues. This effect is negligible for smallest frequency eigenmodes, but matters a lot for higher frequency ones. In line with the above, the feature type that was the most impacted by *Structural individuality* was liberality, which specifically considers the fraction of regional time courses associated to the highest frequency eigenmodes. Second, although differences in feature coefficient values or feature vector patterns as a function of structural individuality remain mild compared to those induced by other factors of variation, they are nonetheless significant (see **Table 1**). As such, it is not surprising that they can induce such a large impact. Conceptually speaking, it simply means that when individual structural *and* functional data are included, fingerprinting becomes easier compared to the use of subject-specific functional, but group-averaged structural information; *ergo*, this is a demonstration of the superiority of multimodal fingerprinting ^95^.

While this is an interesting finding in itself, subject-specific SCs cannot be used in all settings. For example, if one aims at behavioural prediction, the functional data needs to be expressed as a function of the same underlying structural basis, so that feature coefficients can be compared, across subjects, in a regression setting. For this reason, we further analyzed our results focusing on the use of a group-wise SC. PSD and alignment then appeared to be the most discriminative approaches overall, ICC-wise, when focusing on the coefficients used for fingerprinting (**Figure 9A**, top right panel). Furthermore, we observed that fingerprinting accuracy could reach 100% for PSD, alignment and SDI with the right parameter combinations, but all feature types exhibited different properties: for PSD, there was never more than 50% used features for fingerprinting, while the range was much broader for SDI; in other words, parsimony of the feature set was more pronounced in the former case. For alignment (as well as for liberality), almost all features had to be used for optimal fingerprinting in many cases. While a more parsimonious set of insightful features may not play an important role in the analyses that we present therein, it could contribute to prevent overfitting if more sophisticated machine learning pipelines were implemented for fingerprinting in future work ^96^. The reliance on only a subset of coefficients likely implies that only a fraction of eigenmodes contain individual-related information.

Interestingly, the assessment of fingerprinting also revealed the importance of the scale of the atlas used for the analyses: while S3^-^ results were similar to S3 ones, the use of S1 data yielded a maximal accuracy of only 75% (PSD) to 90% (SDI). This implies that a minimal amount of granularity in the used atlas is required to fully exploit GSP analysis. In addition, when considering the change in accuracy upon removal of the subcortex/cerebellum or a switch to S1, PSD and liberality were the worst approaches overall.

All in all, when combining all quality criteria, PSD and SDI emerged as the most all-around choices for optimal fingerprinting. It is interesting that these two feature types, while very different from each other, are nonetheless related in that the cutoff frequency used in SDI quantification is based on the power spectrum. The relevance of PSD as a well-performing feature type in fingerprinting hints at the fact that there is more information in power spectral density than a mere subdivision between low and high frequencies. It also opens up interesting possibilities for future work: one would be the combination of PSD and SDI features to determine whether outcomes of interest improve, while another would be the improvement of the current SDI approach, so that it can incorporate finer details regarding the power spectrum.

### Limitations and perspectives

There are several limitations to our study. First, largely owing to computational feasibility, we focused our analyses on only a subset of 100 subjects from the HCP. In future work, it will be important to examine how the ability to fingerprinting evolves when a larger set of subjects is considered. In a past study, fingerprinting accuracy dropped as the number of included subjects increased, as further confirmed by a parametric modelling approach ^63^. It could be interesting to examine whether some feature types are more sensitive to this effect than others, and how much parsimony in the feature set impacts this trend. Furthermore, another possibility stemming from the same work could be the quantification of *confidence* (*e*.*g*., number of cases for which session matching is sure beyond 99.9%), as a criterion more amenable to clinical applications ^63^.

Second, our population of subjects included only particularly low movers. While this enabled us to reproduce past results obtained on a set of average movers ^38^, and to explore the impacts of head movement in an optimal set of subjects, it also leaves open the possibility that we only considered a behaviourally specific type of individuals (*i*.*e*., those conforming to the guidelines not to move inside the scanner) in our analyses. Third, the recordings from the HCP are not reflective, in terms of their parameters, of typical clinical datasets.

Fourth, while we attempted to be as comprehensive as possible, several further branches within GSP analysis were left out and should be considered in future work. In particular, in the present study, we systematically factored out the temporal dimension by computing the L2-norm across time, thus avoiding the analysis of temporal dynamics. However, recent work shows that there are structured fluctuations in the aligned and liberal components of dynamic functional connectivity across time ^97^. In the context of fingerprinting with EEG recordings, properties reflective of the temporal dynamics of transcranial magnetic stimulation-evoked potentials also enabled fingerprinting when probed across several brain locations, further evidencing the interest of time-resolved properties in individuality^98^.

Another potentially interesting direction could be to steer GSP analysis towards specific subtypes of connections involving dedicated sub-networks while deriving the eigenmodes. This can be achieved through the use of Slepian vectors^99^, which are essentially eigenmodes with an added constraint of energy localization within a user-defined set of nodes. At the level of resting-state functional connectivity, it has been repeatedly shown that focusing on specific sets of connections– particularly within the frontoparietal, default mode and dorsal attention networks^59,61,64^–can boost fingerprinting performance. It will be interesting to address whether similar observations also hold for GSP analysis.

Fifth, we only considered resting-state data, without relying on the battery of available taskbased recordings provided by the HCP. Maximum differentiability across subjects has been shown to be achieved when resting-state recordings are jointly analyzed with at least one task recordings ^100^; thus, it could be that our ranking of GSP features changes when broadening the set of analyzed input time courses.

Sixth, while we examined the ability to fingerprinting subjects, we did not dive into the arguably more directly relevant question of behavioural prediction. At the level of resting-state functional connectivity, the underlying correlates of both have been shown to differ; in fact, while the set of features enabling optimal fingerprinting shows a large variability across subjects, the same does not hold for features contributing to behavioural prediction^65^. Thus, there is no guarantee that the approaches that we have deemed best for fingerprinting will fare as well for behavioural prediction, and a new comparative analysis will likely be required to answer this question.

Seventh, although we touched upon the question of the optimal scale regarding the parcellation to use in GSP analyses, we have limited ourselves to a regional viewpoint, while voxel-wise investigations are also possible.

Turning to future perspectives, we have contributed a principled comparative approach which can be extended to compare not only GSP variants, but also other types of analytical approaches, on the same outcome measure(s) of interest. To exemplify this, we also illustrated the results of our methodology at the level of graph metrics inferred from FCs, so that we would handle data with a dimensionality similar to that of GSP feature vectors. Note that fingerprinting using graph metrics rather than functional connectivity values has been shown effective in past work ^66^. We saw that while reasonable performance can be achieved in such a setting (up to 80% fingerprinting accuracy), together with low sensitivity to atlas specificity and head movement, the overall quality profile of graph metrics could not rival those from the best GSP approaches, further speaking in favour of the use of multimodal analytical approaches.

GSP analysis is particularly remarkable in its versatile nature. On top of the many ways it offers to combine structural and functional MRI brain data as explored therein, it can also be seamlessly applied to EEG or MEG data ^101–104^, or to a voxel-wise viewpoint, for example by using a structural graph to smooth non-isotropically ^105^ or by interpolating activity in the white matter ^106^. It has merits in modelling as well ^107^, and enables many more possibilities than the ones discussed here, such as the comparison between the eigenmodes obtained from structural and functional graphs by projection ^108^. Also, note that from aligned or liberal time courses, one can also append conventional analytical steps, such as the extraction of an FC ^109^ or of dynamic functional connectivity states ^110^.

## Conclusion

In this work, we conducted a large-scale comparative analysis of graph signal processing pipeline variants to establish optimal settings in the context of fingerprinting. We showed that all studied factors of variation played major roles in shaping GSP features, in ways that were feature-specific and involved interactions. We ranked competing approaches in terms of a set of quality criteria, and pinpointed power spectral density and the structural decoupling index as optimal approaches. We hope that our work will demonstrate the versatility of GSP analysis, while helping the community to refine their investigations by more informed choices.

## Acknowledgments and data availability

Data were provided by the Human Connectome Project, MGH-USC Consortium (Principal Investigators: Bruce R. Rosen, Arthur W. Toga and Van Wedeen; U01MH093765) funded by the NIH Blueprint Initiative for Neuroscience Research grant; the National Institutes of Health grant P41EB015896; and the Instrumentation Grants S10RR023043, 1S10RR023401, 1S10RR019307.

## Acknowledgements

Data were provided by the Human Connectome Project, WU-Minn Consortium (Principal Investigators: David Van Essen and Kamil Ugurbil; 1U54MH091657) funded by the 16 NIH Institutes and Centers that support the NIH Blueprint for Neuroscience Research; and by the McDonnell Center for Systems Neuroscience at Washington University. This work has been supported by Swiss National Science Foundation grant #197787.

## Author contributions

T.B. and P.H. designed the study. M.S. processed the resting-state functional MRI data. J.P. processed the diffusion-weighted MRI data. H.F. contributed to the determination of the explored factors of variation. Y.A. devised the atlas used in the study. T.B. performed all analyses and wrote the manuscript, which was reread by all authors.

## Footnotes

1 Arash Salarian (2023). Intraclass Correlation Coefficient (ICC) (https://www.mathworks.com/matlabcentral/fileexchange/22099-intraclass-correlation-coefficient-icc), MATLAB Central File Exchange. Retrieved October 18, 2023.

2 Moses (2023). spider plot (https://github.com/NewGuy012/spider_plot/releases/tag/20.4), GitHub. Retrieved October 18, 2023.

