## Supplementary Materials for "Large-scale comparative analysis reveals top graph signal processing features for subject identification"

### Supplementary Results

In what follows, we provide supplementary visualizations of our results, explain how they complement our main results, and briefly comment on the most salient features that transpire from them.

*Impacts of factors of variation on pattern similarity*

In **Supplementary Figures 1-3** (panels A to E), we provide visualizations of feature vectors for each feature type (PSD, energy, alignment, liberality and SDI), across feature index (from left to right) and parameter combinations (top to bottom), respectively for S3, S3$^-$ and S1 input data. To ease visualization, the feature vector data was first averaged across scans, then across subjects, and eventually z-scored. Note that for alignment/liberality, we selected only one indicative set of coefficients ($n_H$ = 22, 18 or 12 for S3, S3$^-$ and S1, respectively), while for SDI, we focused on the extraction of a cutoff frequency through subject-wise mean PSD.

**Supplementary Figure 1** complements the main results shown in **Figure 2**, as the low-dimensional spaces outlined in the latter figure were constructed based on distances between individual rows of each heatmap. For PSD and energy, it can clearly be observed that the maximal values when using the modularity matrix lie in intermediate coefficient indices, contrasting with all other operator subtypes. It can also be seen that patterns are smoother when considering subject-specific as opposed to group-wise SCs, and more spread when z-scoring is not performed. The strong impact of *Property* in the case of liberality is also evident, particularly in the NF(D) *versus* MFL/MFA dissociation. Furthermore, for alignment and SDI, the use of the modularity matrix yields patterns that are anti-correlated to those obtained with other operator subtypes. Overall similar observations also apply to **Supplementary Figures 2-3**, which shows that our observations regarding feature vector patterns are robust to parcellation specificities.

For spectral domain (**Supplementary Figure 4**) and regional domain (**Supplementary Figure 5**) features, we provide $n_p \times n_p$ matrices summarizing the mean and the standard deviation of similarity matrices across subjects, where $n_p = 128$ is the total set of parameter combinations at hand and the same specific cases as above were selected for alignment, liberality and SDI. Again, these figures provide complementary information to that presented in **Figure 2**: first, Pearson's correlation values are provided, which can sometimes differ from the similarity values used to construct **Figure 2** (normalized for each parameter combination; see **Materials and methods**). Second, they provide insight into differences across subjects (through the standard deviation) on top of considering averages across subjects.

Fitting with the observations made from **Supplementary Figure 1**, the use of the modularity matrix yields particularly different outputs compared to that of other operator subtypes for PSD (**Supplementary Figure 4A**), and patterns anti-correlated to the others for alignment and SDI (**Supplementary Figure 5A,C**). PSD, energy, alignment and SDI are all impacted by the removal of subcortical and cerebellar structures (compare the top left and top middle matrices in each case), while the use of S1 data most strongly affects liberality outcomes (compare the top left and top right matrices). On the whole, energy and SDI are associated to the largest variability of similarity patterns across subjects.

**Supplementary Figure 6** provides complementary low-dimensional representations of the data akin to those presented in **Figure 2**. For PSD, the absence of subcortical and cerebellar structures (input data type $S3^-$) makes the distinction between modularity matrix and other operator subtypes much more influential compared to other factors (**Supplementary Figure 6A**, top middle plot), which fits with the corresponding data in **Supplementary Figure 4A**. For alignment, regardless of input data type, it can be seen that data points associated to the use of the smallest probed number of retained coefficients are particularly distant from the others (**Supplementary Figure 6B**, top row): this highlights that alignment outputs tend to converge when a sufficient number of coefficients is included, and then stabilize. Note that this does not apply to liberality. These observations are well in line with the results from **Figure 3**, where *Coefficients* was more impactful for alignment than for liberality.

*Impacts of factors of variation on feature values*

The Bonferroni-corrected $p$-values associated to the mixed-effects model results presented in **Figure 3A** are displayed in **Supplementary Figure 7**, while **Supplementary Figures 8-9** summarize, respectively, the F-statistics and $p$-values obtained for S3- and S1 input data. **Supplementary Figure 7** shows that, as outlined in the main results, the large majority of individual feature coefficients showed a significant impact of all investigated fixed effects (factors of variation and their interactions). **Supplementary Figures 8-9** show that the observations made for S3 input data, in terms of F-statistic patterns and impacts of individual factors of variation/interactions on the feature types, also hold for S3- and S1 input data. This is further complemented by **Supplementary Tables 1-2**, which summarize the actual F-statistics at hand akin to **Table 1**.

*Random effects*

The random effects associated to S3- and S1 input data are shown in **Supplementary Figure 10A-B**. Overall, the observations made from **Figure 4** also apply to these cases: indeed, PSD and alignment show the most stable random effect values across coefficients.

*Robustness to acquisition settings and head movement*

**Supplementary Figure 11** provides displays of average similarity across sessions acquired on the same day (X-axis) as opposed to across days with similar phase encoding direction (Y-axis) for all the considered feature types. These displays can thus be compared to **Figure 5A**, where average data across subjects was shown (alongside individual subject-wise data points) for two selected SDI subcases. If a data point lies above the main diagonal, it means that similarity between sessions acquired on two separate days with similar acquisition parameters was larger than similarity between sessions acquired on the same day with distinct acquisition parameters, which highlights an unwanted modulation of feature vectors by acquisition settings. Across feature types, this primarily occurs when temporal z-scoring is not performed. It can also be appreciated how a broad range of similarity values is taken across parameter combinations, with optima that differ as a function of feature type.

**Supplementary Figures 12/13** display similar representations for S3$^-$ and S1 input data **Supplementary Figures 12/13A**, respectively, complemented by associated visualizations in terms of same-day sessions' similarity *versus* penalty (**Supplementary Figures 12/13B**). In terms of similarity values, similar observations as above can be made regardless of parcellation specificities. About quality criteria, however, there is a clear impact of input data type: for example, the worst parameter combinations are completely different for alignment between S3$^-$ and S1 cases. This shows the importance, in extracting optimal GSP parameter combinations, of not just considering robustness to external factors, but also generalizability in terms of parcellation specificities as we did in **Figure 9**.

**Supplementary Figure 14** summarizes the average associations between mean framewise displacement and feature coefficients across scans for S3, S3$^-$ and S1 input data. **Supplementary Figure 15** plots the percentage of incongruent cases $P_I$ against the mean percentage of explained variance by mean framewise displacement $P_{\text{FD}}$ for a representation of our two final robustness quality criteria in the case of S3$^-$ and S1 input data. The main observation from **Supplementary Figure 14** is the presence of associations between feature coefficient values and mean framewise displacement, which are structured across coefficients in a way that differs across feature types and depending on the investigated parameter combination. Recall that this is the case despite specifically considering the 100 subjects with the smallest head displacement from within the HCP dataset, preprocessed according to state-of-the-art guidelines. In our view, this highlights the fact that even when dealt with to the best of one's abilities, head movement remains a problematic confound that must be adequately addressed. As for quality metrics (**Supplementary Figure 15**), the observations made from **Figure 5** (absence of outliers in terms of $P_{\text{FD}}$ for SDI, but larger minimum in terms of $P_I$) generalize to S3$^-$ and S1 input data.

*Fingerprinting ability*

**Supplementary Figures 16/17** display ICC values for S3$^-$ and S1 input data cases, alongside representations of average ICC values across coefficients plotted against same-day sessions' similarity. Overall, the key observations made from **Figure 6** also apply to these other input data cases: first, the use of subject-specific SCs considerably increases ICC values across coefficients, and second, parameter combinations characterized by identical similarity between same-day sessions often takes distinct mean ICC values.

In **Supplementary Figure 18**, we see an overview of how fingerprinting accuracy evolves as a function of the fraction of feature coefficients used for fingerprinting. Beyond the fact that different parameter combinations yield distinct accuracies, it is particularly interesting to notice the different behaviours across feature types: for alignment and liberality, accuracy often reaches its optimum only when close to all features are used. For SDI, high accuracies can be reached when using fewer feature coefficients already, but performance remains high when more are incorporated. For PSD, however, there is only a specific amount of coefficients that yield optimal fingerprinting, past which performance degrades. These observations are in line with those made from **Figure 8**, where a limited set of coefficients was shown to be optimal for fingerprinting with PSD as opposed to other feature types.

while **Supplementary Figure 19** shows, for scale $3^-$ and S1 input data, for which percentage of feature coefficients the best fingerprinting accuracy was achieved across feature types and parameter combinations, similarly to **Figure 8E-F** in the main results. The results for S$3^-$ input data are similar to the main ones: PSD and SDI both yield 100% accuracy in many cases, few coefficients are required for optimal fingerprinting using PSD, alignment and liberality work optimally when considering all coefficients, and the fraction of used coefficients for SDI can vary from case to case. For S1 input data, however, there are noticeable differences: first, perfect fingerprinting accuracy is never reached for any feature type, which highlights the fact that the available information is then too little (*i.e.*, not enough parcels available) for good fingerprinting. Second, even for PSD, larger percentages of features must be used for performance to be optimized. These observations can also be drawn from **Figure 8I** compared to **Figure 8G-H**.

*Comparison of candidate feature types and parameter combinations*

**Supplementary Figures 20/21** summarize quality criteria for the S$3^-$ and S1 input data cases, respectively. They are thus the equivalent of **Figure 9** from the main results. For S$3^-$ input data (**Supplementary Figure 20**), individual quality criteria behave similarly to the S3 case, and PSD and SDI still emerge as the top feature types. For S1 input data (**Supplementary**

**Figure 21**), however, the picture partly changes: alignment becomes more robust to acquisition settings, while SDI worsens; alignment also shows the best accuracy outcomes for top parameter combinations, and improves in generalizability while PSD worsens. Consequently, for specific subcases, alignment now competes with PSD and SDI, both of which are particularly impacted in terms of robustness. This highlights the need for sufficient areas in the used parcellation if PSD or SDI are to be used as feature types of interest.

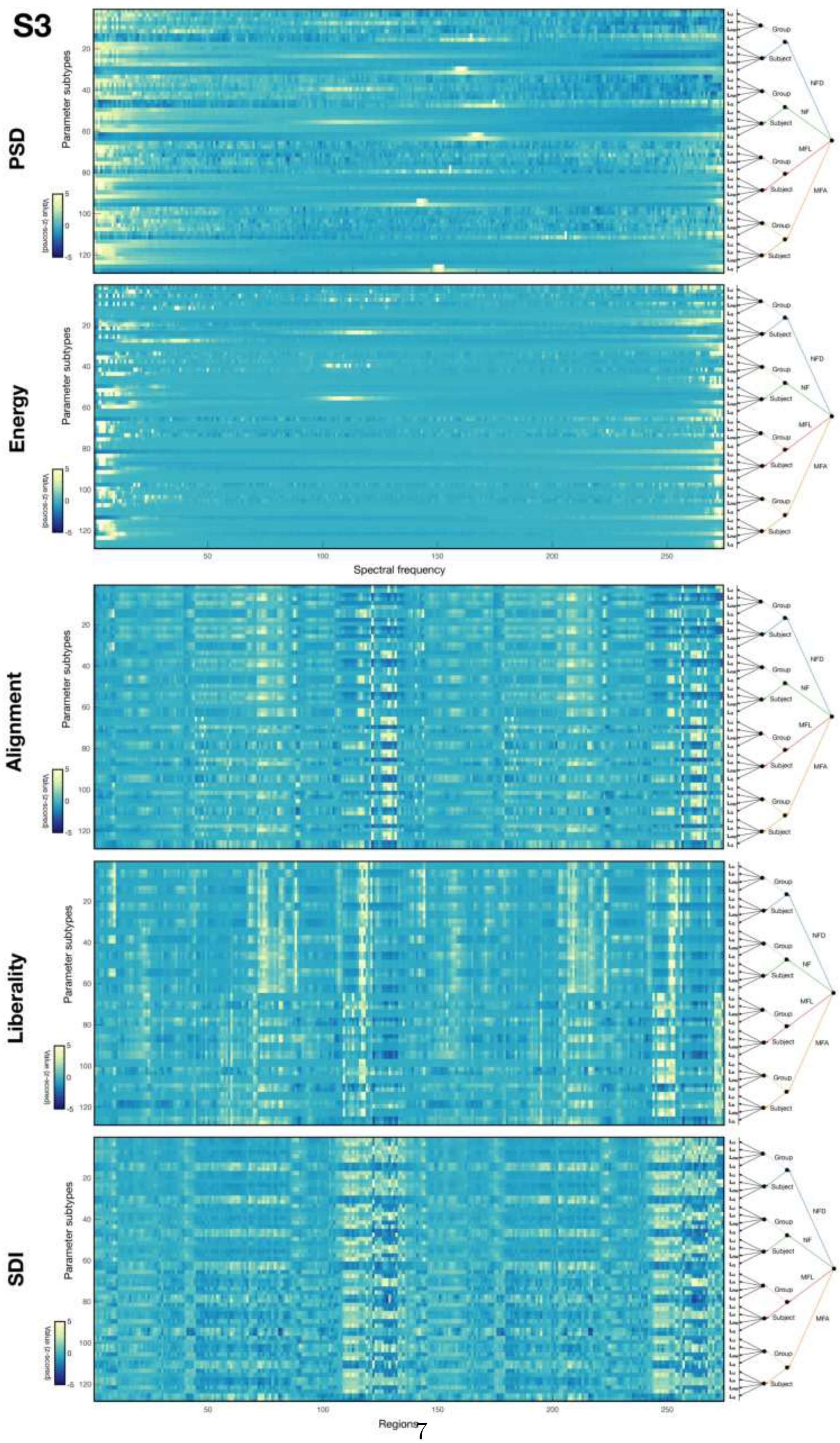




Figure 1: Average feature vector patterns (across scans, then subjects) across parameter combinations for all five analyzed feature types on scale 3 (S3) data. On the right side of each heatmap, a tree representation summarizes the key parameter values associated to each row of the display. Note that for spectral domain features (PSD and energy), the X-axis denotes spectral frequency index, while for regional domain features (alignment, liberality and SDI), it instead represents region index. The data from each row is z-scored to enable the comparison of patterns rather than original values.

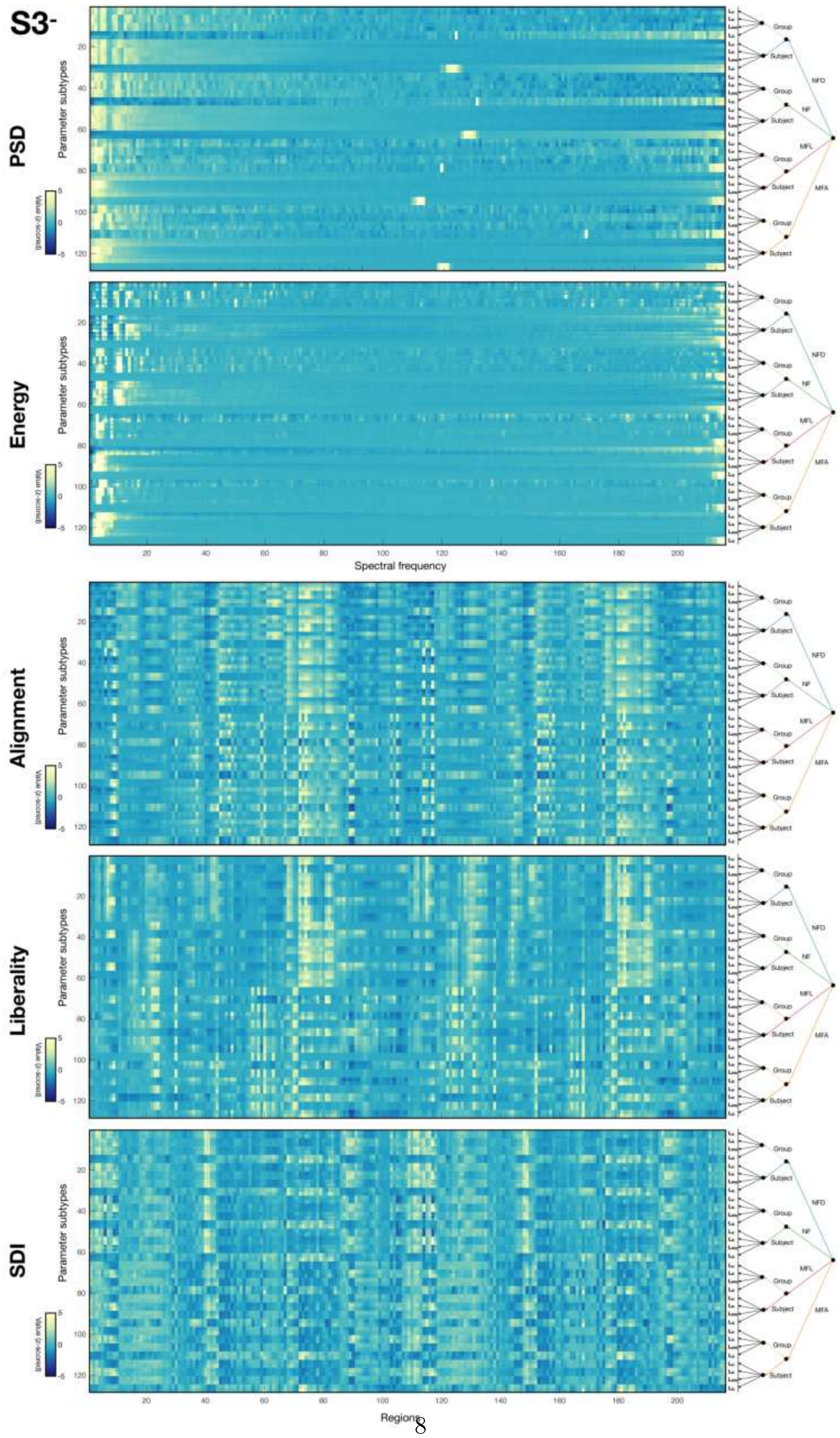




Figure 2: Average feature vector patterns (across scans, then subjects) across parameter combinations for all five analyzed feature types on scale 3 data without cerebellum and subcortex (S3⁻). On the right side of each heatmap, a tree representation summarizes the key parameter values associated to each row of the display. Note that for spectral domain features (PSD and energy), the X-axis denotes spectral frequency index, while for regional domain features (alignment, liberality and SDI), it instead represents region index. The data from each row is z-scored to enable the comparison of patterns rather than original values.

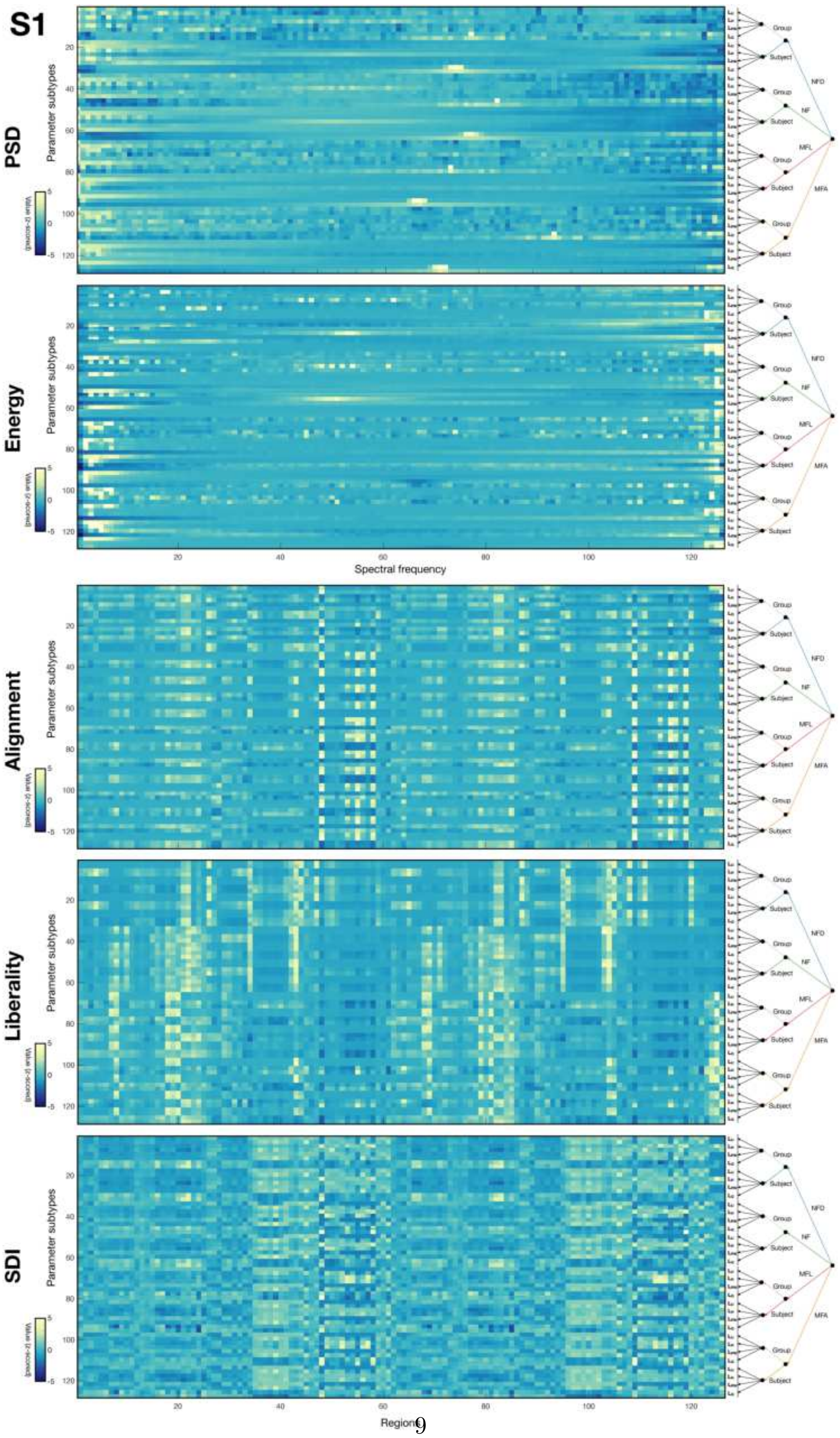




Figure 3: Average feature vector patterns (across scans, then subjects) across parameter combinations for all five analyzed feature types on scale 1 (S1) data. On the right side of each heatmap, a tree representation summarizes the key parameter values associated to each row of the display. Note that for spectral domain features (PSD and energy), the X-axis denotes spectral frequency index, while for regional domain features (alignment, liberality and SDI), it instead represents region index. The data from each row is z-scored to enable the comparison of patterns rather than original values.

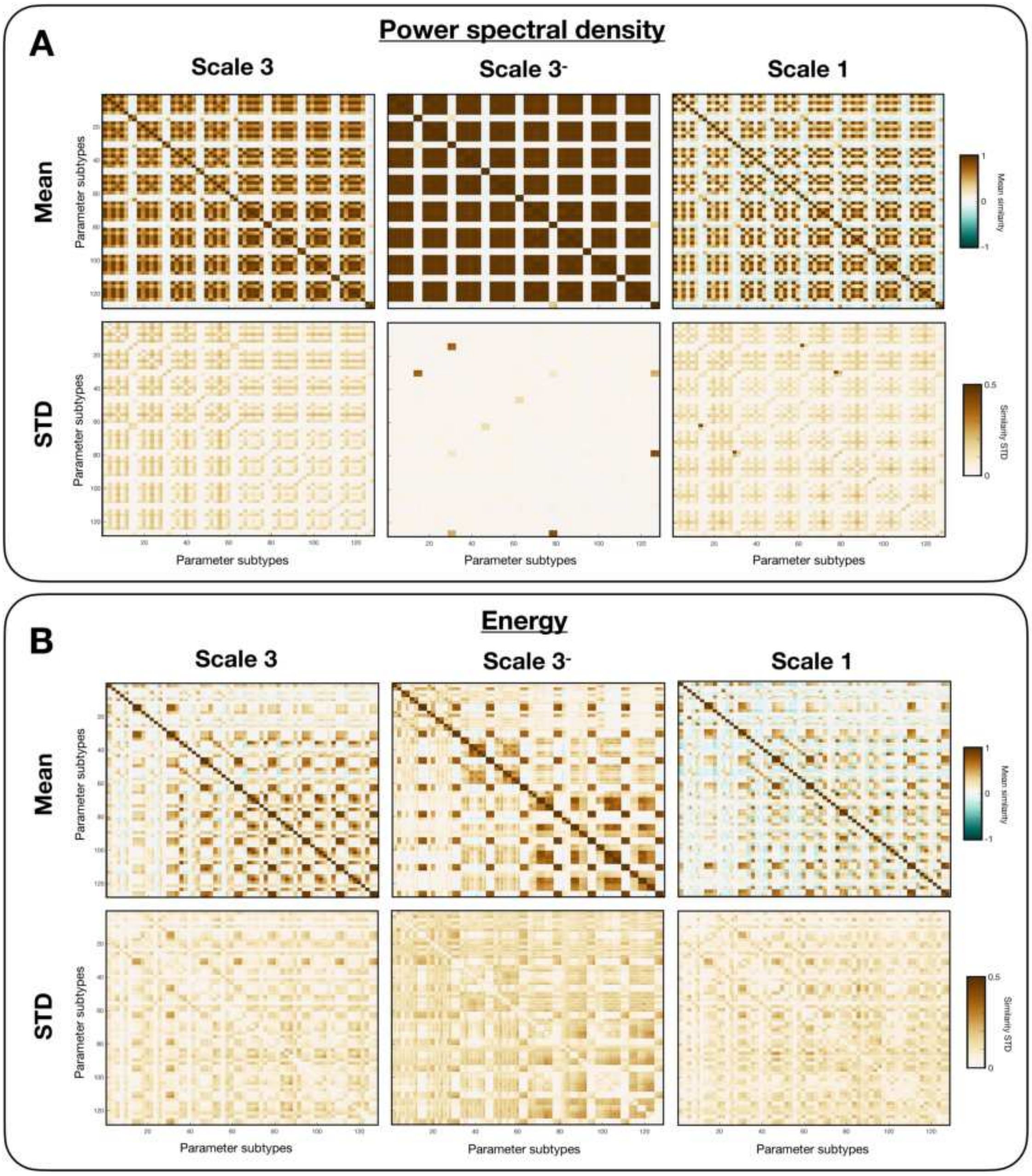


Figure 4: For PSD **(A)** and energy **(B)**, average (top row) and standard deviation (bottom row) of similarity values across subjects when considering scale 3 data (left column), scale 3 data without cerebellum and subcortex (middle column), or scale 1 data (right column).

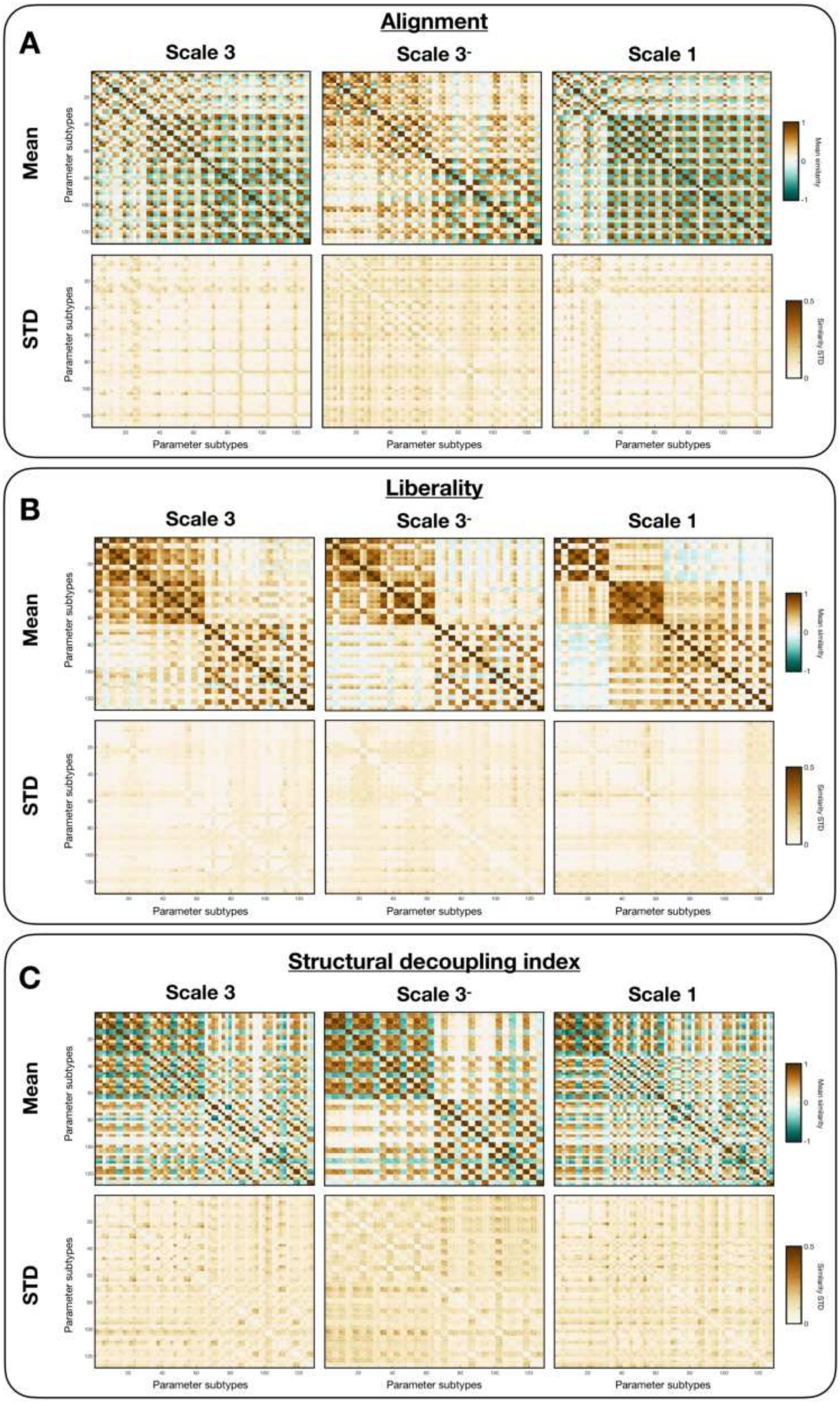


Figure 5: For alignment **(A)**, liberality **(B)** and SDI **(C)**, average (top row) and standard deviation (bottom row) of similarity values across subjects when considering scale 3 data (left column), scale 3 data without cerebellum and subcortex (middle column), or scale 1 data (right column).

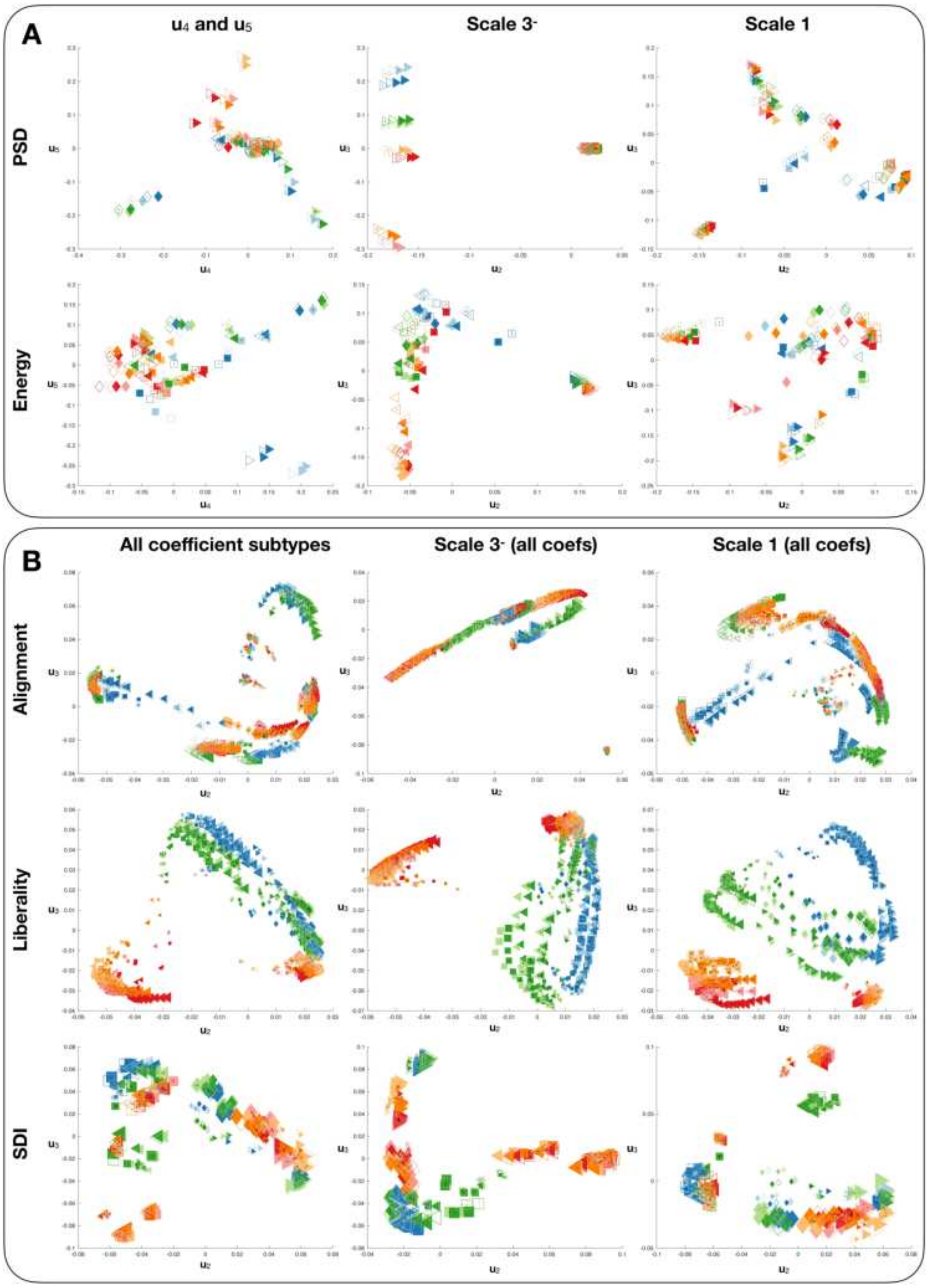


Figure 6: For spectral domain features **(A)** and regional domain features **(B)**, complementary low-dimensional visualizations of similarity between feature vectors. For spectral domain features, we display the third and fourth discriminating dimensions (left column), and the results for S3⁻ or S1 input data (middle or right column). For regional domain features, in the left column, we show the results of the main S3 analysis when all candidate numbers of coefficients are included (for alignment or liberality; seen as data points of increasing size for larger numbers of retained coefficients), or when the four ways to select a cutoff frequency are considered (for SDI; also shown as data points of increasing size for 50/50 selection, time point-specific selection, mode-based selection and subject-wise mean PSD selection). In the middle and right columns, similar displays are shown when considering S3⁻ or S1 input data instead.

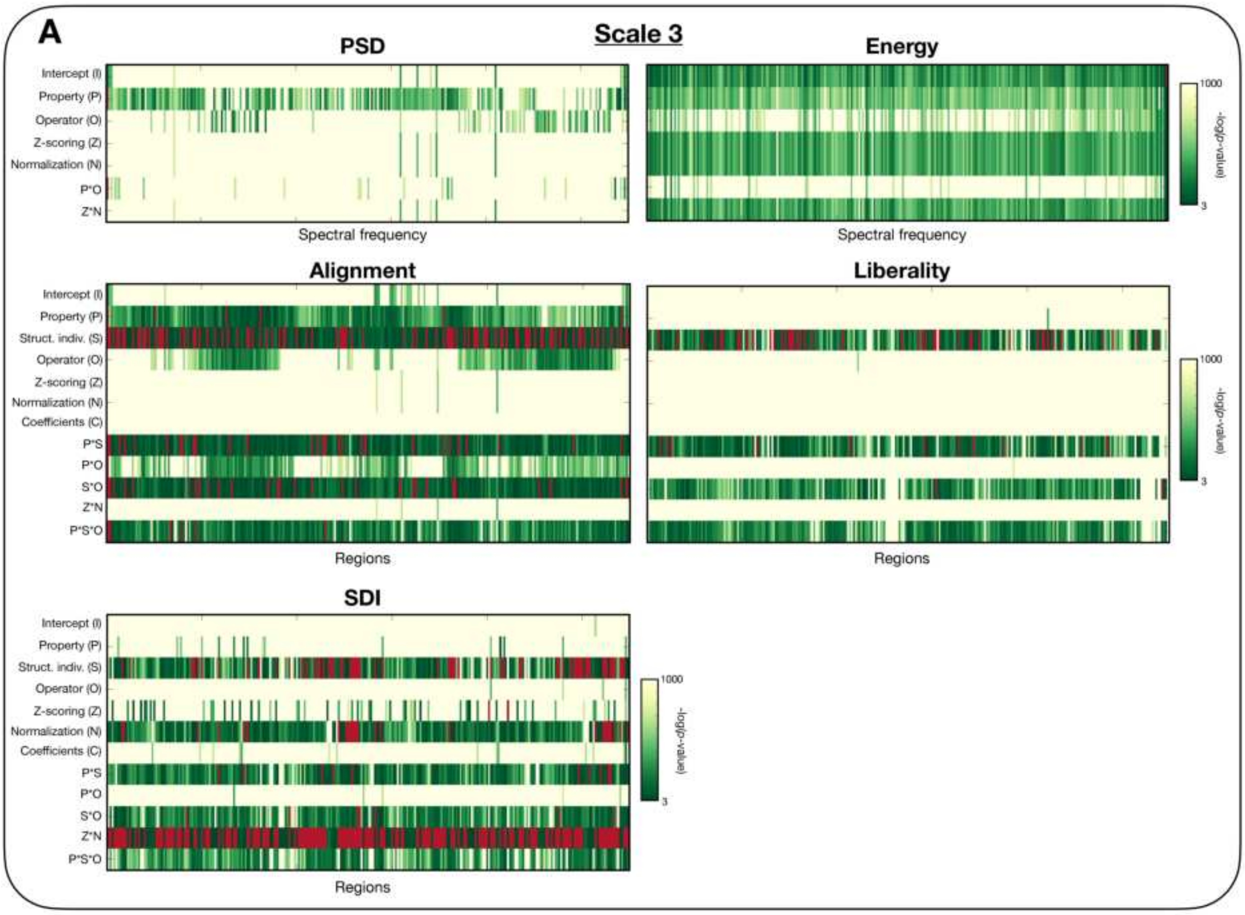


Figure 7: For scale 3 data and all considered feature types, Bonferroni-corrected $p$-values across feature coefficients (left to right) and modelled fixed effects (top to bottom). Note that $p$-values are shown as $-\log(p)$, so that the higher the value, the more significant the outcome. Significance is achieved when $p > 3$. Non-significant outcomes are displayed in red.

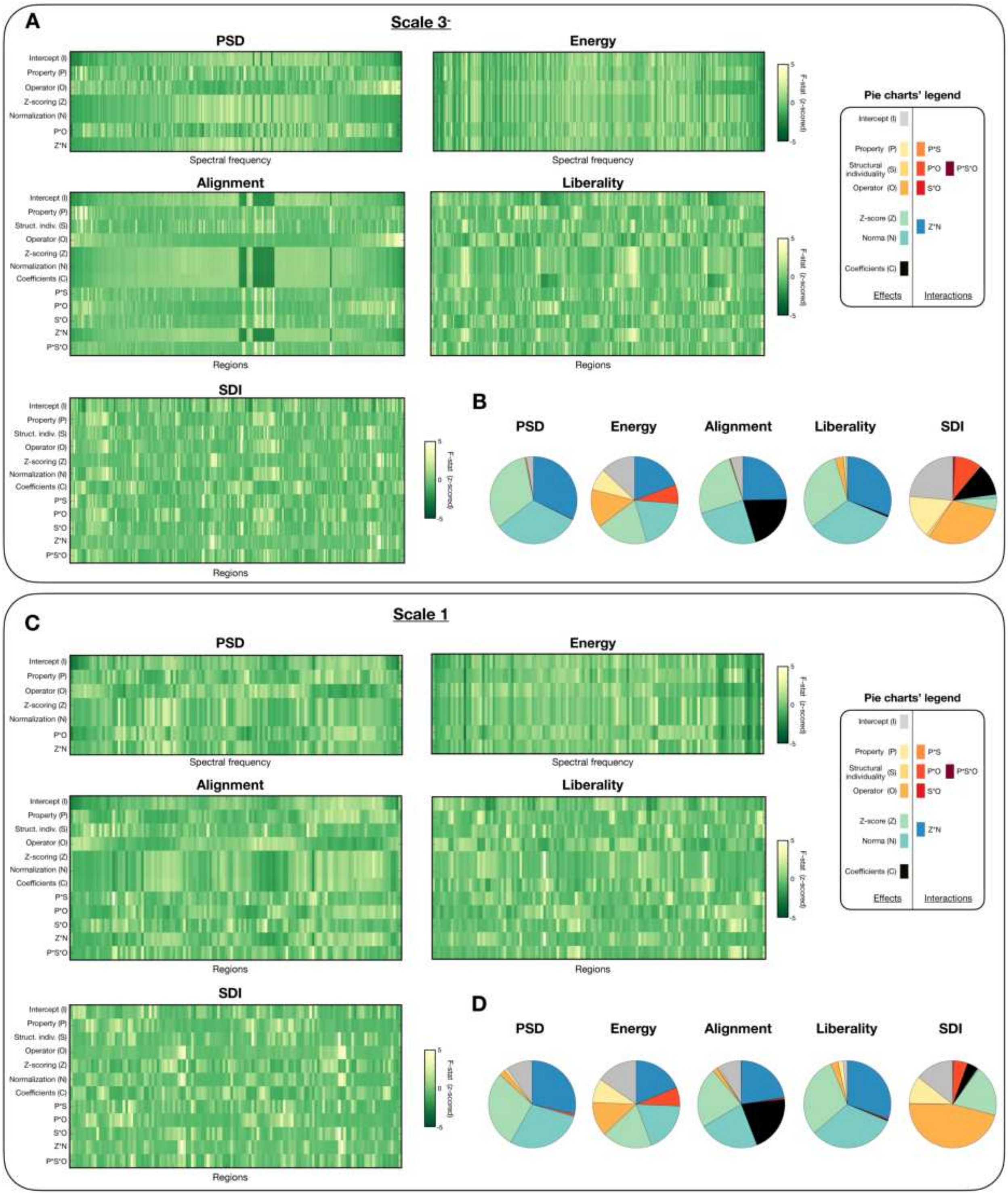


Figure 8: For S3- **(A)** or S1 **(C)** input data, across all considered feature types, F-statistics across feature coefficients (left to right) and modelled fixed effects (top to bottom). Note that the displayed F-statistics have been z-scored for each row to enable the visualization of the pattern across feature coefficients; for a summary of actual values, see **Supplementary Tables 1-2**. Pie chart representations of the respective contributions of each factor of variation to a given feature type are also shown for S3- **(B)** or S1 **(D)** input data.

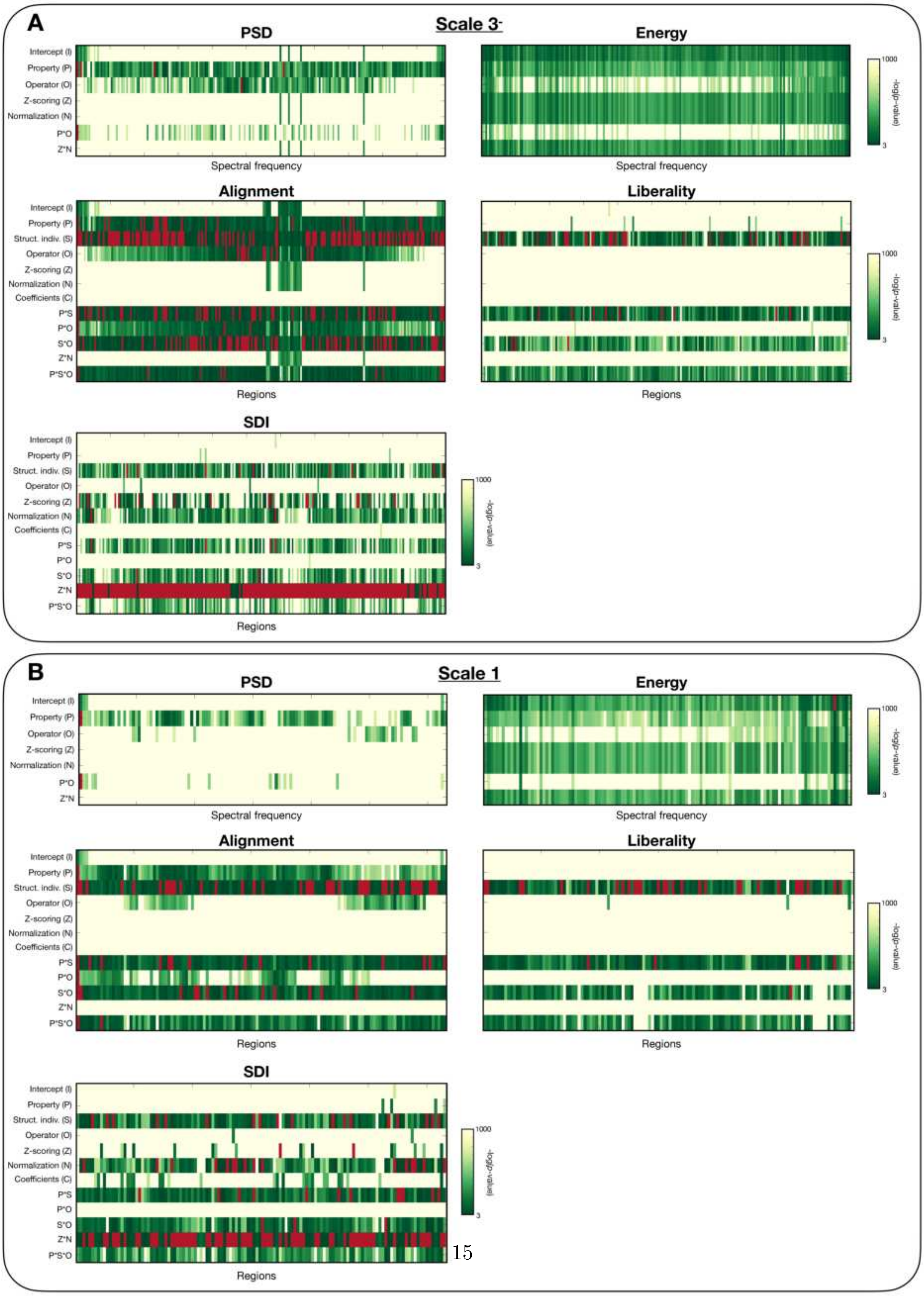


Figure 9: For S3- **(A)** or S1 **(B)** input data, across all considered feature types, Bonferroni-corrected $p$-values across feature coefficients (left to right) and modelled fixed effects (top to bottom). Note that $p$-values are shown as $-\log(p)$, so that the higher the value, the more significant the outcome. Significance is achieved when $p > 3$. Non-significant outcomes are displayed in red.

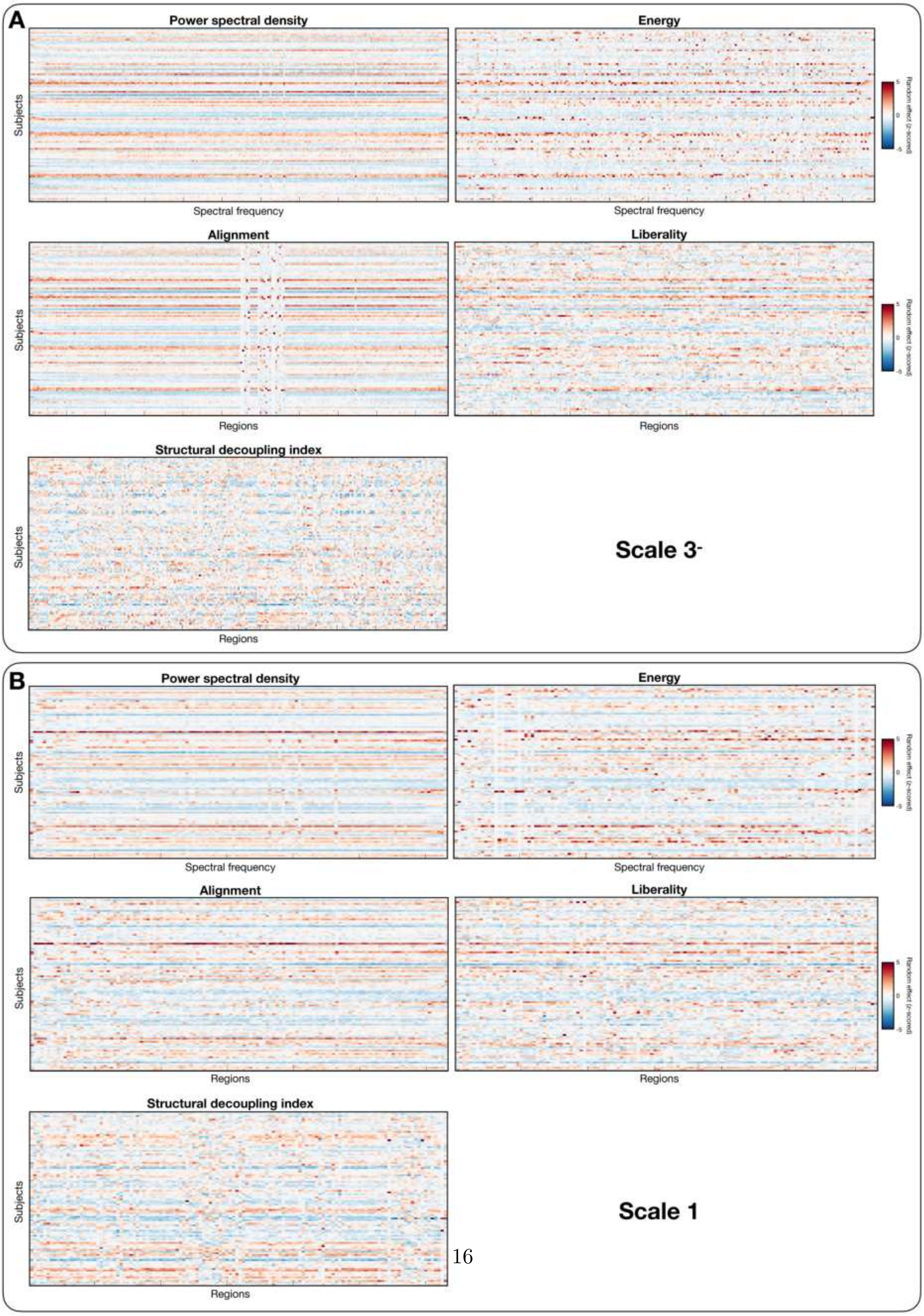




Figure 10: For S3- **(A)** or S1 **(B)** input data, summary of random effects across all considered feature types. Note that the data was z-scored for each column to enable a comparison of the underlying patterns across feature coefficients regardless of actual values.

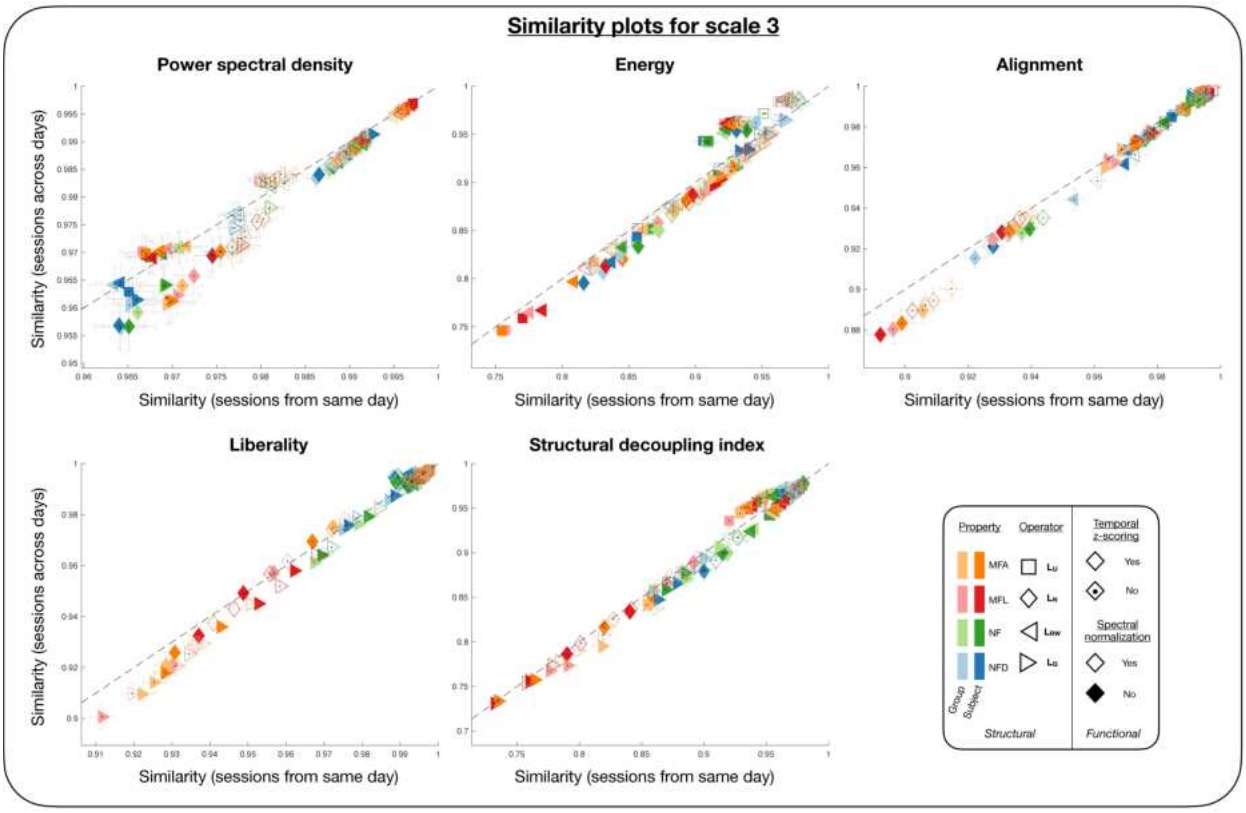


Figure 11: For all five feature types of interest, similarity between both scans acquired on the same day (X-axis) is contrasted to similarity between both scans acquired on different days, but with similar phase encoding direction (Y-axis). The similarity data was originally averaged across both available pairs of sessions in each case (*e.g.*, day-1 scans and day-2 scans). The error bars denote standard error of the mean across subjects. Each data point is associated to one specific set of parameters as described in the bottom right legend.

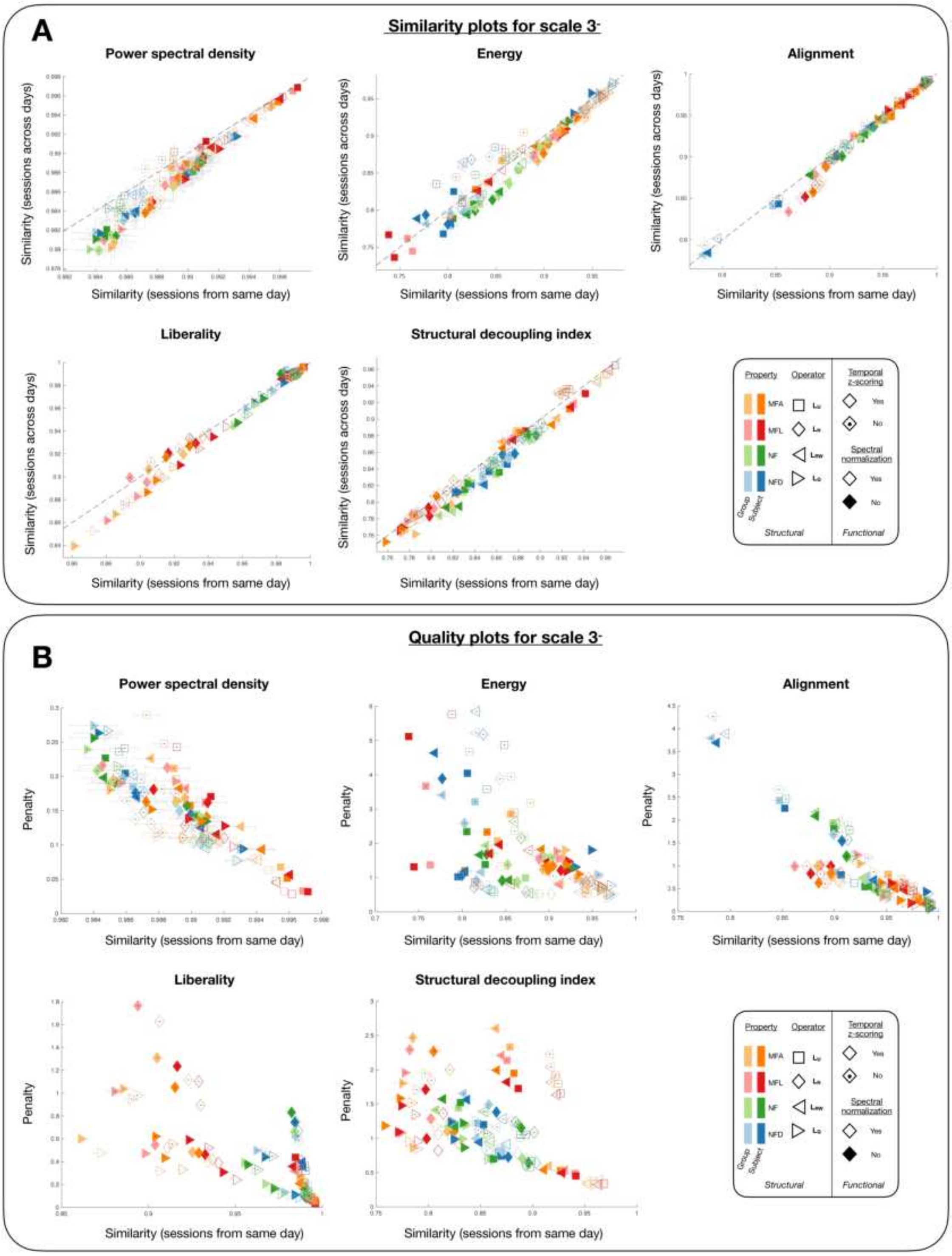


Figure 12: **(A)** On S3- input data, for the five feature types of interest, similarity between both scans acquired on the same day (X-axis) is contrasted to similarity between both scans acquired on different days, but with similar phase encoding direction (Y-axis). The similarity data was originally averaged across both available pairs of sessions in each case (*e.g.*, day-1 scans and day-2 scans). The error bars denote standard error of the mean across subjects. Each data point is associated to one specific set of parameters as described in the bottom right legend. **(B)** For the same five feature types, similarity between same-day scans (X-axis) is also contrasted to the devised penalty measure $\Omega$ (Y-axis), where a larger penalty value highlights that there were more subjects with larger similarity between different-day scans than between same-day scans (which should not happen if acquisition settings exerted no impacts), and/or that such contributions were stronger. Error bars depict standard error of the mean across subjects.

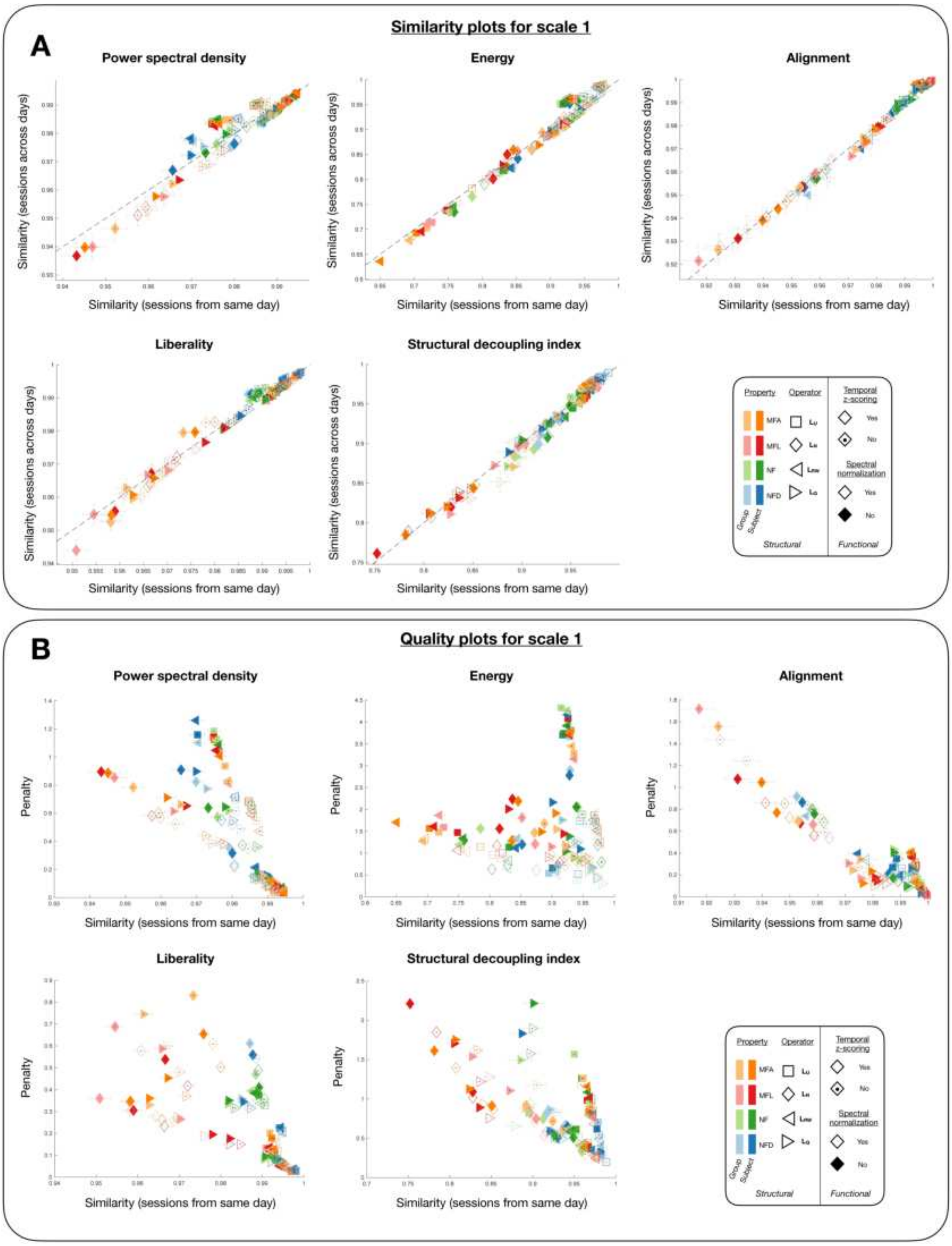


Figure 13: **(A)** On S1 input data, for the five feature types of interest, similarity between both scans acquired on the same day (X-axis) is contrasted to similarity between both scans acquired on different days, but with similar phase encoding direction (Y-axis). The similarity data was originally averaged across both available pairs of sessions in each case (*e.g.*, day-1 scans and day-2 scans). The error bars denote standard error of the mean across subjects. Each data point is associated to one specific set of parameters as described in the bottom right legend. **(B)** For the same five feature types, similarity between same-day scans (X-axis) is also contrasted to the devised penalty measure $\Omega$ (Y-axis), where a larger penalty value highlights that there were more subjects with larger similarity between different-day scans than between same-day scans (which should not happen if acquisition settings exerted no impacts), and/or that such contributions were stronger. Error bars depict standard error of the mean across subjects.

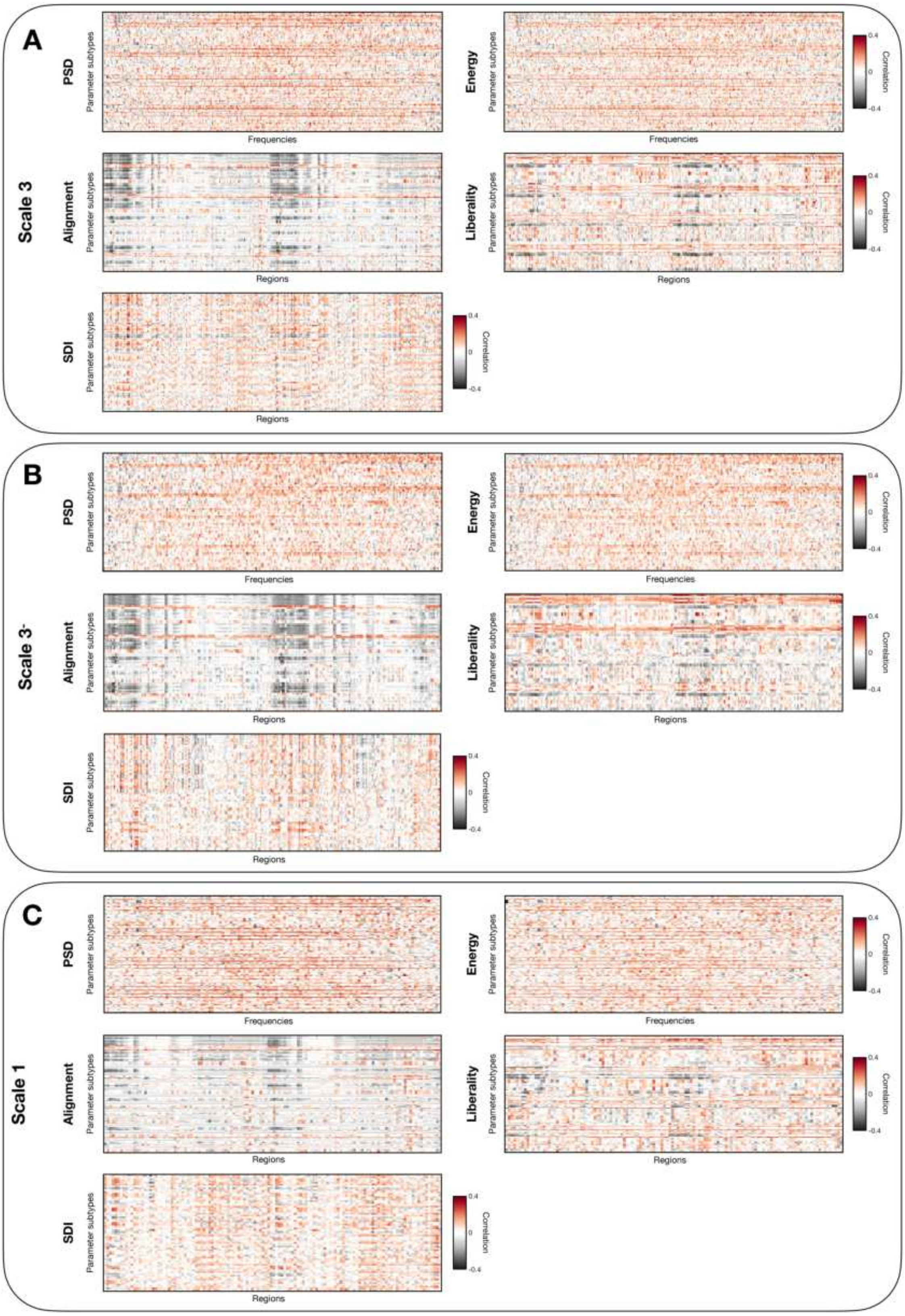


Figure 14: For S3 **(A)**, S3$^{-}$ **(B)** and S1 **(C)** input data, average Pearson's correlation between mean framewise displacement and feature coefficients across all four available scans. Each heatmap summarizes the results for one feature type, feature coefficients are shown from left to right, and parameter combinations from top to bottom.

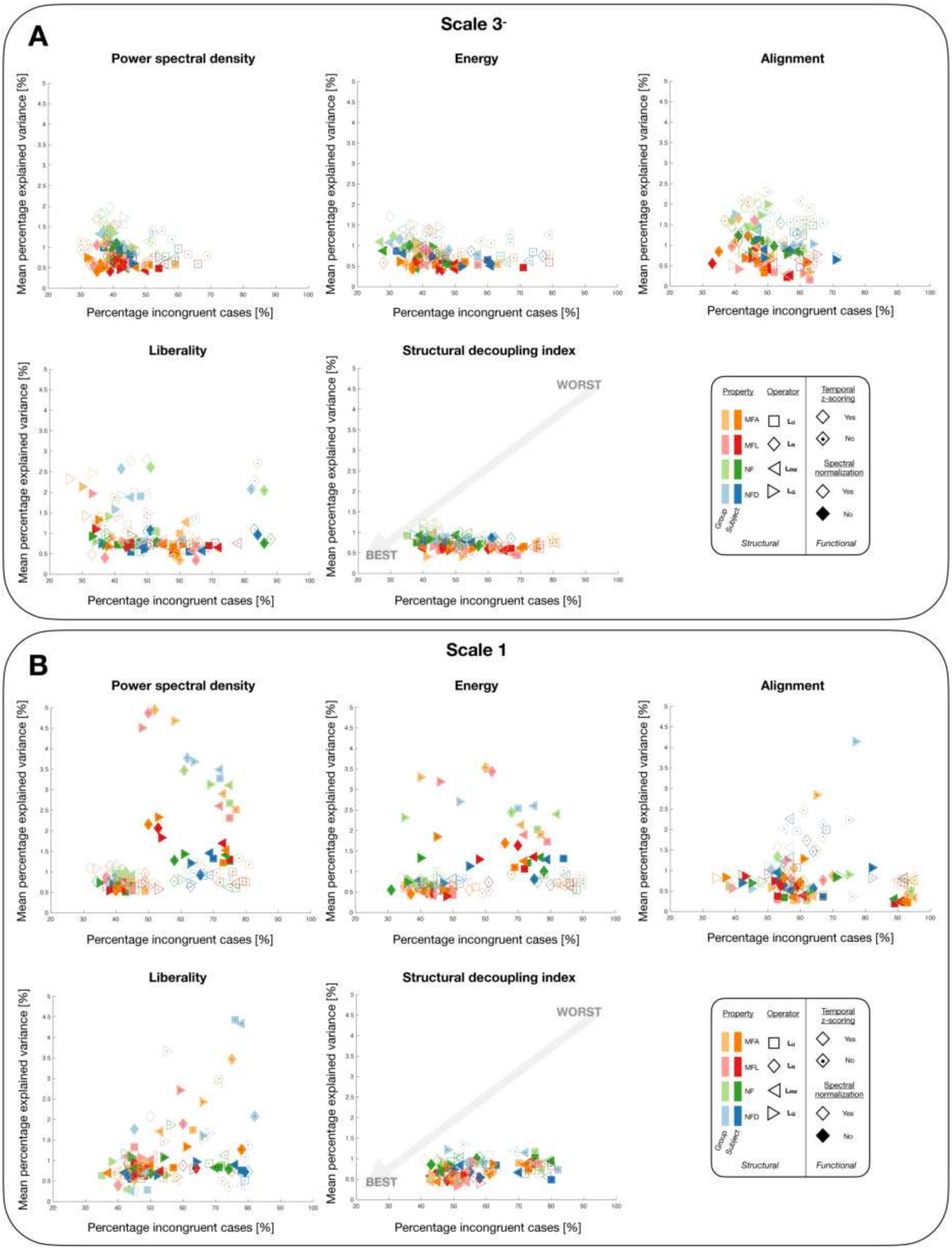


Figure 15: For S3- **(A)** and S1 **(B)** input data, summary displays of our two quality criteria regarding robustness. Each plot is linked to one feature type, and shows the percentage of incongruent subjects $P_I$ (*i.e.*, for whom same-day session similarity was lower than across-day session similarity), as well as the mean percentage of feature coefficient variance explained by mean framewise displacement (first averaged across coefficients, then across scans). The most robust outcomes are the ones for which both measures are the smallest (bottom left of the display). Each data point is associated to one specific set of parameters as described in the bottom right legend.

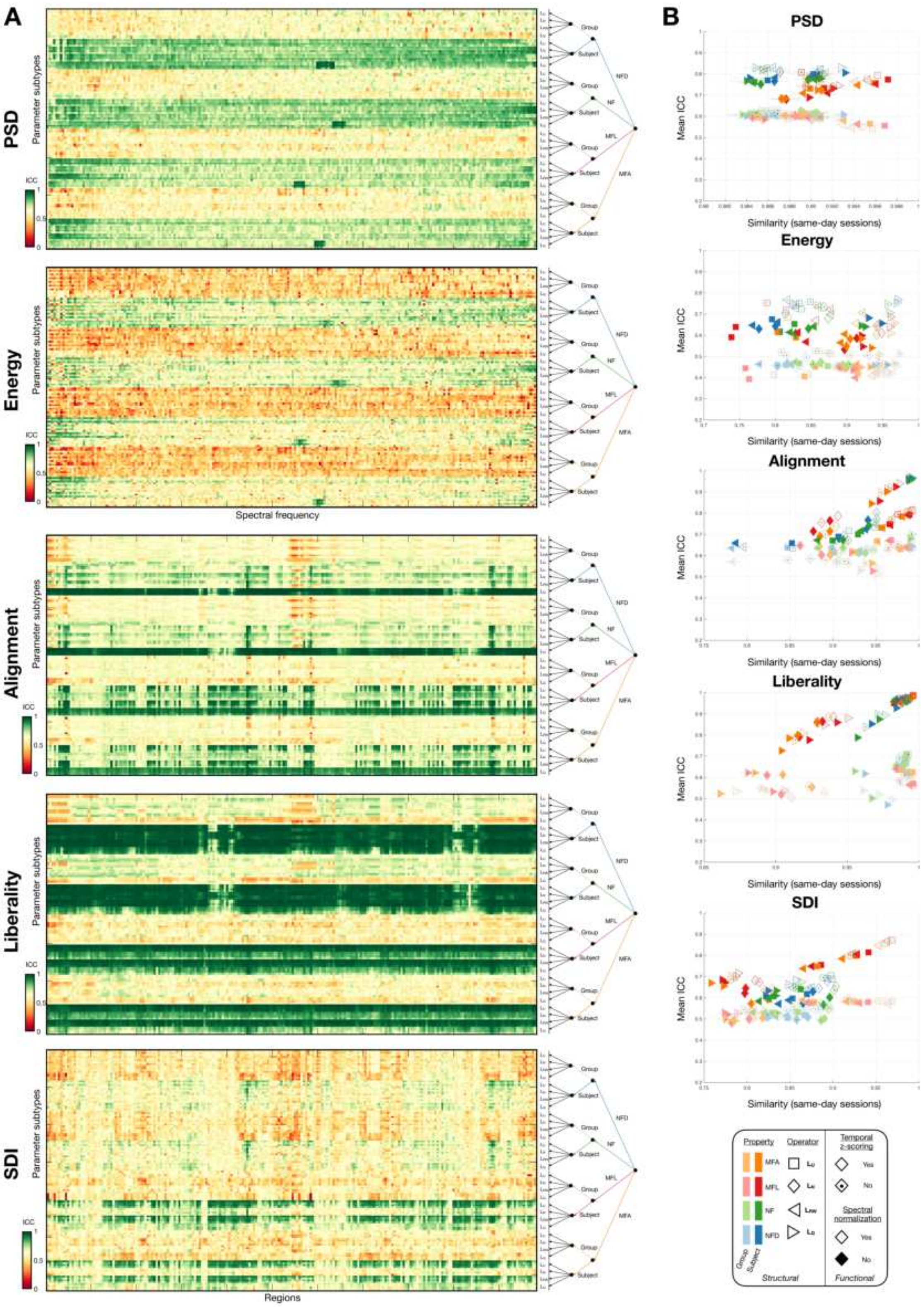


Figure 16: **(A)** For S3⁻ input data and the five feature types of interest, intra-class correlation (ICC) values are shown in heatmap representations across feature coefficients (from left to right) and parameter combinations (from top to bottom). On the right side of each heatmap, a tree representation summarizes the key parameter values associated to each row of the display. Note that for spectral domain features (PSD and energy), the X-axis denotes spectral frequency index, while for regional domain features (alignment, liberality and SDI), it instead represents region index. **(B)** For each feature type, same-day sessions' similarity (X-axis, a measure of robustness to acquisition settings, or intra-subject variability) is contrasted to mean ICC across feature coefficients (Y-axis, a measure of how much inter-subject variability outweighs intra-subject variability). The error bars denote standard error of the mean across subjects for similarity, and standard error of the mean across coefficients for ICC.

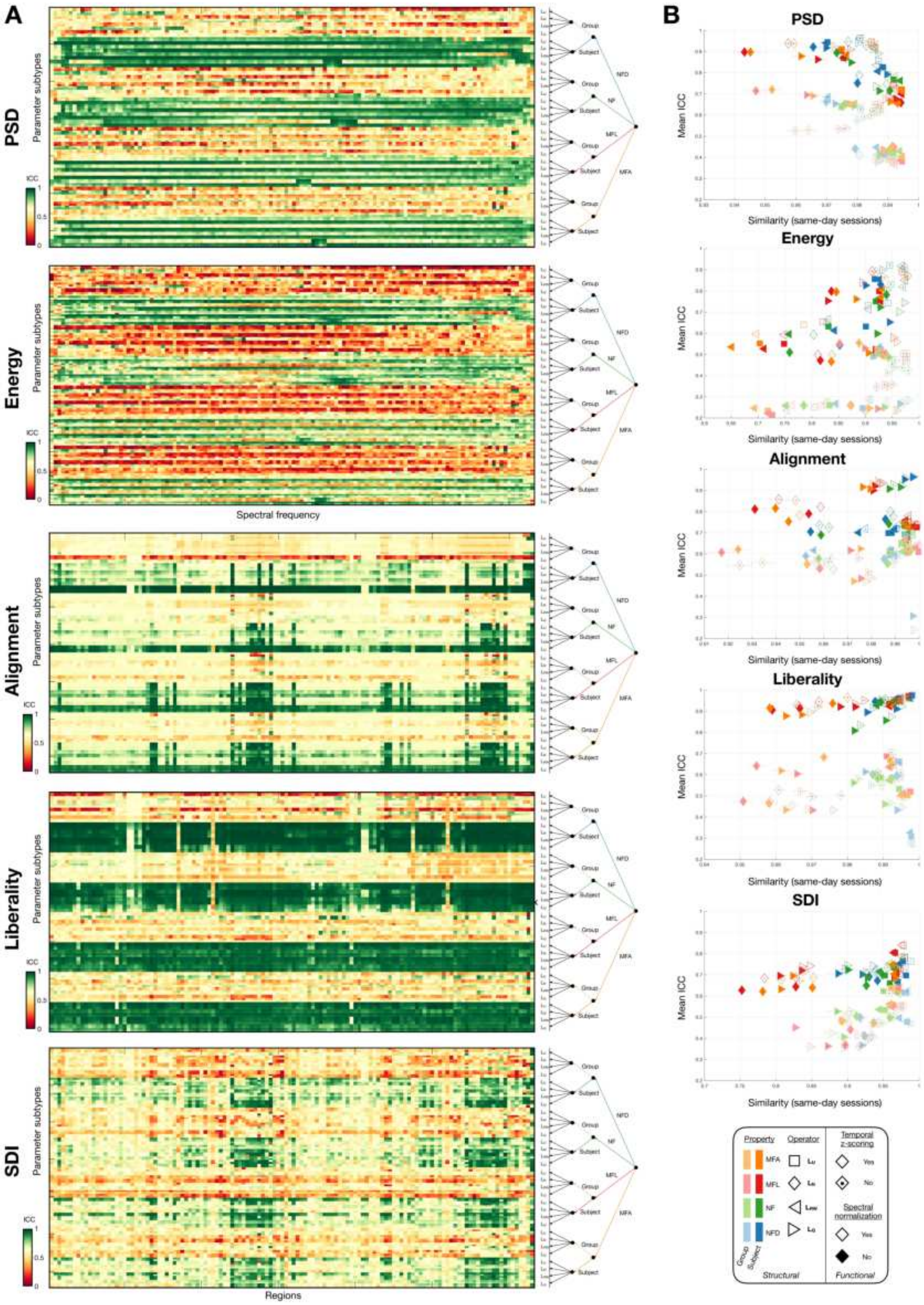


Figure 17: **(A)** For S1 input data and the five feature types of interest, intra-class correlation (ICC) values are shown in heatmap representations across feature coefficients (from left to right) and parameter combinations (from top to bottom). On the right side of each heatmap, a tree representation summarizes the key parameter values associated to each row of the display. Note that for spectral domain features (PSD and energy), the X-axis denotes spectral frequency index, while for regional domain features (alignment, liberality and SDI), it instead represents region index. **(B)** For each feature type, same-day sessions' similarity (X-axis, a measure of robustness to acquisition settings, or intra-subject variability) is contrasted to mean ICC across feature coefficients (Y-axis, a measure of how much inter-subject variability outweighs intra-subject variability). The error bars denote standard error of the mean across subjects for similarity, and standard error of the mean across coefficients for ICC.

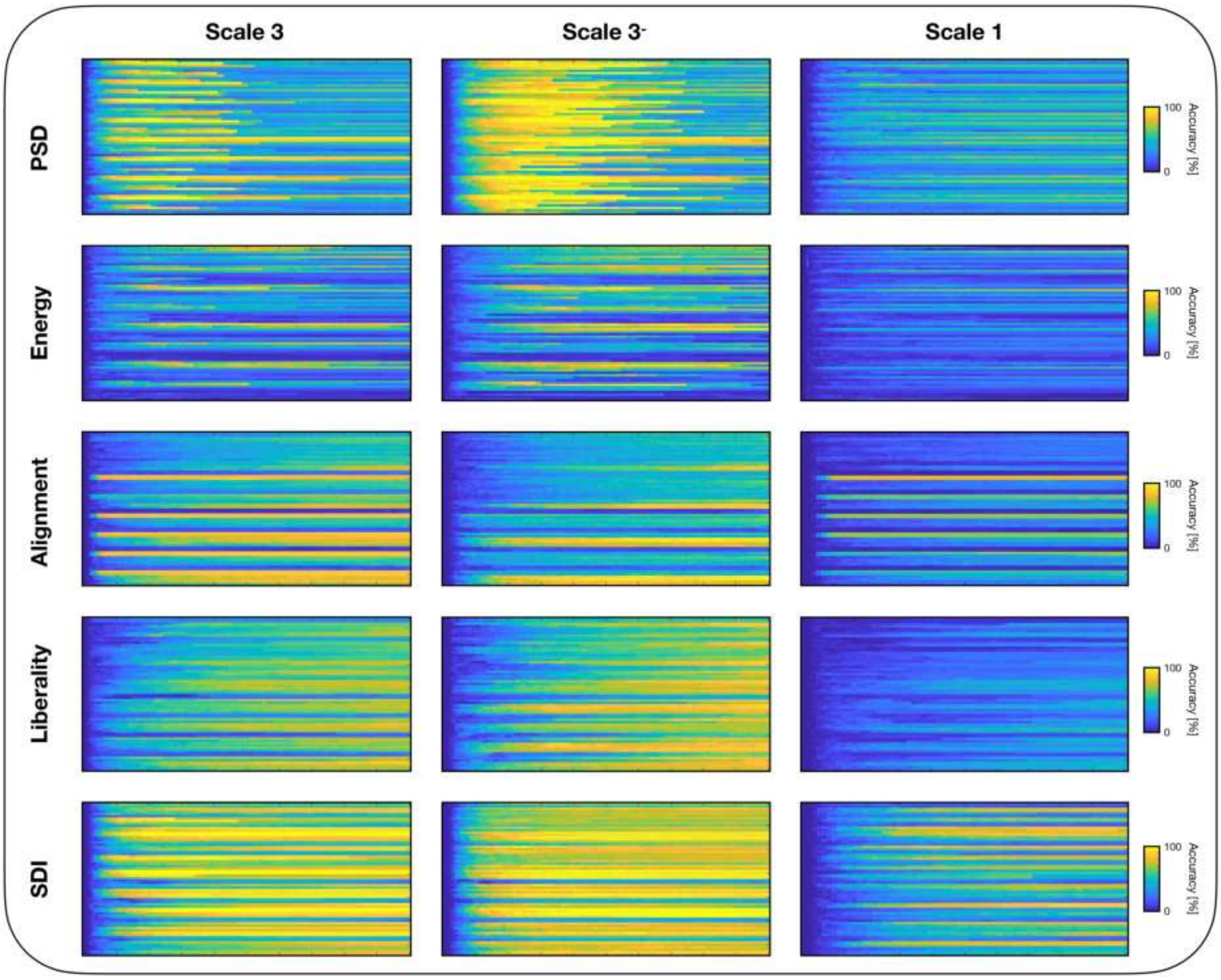


Figure 18: Fingerprinting accuracies are shown, for all feature types (rows) and S3 (left column), S3- (middle column) and S1 (right column) input data, for increasing percentages of used feature coefficients (from left to right in each heatmap display) and across parameter combinations (top to bottom). Features were gradually incorporated as a function of their ICC values, taken as a measure of cross-subject discriminability.

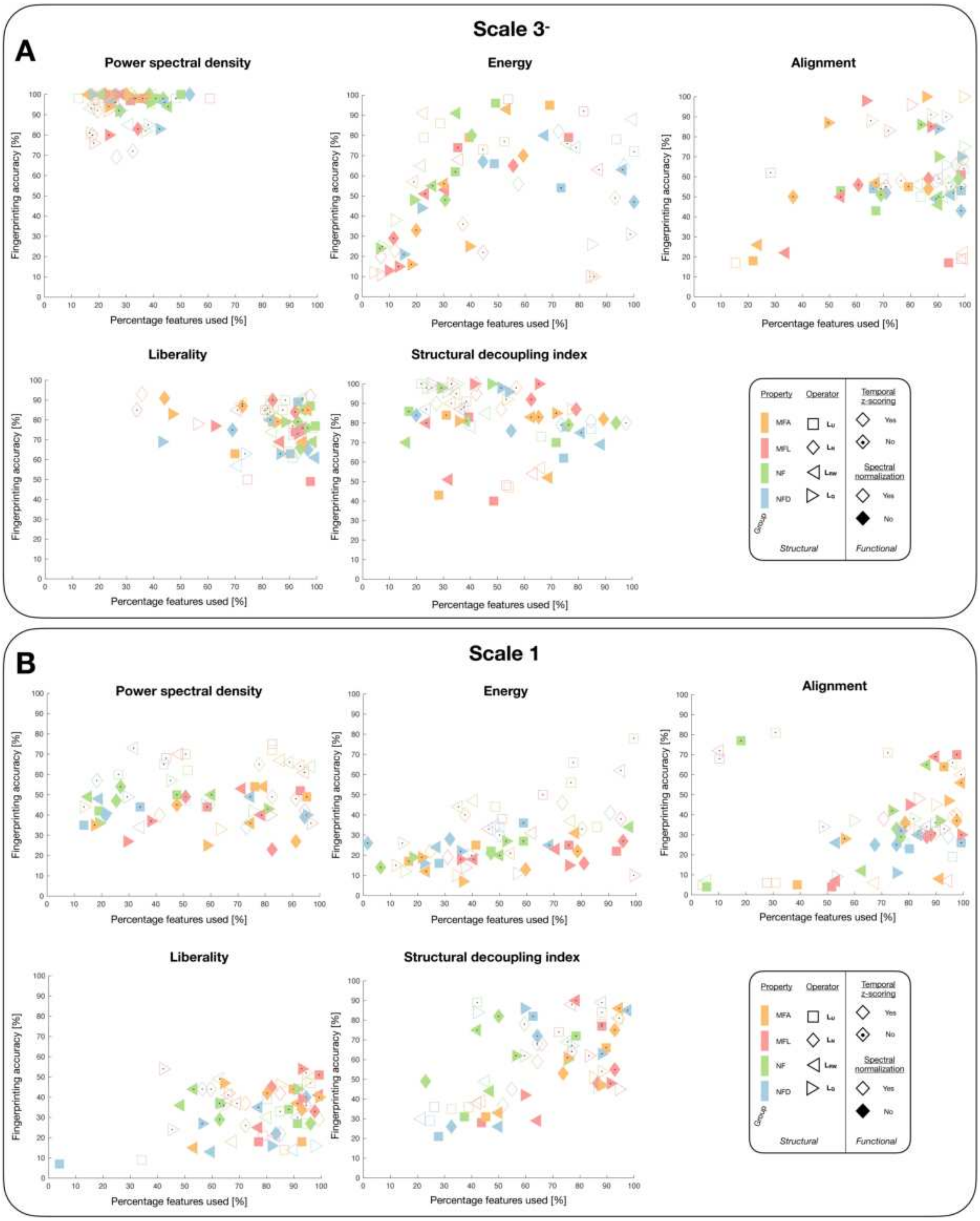


Figure 19: For S3- **(A)** and S1 **(B)** input data, for all five feature types of interest, the best achieved fingerprinting accuracies (Y-axis) are contrasted to the associated percentages of retained feature coefficients (X-axis). Each data point is associated to one specific set of parameters as described in the bottom right legend. Note that here, we focus on the 64 cases in which a group-wise SC was used, which is why there are only light-colored data points in the displays.

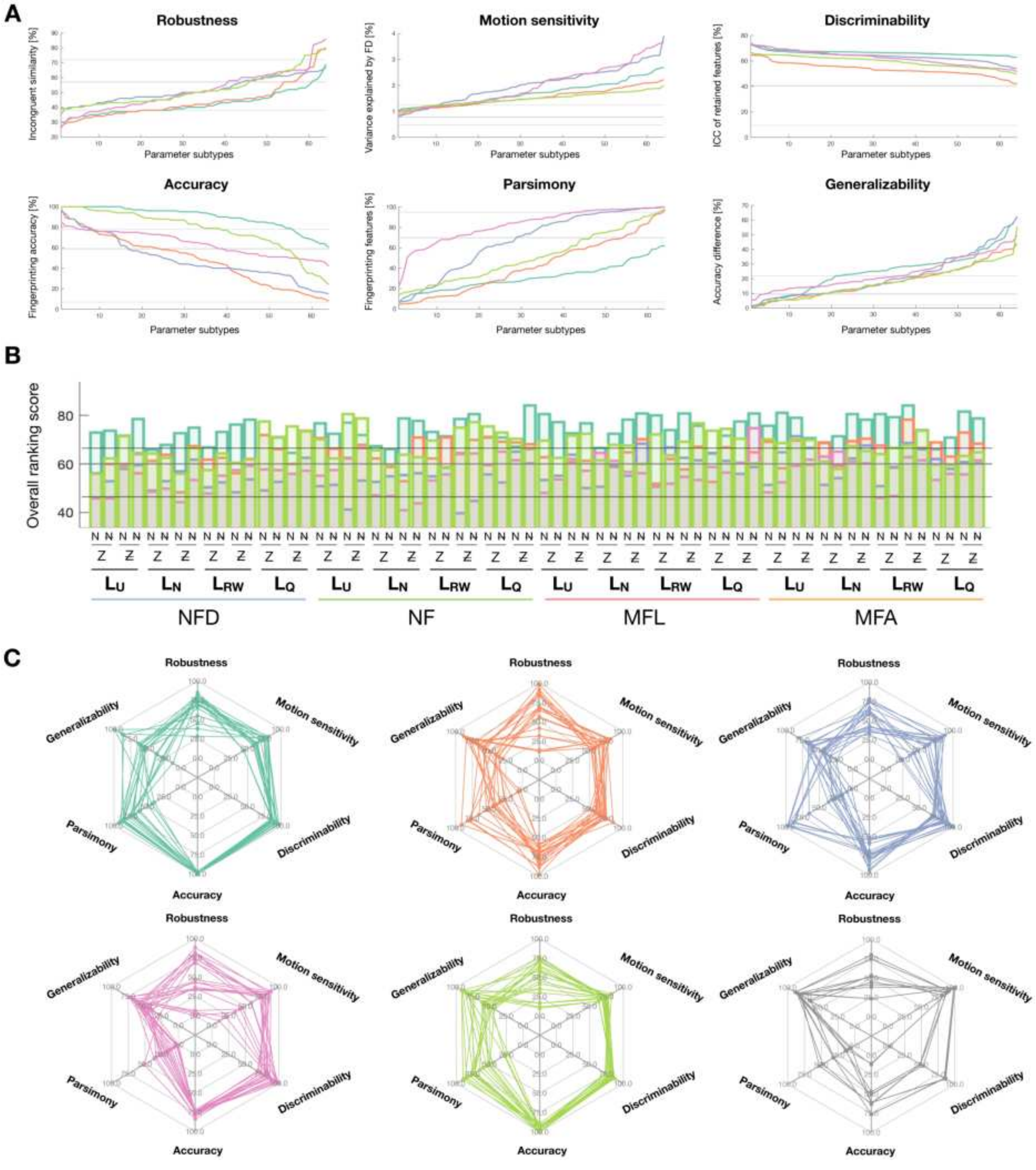


Figure 20: For S3⁻ input data, comparison of the six computed quality criteria across feature types **(A)**, where the data is ordered so that better values come first and the gray lines depict the minimum, median and maximum values achieved with graph metrics. Note that we focus on the 64 cases in which a group-wise SC was used. This is complemented by the overall ranking score across these 64 cases **(B)** for all feature types, where minimum, median and maximum scores for graph metrics are again shown as gray horizontal bars, and by spider plots summarizing the top 20 cases with maximal fingerprinting accuracy **(C)**.

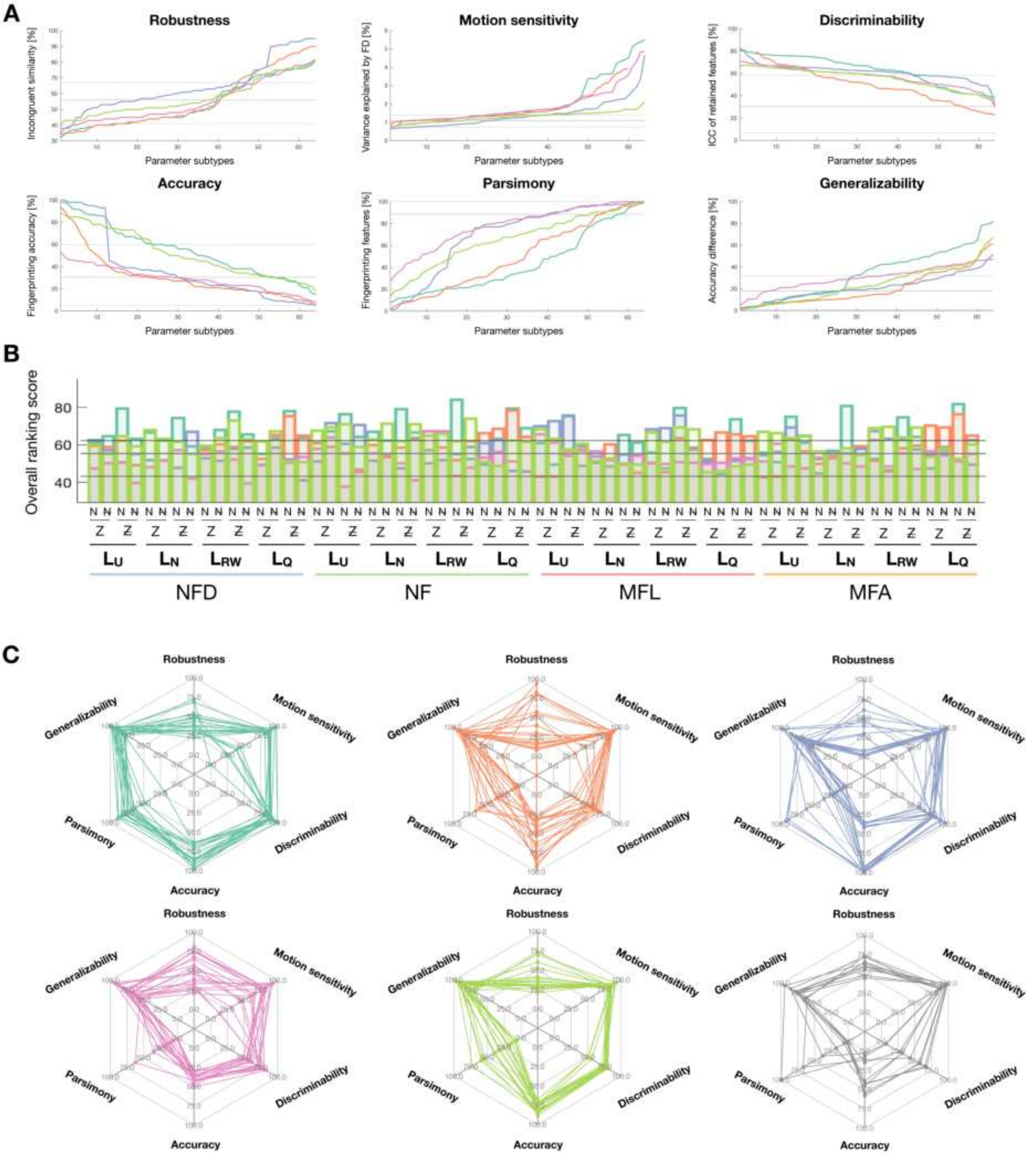


Figure 21: For S1 input data, comparison of the six computed quality criteria across feature types **(A)**, where the data is ordered so that better values come first and the gray lines depict the minimum, median and maximum values achieved with graph metrics. Note that we focus on the 64 cases in which a group-wise SC was used. This is complemented by the overall ranking score across these 64 cases **(B)** for all feature types, where minimum, median and maximum scores for graph metrics are again shown as gray horizontal bars, and by spider plots summarizing the top 20 cases with maximal fingerprinting accuracy **(C)**.

Table 1: For the scale 3 atlas without subcortex, summary of mixed-effects model results across feature types for all investigated fixed effects. The *Extra parameter* field stands for *Coefficients* for alignment/liberality, and *Cutoff frequency* for SDI. F-statistics are provided as mean $\pm$ standard deviation across coefficients, complemented by the minimum, median and maximum in brackets. The percentages of frequencies/regions for which significance was reached following Bonferroni correction are displayed in parentheses.

| | PSD | Energy | Alignment | Liberality | SDI |
|---|---|---|---|---|---|
| Intercept | $3302.3 \pm 1173.4$ (100%) $[239.31, 3466.7, 5419.1]$ | $405.2 \pm 182.62$ (100%) $[38.01, 400.78, 908.8]$ | $2786.3 \pm 1270.7$ (100%) $[111.75, 2987.5, 6008.6]$ | $4991.4 \pm 1761.2$ (100%) $[1440, 4853.3, 11168]$ | $13691 \pm 5053$ (100%) $[1362.9, 13899, 26422]$ |
| Property (P) | $193.63 \pm 142.34$ (98.15%) $[0.86, 163.71, 743.88]$ | $255.37 \pm 80.83$ (100%) $[41.73, 261.72, 488.08]$ | $50.12 \pm 64.17$ (89.81%) $[0.53, 32.64, 558.53]$ | $6288.3 \pm 5175.4$ (100%) $[162.78, 4772.5, 24608]$ | $9270.1 \pm 8008.1$ (100%) $[352.5, 6922.2, 36100]$ |
| Struct. indiv. (S) | n.a. | n.a. | $31.72 \pm 39.52$ (49.54%) $[1.5 \cdot 10^{(}-4), 15.81, 242.28]$ | $333.09 \pm 442.63$ (84.72%) $[0.05, 147.07, 2143.7]$ | $658.81 \pm 676.9$ (95.83%) $[0.23, 474.62, 4940.8]$ |
| Operator (O) | $429.43 \pm 297.28$ (99.54%%) $[5.03, 379.96, 1471.5]$ | $447.95 \pm 175.46$ (100%) $[48.66, 459.9, 829.42]$ | $217.74 \pm 276.88$ (93.98%) $[0.63, 114.67, 1705.9]$ | $21119 \pm 9226.4$ (100%) $[3895.1, 22062, 36531]$ | $17813 \pm 18315$ (100%) $[91.06, 11734, 1.19 \cdot 10^5]$ |
| Z-score (Z) | $38969 \pm 23033$ (100%) $[434.26, 35578, 10^5]$ | $603.87 \pm 218.34$ (100%) $[51.68, 630.4, 1224.7]$ | $14585 \pm 5902.1$ (100%) $[339.43, 16754, 24186]$ | $2.14 \cdot 10^5 \pm 71220$ (100%) $[1.0 \cdot 10^5, 2.01 \cdot 10^5, 4.8 \cdot 10^5]$ | $2490.9 \pm 3782.1$ (92.13%) $[0.29, 823.4, 25289]$ |
| Normalization (N) | $39050 \pm 23084$ (100%) $[435.11, 35643, 10^5]$ | $603.87 \pm 218.34$ (100%) $[51.68, 630.4, 1224.7]$ | $14713 \pm 5965.4$ (100%) $[341.45, 16912, 24425]$ | $2.33 \cdot 10^5 \pm 77645$ (100%) $[1.16 \cdot 10^5, 2.17 \cdot 10^5, 5.26 \cdot 10^5]$ | $806.91 \pm 684.8$ (99.07%) $[2.93, 626.9, 3666.9]$ |
| Extra parameter | n.a. | n.a. | $12092 \pm 4618.5$ (100%) $[304.85, 14042, 16220]$ | $3040.1 \pm 1224.2$ (100%) $[339.07, 2840.5, 6844]$ | $6948.5 \pm 4481$ (100%) $[486.51, 6179.5, 21767]$ |
| P * S | n.a. | n.a. | $25.69 \pm 34.9$ (76.39%) $[0.002, 17.36, 251]$ | $119.77 \pm 126.79$ (95.83%) $[1.07, 73.44, 648.05]$ | $323.85 \pm 301.98$ (98.15%) $[1.92, 230.06, 1792]$ |
| P * O | $209.49 \pm 82.78$ (99.54%) $[0.84, 208.04, 472.19]$ | $220.64 \pm 71.82$ (100%) $[41.02, 225.89, 390.91]$ | $54.44 \pm 48.75$ (98.61%) $[0.92, 40.14, 297.08]$ | $1135.5 \pm 823.62$ (100%) $[139.68, 876.89, 4337.6]$ | $5473.1 \pm 4526.8$ (100%) $[161.01, 3951.9, 18045]$ |
| S * O | n.a. | n.a. | $23.24 \pm 32.88$ (68.98%) $[0.013, 13.8, 241.18]$ | $242.33 \pm 154.89$ (98.61%) $[0.96, 215.48, 871.87]$ | $336.99 \pm 306.06$ (99.07%) $[4.35, 245.94, 2050]$ |
| Z * N | $38969 \pm 23033$ (100%) $[434.26, 35578, 10^5]$ | $603.87 \pm 218.34$ (100%) $[51.68, 630.4, 1224.7]$ | $14585 \pm 5902$ (100%) $[339.43, 16753, 24176]$ | $2.14 \cdot 10^5 \pm 71234$ (100%) $[1.08 \cdot 10^5, 2.01 \cdot 10^5, 4.8 \cdot 10^5]$ | $6.37 \pm 11.49$ (7.41%) $[8.02 \cdot 10^{(}-5), 2.89, 95.59]$ |
| P * S * O | n.a. | n.a. | $22.65 \pm 32.37$ (95.37%) $[0.002, 14.84, 248.73]$ | $72.52 \pm 43.93$ (100%) $[6.66, 64.27, 251.47]$ | $156.11 \pm 108.93$ (100%) $[7.48, 129.17, 537.59]$ |

Table 2: For the scale 1 atlas, summary of mixed-effects model results across feature types for all investigated fixed effects. The *Extra parameter* field stands for *Coefficients* for alignment/liberality, and *Cutoff frequency* for SDI. F-statistics are provided as mean ± standard deviation across coefficients, complemented by the minimum, median and maximum in brackets. The percentages of frequencies/regions for which significance was reached following Bonferroni correction are displayed in parentheses.

| | PSD | Energy | Alignment | Liberality | SDI |
|---|---|---|---|---|---|
| Intercept | 3736.4.3 ± 1040.3 (100%)<br>[375.4, 4020.3, 5484.4] | 650.8 ± 292.97 (99.21%)<br>[12.2, 669.78, 1406.8] | 3247.6 ± 1019.3 (100%)<br>[377.47, 3262.6, 5577.8] | 9679 ± 3649.9 (100%)<br>[1958.8, 9524.1, 19382] | 16605 ± 9456.3 (100%)<br>[1426.3, 13794, 52834] |
| Property (P) | 415.1 ± 264.31 (99.21%)<br>[1.23, 369.58, 1248.9] | 377.31 ± 119.67 (100%)<br>[16.19, 380.45, 790.71] | 214.04 ± 147.64 (99.21%)<br>[1.27, 183.57, 542.9] | 11593 ± 7208.3 (100%)<br>[2597.9, 9392.2, 34986] | 11771 ± 12419 (100%)<br>[85.14, 6979.6, 50383] |
| Struct. indiv. (S) | n.a. | n.a. | 62.75 ± 77.2 (68.25%)<br>[$7.27 \cdot 10^(-4)$, 33.55, 398.94] | 232.86 ± 315.8 (76.98%)<br>[0.17, 99.29, 1730.7] | 326.1 ± 410.03 (85.71%)<br>[0.02, 164.18, 2392.4] |
| Operator (O) | 906.76 ± 387.27 (100%%)<br>[111.99, 887.91, 1843.6] | 535.3 ± 189.87 (100%)<br>[16.32, 539.27, 925.84] | 664.08 ± 374.82 (100%)<br>[32.09, 629.1, 1614.5] | 15773 ± 9777.7 (100%)<br>[143.41, 15785, 33418] | 52967 ± 83273 (100%)<br>[125.3, 25080, $4.37 \cdot 10^5$] |
| Z-score (Z) | 10977 ± 6405.7 (100%)<br>[2131.3, 9846.3, 30461] | 789.63 ± 332.68 (100%)<br>[48.34, 756.35, 1944.1] | 8386.9 ± 2392.7 (100%)<br>[3884.3, 8452.2, 14804] | $1.83 \cdot 10^5 \pm 1.21 \cdot 10^5$ (100%)<br>[37698, $1.44 \cdot 10^5$, $8.11 \cdot 10^5$] | 21812 ± 17488 (98.41%)<br>[9.39, 19308, 70375] |
| Normalization (N) | 10984 ± 6409.6 (100%)<br>[2133.4, 9850.8, 30475] | 789.63 ± 332.68 (100%)<br>[48.34, 756.35, 1944.1] | 8402.8 ± 2400.4 (100%)<br>[3893.1, 8468.7, 14846] | $1.96 \cdot 10^5 \pm 1.32 \cdot 10^5$ (100%)<br>[38981, $1.55 \cdot 10^5$, $8.8 \cdot 10^5$] | 530.03 ± 691.61 (87.3%)<br>[0.13, 220.08, 3648.4] |
| Extra parameter | n.a. | n.a. | 7846.2 ± 2051.4 (100%)<br>[3745.5, 7871.7, 11886] | 2342.2 ± 975.31 (100%)<br>[623.4, 2215.5, 5506.5] | 4621.1 ± 3832.8 (100%)<br>[38.09, 3971.1, 15674] |
| P * S | n.a. | n.a. | 52.86 ± 54.49 (88.89%)<br>[0.005, 36.15, 365.95] | 102.1 ± 104.17 (95.24%)<br>[2.26, 70.21, 435.24] | 92.53 ± 95.18 (90.48%)<br>[1.27, 55.53, 433.62] |
| P * O | 330.2 ± 125.73 (99.21%)<br>[0.98, 322.37, 636.11] | 294.18 ± 86.44 (100%)<br>[20.24, 303.03, 439.98] | 161.11 ± 86.56 (99.21%)<br>[0.87, 156.96, 393.24] | 2709.8 ± 1516.9 (100%)<br>[293.19, 2422.7, 7691.1] | 6091.7 ± 6389.6 (100%)<br>[205.32, 3037.4, 27656] |
| S * O | n.a. | n.a. | 54.24 ± 49.14 (88.89%)<br>[0.042, 42.47, 265.37] | 210.22 ± 224.99 (99.21%)<br>[0.62, 132.98, 1002.9] | 147.07 ± 154.27 (97.62%)<br>[1.93, 99.36, 991.38] |
| Z * N | 10977 ± 6405.7 (100%)<br>[2131.3, 9846.3, 30461] | 789.63 ± 332.68 (100%)<br>[48.34, 756.35, 1944.1] | 8387 ± 2392.5 (100%)<br>[3884.5, 8452.5, 14797] | $1.83 \cdot 10^5 \pm 1.21 \cdot 10^5$ (100%)<br>[37677, $1.45 \cdot 10^5$, $8.14 \cdot 10^5$] | 29.72 ± 49.91 (42.06%)<br>[0.002, 9.5, 312.86] |
| P * S * O | n.a. | n.a. | 48.45 ± 40.68 (98.41%)<br>[0.0054, 33.16, 199] | 81.71 ± 77.22 (100%)<br>[7.97, 54.13, 358.07] | 82.81 ± 62.33 (100%)<br>[4.31, 69.99, 293.66] |
